# The Genetic Architecture of the Human Corpus Callosum and its Subregions

**DOI:** 10.1101/2024.07.22.603147

**Authors:** Ravi R. Bhatt, Shruti P. Gadewar, Ankush Shetty, Iyad Ba Gari, Elizabeth Haddad, Shayan Javid, Abhinaav Ramesh, Elnaz Nourollahimoghadam, Alyssa H. Zhu, Christiaan de Leeuw, Paul M. Thompson, Sarah E. Medland, Neda Jahanshad

## Abstract

The corpus callosum (CC) is the largest set of white matter fibers connecting the two hemispheres of the brain. In humans, it is essential for coordinating sensorimotor responses, performing associative/executive functions, and representing information in multiple dimensions. Understanding which genetic variants underpin corpus callosum morphometry, and their shared influence on cortical structure and susceptibility to neuropsychiatric disorders, can provide molecular insights into the CC’s role in mediating cortical development and its contribution to neuropsychiatric disease. To characterize the morphometry of the midsagittal corpus callosum, we developed a publicly available artificial intelligence based tool to extract, parcellate, and calculate its total and regional area and thickness. Using the UK Biobank (UKB) and the Adolescent Brain Cognitive Development study (ABCD), we extracted measures of midsagittal corpus callosum morphometry and performed a genome-wide association study (GWAS) meta-analysis of European participants (combined *N* = 46,685). We then examined evidence for generalization to the non-European participants of the UKB and ABCD cohorts (combined *N* = 7,040). Post-GWAS analyses implicate prenatal intracellular organization and cell growth patterns, and high heritability in regions of open chromatin, suggesting transcriptional activity regulation in early development. Results suggest programmed cell death mediated by the immune system drives the thinning of the posterior body and isthmus. Global and local genetic overlap, along with causal genetic liability, between the corpus callosum, cerebral cortex, and neuropsychiatric disorders such as attention-deficit/hyperactivity and bipolar disorders were identified. These results provide insight into variability of corpus callosum development, its genetic influence on the cerebral cortex, and biological mechanisms related to neuropsychiatric dysfunction.

## Introduction

The corpus callosum (CC) is the largest white matter tract in the human brain, facilitating higher order functions of the cerebral cortex by allowing the two hemispheres of the brain to communicate^1,2^. This connection is essential for coordinating sensorimotor responses, performing associative and executive functions, and representing information in multiple dimensions^3,4^. Most CC fibers connect corresponding left and right cortical regions of the brain, with the organization, development of axonal elongation, and myelination of callosal fibers being correlated with the rostro-caudal (front-to-back) distribution of functional areas^5,6^. Regional alterations in CC shape are easily assessed with neuroimaging studies, which have found local callosal abnormalities in complex neurodevelopmental and neuropsychiatric disorders^6–11^, such as lower anterior volumes in autism^12^ and lower posterior thickness in bipolar disorder^13^. Twin studies show up to 66% heritability for CC area^14,15^, and previous single-cohort studies of genetic influences on CC volume and its relationship to neuropsychiatric disorders have found heritability estimates between 22-39%^16,17^. Yet, the interplay between genetic variants influencing CC morphometry, the cerebral cortex, and associated neuropsychiatric disorders is not well understood.

3D magnetic resonance imaging (MRI) provides a non-invasive approach to quantify individual variations in brain regions and connections^6^, including the morphology of the CC, and how they are associated with brain-based traits and diseases. The midsagittal section of an anatomical brain MRI scan is able to capture the entire rostro-caudal formation of the CC, which is almost always in the field of view of 2D clinical and 3D research MRI scans alike. This 2D midsagittal representation can be segmented to offer a lower dimensional projection of the anatomical intricacies of the CC, allowing for structural measures of CC area and thickness to be computed^18,19^. We developed and validated a fully automated artificial intelligence based CC feature extraction tool, *Segment, Measure, and AutoQC the midsagittal CC* (*SMACC)*, which we make publicly available at https://github.com/USC-LoBeS/smacc^20^.

Using data from the UK Biobank^21^ (UKB) and Adolescent Brain Cognitive Development^22^ (ABCD) studies, here we present results from a genome-wide association study (GWAS) meta-analysis of total area and mean thickness of the CC derived using *SMACC.* We also present the results for five differentiated areas based on distinguishable projections to (1) prefrontal, premotor and supplementary motor, (2) motor, (3) somatosensory, (4) posterior parietal and superior temporal, and (5) inferior temporal and occipital cortical brain regions^23,24^. These regions are believed to represent structural-functional coherence^6^. We performed a GWAS meta-analysis using two population-based cohorts, one of adolescents and another of older adults, to examine genetic influences on CC area and thickness^25,26^. The primary analyses were in individuals of European ancestry and the same analyses were then repeated using the data from non-European participants to assess consistency in the magnitude and direction of effect sizes. Downstream post-GWAS analyses investigated the enrichment of genetic association signals in tissue types, cell types, brain regions, and biological pathways. We examined the genetic overlap at the global and local level, using LD Score regression (LDSC)^27^ and Local Analysis of Variant Association (LAVA)^28^, respectively, and the causal genetic relationships between CC phenotypes, cortical morphometry, and related neuropsychiatric conditions.

## Results

### Characterization of corpus callosum shape associated loci

We conducted a GWAS of area and mean thickness of the whole corpus callosum, and five regions of the Witelson parcellation scheme (**Fig. 1**)^23,24^, using data from participants of European ancestry from the UKB (*N* = 41,979) and ABCD cohorts (*N* = 4,706). A meta-analysis of GWAS summary statistics of all CC derived metrics in UKB and ABCD was performed using METAL and the random-metal extension^29,30^, based on the DerSimonian-Laird random-effects model (**Methods**). To examine the generalizability of single nucleotide polymorphism (SNP) effects across ancestries, these same analyses were run using data from non-European participants (total *N* = 7,040).

**Figure 1:**
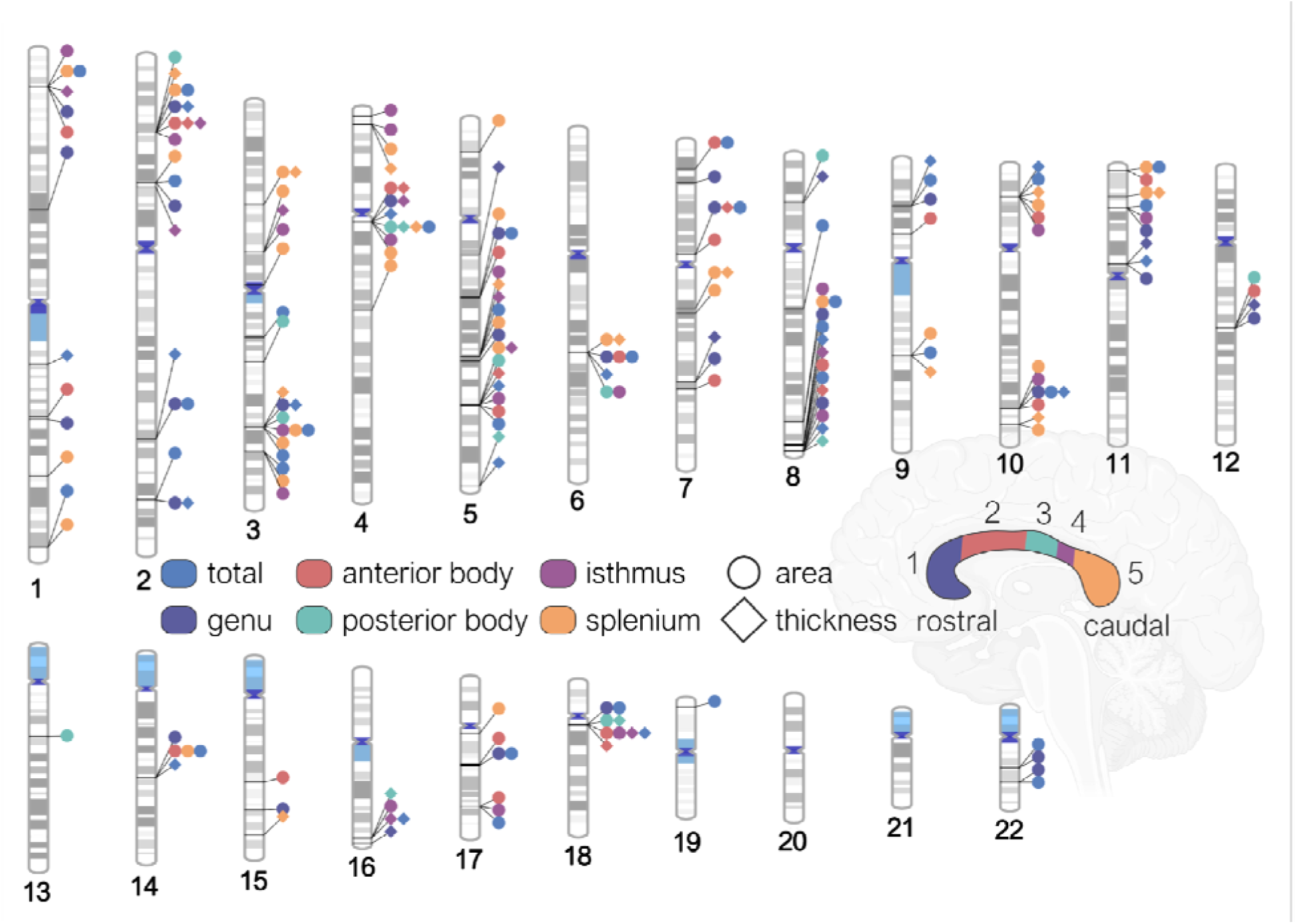
Regions of the midsagittal corpus callosum and associated genomic loci. An ideogram representing loci that influence total corpus callosum area, its mean thickness, and area and thickness of individual parcellations determined by the Witelson parcellation scheme in a rostral-caudal gradient (1-5). All loci are significant at the Bonferroni corrected, experiment-wide threshold of p < 6.13 × 10-9.

The GWAS meta-analysis identified 48 independent significant SNPs for total area and 18 independent SNPs for total mean thickness. Independent significant SNPs were determined in FUMA using the default threshold of *r*^2^ = 0.6, and genomic loci were determined at *r*^2^ = 0.1. This identified 28 genomic loci for total cross-sectional area, and 11 genomic loci for total mean thickness. All significant loci for total area and mean thickness showed concordance in the direction of effect between the two cohorts. There were 5 loci, all in intronic regions, each positionally mapped to genes^31^ that overlapped between area and mean thickness. These included *IQCJ-SHIP1* (multimolecular complexes of initial axon segments and nodes of Ranvier, and calcium mediated responses)^32^, *FIP1L1* (RNA binding and protein kinase activity)^33^, *HBEGF* (growth factor activity and epidermal growth factor receptor binding)^34^, *CDKN2B-AS1* (involved in the NF-κB signaling pathway with diverse roles in the nervous system)^35,36^, and *FAM107B* (cytoskeletal reorganization in neural cells and cell migration/expansion)^37^. The genomic locus mapped to *IQCJ-SHIP1* had a positive effect for total area (rs11717303, effect allele: C, effect allele frequency (EAF): 0.689, β = 4.28, s.e. = 0.51, *p* = 4.54 × 10^−17^). The same locus showed a negative effect for a different SNP on total thickness (rs12632564, effect allele: T, EAF: 0.305, β = −0.042, s.e. = 0.006, *p* = 2.59 × 10^−12^). The strongest locus for total area (rs7561572, effect allele: A, EAF: 0.532, β = −4.13, s.e. = 0.46, *p* = 1.98 × 10^−18^) was positionally mapped to the *STRN* gene. The strongest locus for mean thickness (rs4150211, effect allele: A, EAF: 0.265, β = −0.05, s.e. = 0.006, *p* = 8.20 × 10^−18^) was mapped to the *HBEGF* gene.

Loci for area overlapped between parcellations in a rostral-caudal gradient (1-5), such that: rs1122688 on the *SHTN1* (or *KIAA1598*) gene (involved in positive regulation of neuron migration) overlapped between the genu (1) and anterior body (2); rs1268163 near the *FOXO3* gene (involved in IL-9 signaling and *FOXO*-mediated transcription) overlapped between the posterior body (3) and isthmus (4); and rs11717303 on the *IQCJ-SCHIP1* gene overlapped between the isthmus (4) and splenium (5). This gradient pattern was not observed for mean thickness. The strongest regional association was observed with splenium area (rs10901814, effect allele: C, EAF: 0.584, β = −1.69, s.e. = 0.16 *p* = 2.02 × 10^−24^) and thickness (rs11245344, effect allele: T, EAF: 0.570, β = −0.11, s.e. = 0.11, *p* = 6.28 × 10^−22^), both on the *FAM53B* gene. *FAM53B* is involved in positive regulation of the canonical Wnt signaling pathway. We observed a concordance in direction and similar magnitude effect sizes in the analyses within the data from the non-European participants. Detailed annotations and regional association plots of all genomic loci, independent significant SNPs and genes are in **Supplementary Tables S1-S4 and Extended Data 1**.

### SNP heritability and genetic correlation between cohorts

Moderate to high genetic correlations were seen across CC phenotypes between cohorts, with *r_g_*ranging from 0.54 (*s.e.* = 0.27) and 0.92 (*s.e.* = 0.63) for area metrics, and 0.30 (*s.e.* = 0.16) and 0.99 (*s.e.* = 0.69) for thickness metrics. We used the GREML approach implemented in GCTA^38,39^ to estimate SNP heritability (*h^2^_SNP_*) for each cohort. Within the UKB, heritability values ranged for different CC phenotypes from 0.42 - 0.71, with similar results seen in the ABCD cohort (**Supplementary Tables S5-S8)**. Total area (UKB *h^2^_SNP_* = 0.71, *s.e.* = 0.01; ABCD *h^2^_SNP_* = 0.74, *s.e.* = 0.03) and mean thickness (UKB *h^2^_SNP_* = 0.60, *s.e.* = 0.02; ABCD *h^2^_SNP_* = 0.77, *s.e.* = 0.03) showed the highest *h^2^_SNP_* across both cohorts. LDSC^27^ *h^2^_SNP_* estimates from the meta-analysis ranged between 0.10 (*s.e.* = 0.01) and 0.18 (*s.e.* = 0.05) for area, and 0.12 (*s.e.* = 0.01) and 0.16 (*s.e.* = 0.02) for thickness, with the area of the genu showing the highest, and area of the splenium showing the lowest *h^2^_SNP_*estimates. As shown in **Supplementary Tables S5-S8, all** LDSC R_G_ estimates between meta-analyzed CC phenotypes were significant.

### Gene-mapping and gene-set enrichment analyses

Gene-based association analysis in MAGMA^40^ identified 30 genes for the total area, and 34 genes for total mean thickness of the CC, with 5 genes overlapping between area and thickness (*IQCJ-SCHIP1, IQCJ, BPTF, PADI2, CHIC2*). The strongest association seen with area was *AC007382.1* and the strongest association with mean thickness was *HBEGF* (**Fig. 2A**). There were between 15 and 31 genes for area, and between 7 and 25 genes for thickness identified within regions of the CC. Notably, *IQCJ*, *IQCJ-SCHIP1*, and *STRN* overlapped for all parcellations of CC area. *AC007382.1* overlapped for four out of five parcellations, and *STRN* and *PARP10* overlapped for three out of five parcellations of CC thickness (**Fig. 2B**, **Supplementary Tables S1-S4**). Enrichment of SNP heritability in 53 functional categories for each trait was determined via LDSC^41^. The majority of enrichment and the strongest effects across parcellations of the CC were observed in categories related to gene regulation/transcription in chromatin (**Fig. 3A-B**).

**Figure 2:**
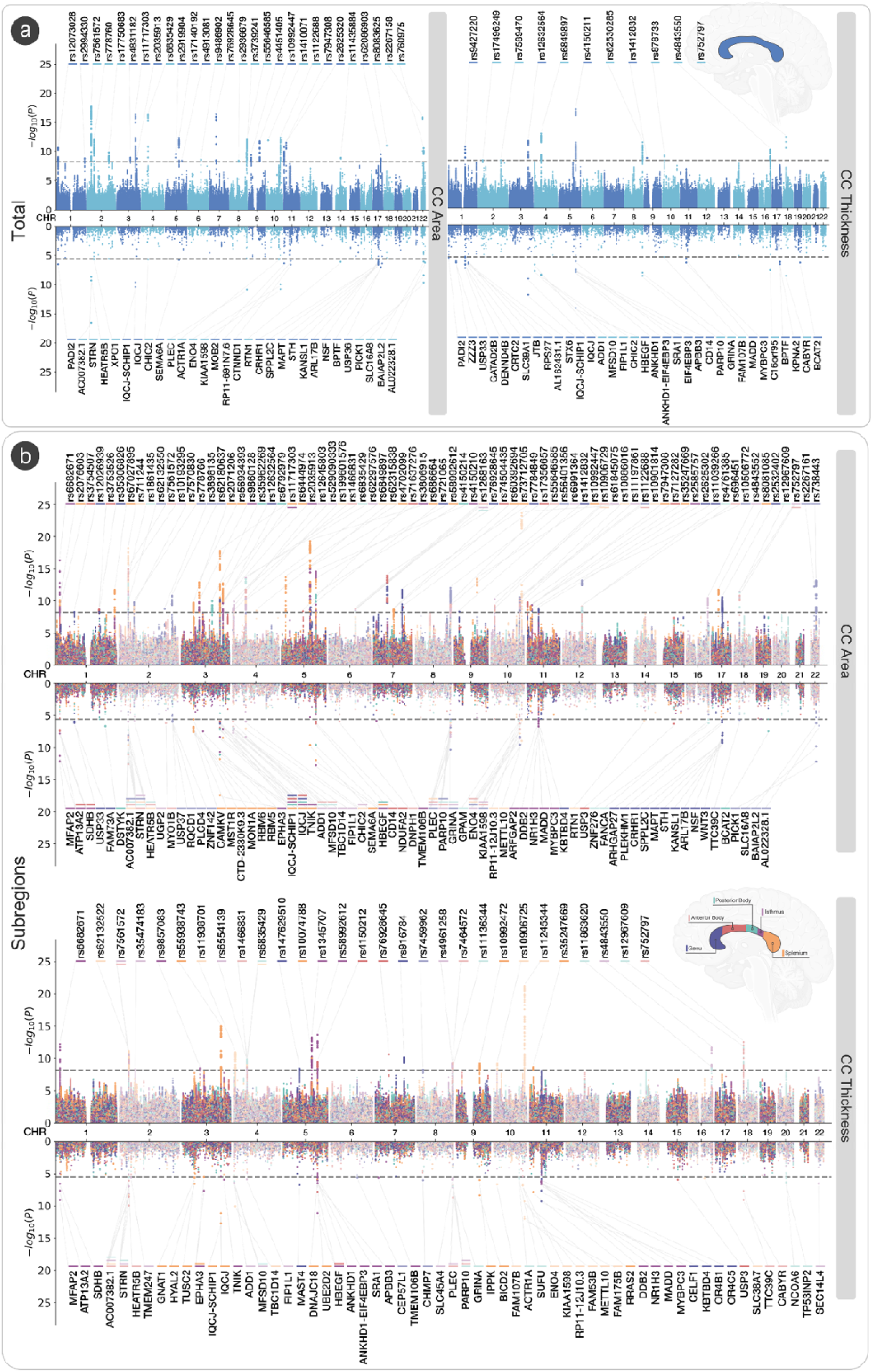
GWAS meta-analysis of midsagittal corpus callosum area and thickness. (A) Miami plot for SNPs (*top*) and genes (*bottom*) based on MAGMA gene analysis for total area and total mean thickness. (B) Miami plot for SNPs (*top*) and genes (*bottom*) based on MAGMA gene analysis for area of thickness of the CC split by the Witelson parcellation scheme^23^. Significant SNPs and genes are color coded by corpus callosum traits.

**Figure 3:**
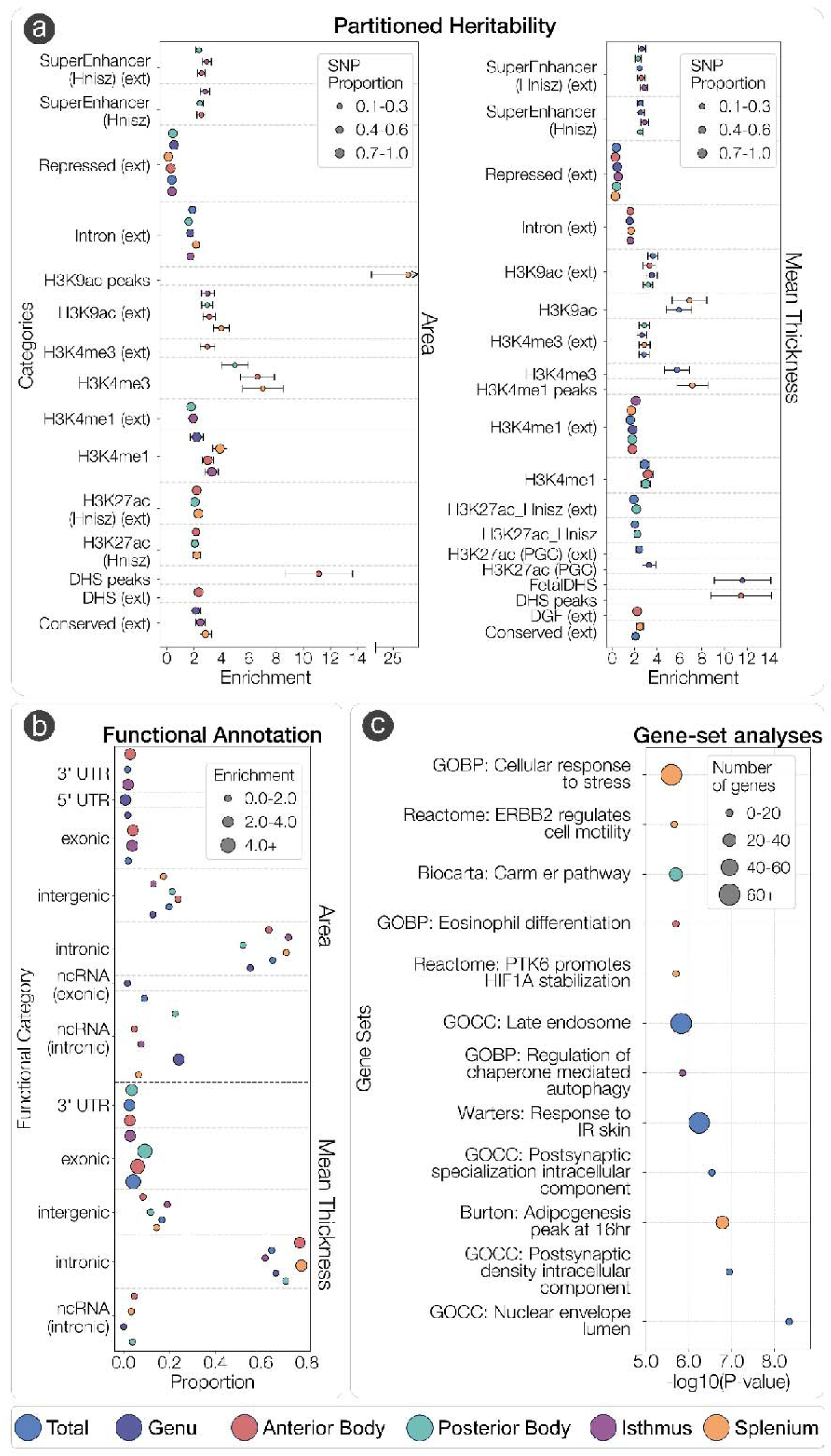
Partitioned heritability, functional annotation and enrichment of gene-sets of CC morpholog associated genetic variants. (A) Significant enrichment of SNP heritability across 53 functional categories compute by LD Score regression for area (*left*) and mean thickness (*right*). Error bars indicate 9% confidence intervals. (B) Proportion of GWAS SNPs in each functional category from ANNOVAR across each CC phenotype. (C) Significant gene-sets across CC phenotypes computed via MAGMA gene-set analysis at the Bonferroni corrected threshold of 3.23 × 10^-6^. GOBP: Gene-ontology biological processes, GOCC: Gene-Ontology Cellular Components.

Gene-set enrichment analyses were also completed in MAGMA (**Fig. 3C**). Strongest effects of significant gene sets included those involved in postsynaptic specialization for total CC area, including GO:009901 (postsynaptic specialization, intracellular component) and GO:009902 (postsynaptic density, intracellular component). A theme of signal transduction related pathways was observed for splenium area including R-HSA-6785631 (*ERBB2 regulates cell motility*) and R-HSA-8857538 (*PTK6 promotes HIF1A stabilization*). Enrichment of the “*CARM1 and regulation of the estrogen receptor*” was found for the posterior body thickness and is implicated transcriptional regulation via histone modifications. Enrichment of GO:1904714 (*regulation of chaperone-mediated autophagy*) was found for the isthmus area, which is implicated in lysosomal-mediated protein degradation. All significant results across all CC phenotypes are in **Supplementary Table 18**.

### Tissue-specific and cell-type specific expression of corpus callosum associated genes

Gene-property enrichment analyses were completed in MAGMA with 54 tissue types from GTEx v8 and BrainSpan^42,43^, which includes 29 samples from individuals representing 29 different ages, as well as 11 general developmental stages. An enrichment of genes associated with isthmus thickness were expressed in the cerebellum (*p_(Bon)_* = 0.017). Area and thickness across parcellations of the CC showed an enrichment of expression of genes in the brain from early prenatal to late mid-prenatal developmental stages. An enrichment of expression of genes associated with area and thickness of the anterior body of the CC was observed in brain tissue prenatally 9 to 24 weeks post conception. Enrichment of expression of genes associated with area of the genu was observed in brain tissue 19 weeks post conception. Enrichment of expression of genes associated with total mean thickness of the CC was observed in brain tissue 19 weeks post conception. All results are shown in **Supplementary Tables S19-S21**. These results, along with the gene-sets involved in histone modifications, were supported by LDSC-SEG analyses using chromatin-based annotations from narrow peaks^44^, which showed a significant enrichment in the heritability by variants located in genes specifically expressed in DNase in the female fetal brain for total CC thickness (*p_(Bon)_* = 0.0105). Chromatin annotations showed a consistent and significant enrichment of splenium area and thickness associated variants in histone marks of the fetal brain and neurospheres (**Supplementary Table S25**).

Using microarray data from 292 immune cell types, area of the posterior body showed a significant enrichment in the heritability by variants located in genes specifically expressed in multiple types of myeloid cells (*p_(Bon)_*< 0.05), and area of the isthmus showed enrichment in innate lymphocytes (*p_(Bon)_* = 0.047). This further validates the aforementioned significant locus on gene *FOXO3,* which overlapped between the posterior body and isthmus (**Supplementary Table S26**).

Cell-type specific analyses were performed in FUMA using data from 13 single-cell RNA sequencing datasets from the human brain. This tests the relationship between cell-specific gene expression profiles and phenotype-gene associations^45^. Of the 12 phenotypes tested, only total CC thickness showed significant results after going through the 3-step process using conditional analyses to avoid bias from batch effects from multiple scRNA-seq datasets. The most significant association was seen with oligodendrocytes located in the middle temporal gyrus (MTG, *p_(Bon)_* = 0.001) from the Allen Human Brain Atlas (AHBA). Oligodendrocytes (*p_(Bon)_* = 0.03) and non-neuronal cells (*p_(Bon)_* = 0.03) located in the lateral geniculate nucleus (LGN) from the AHBA also showed significant associations but were collinear (**Supplementary Table S22**).

LAVA-TWAS analyses^28,46^ (**Fig. 4**) of expression quantitative trait loci (eQTLs) and splicing quantitative trait loci (sQTLs) of protein-coding genes in 16 different brain, cell type, and whole blood tissues revealed the strongest eQTL associations of area and thickness with *CROCC* expression in whole blood for the isthmus (ρ = −0.53, *p* = 1.29 × 10^−10^). Other notable eQTL (**Supplementary Table S29**) findings included total CC area and isthmus area and thickness being positively associated with *ATP13A2* expression in fibroblasts (ρ = 0.48, *p* = 1.58 × 10^−7^). The strongest sQTL association was a positive association observed with *KANSL1* cluster 11710 in fibroblasts for genu area (ρ = 0.83, *p* = 1.46 × 10^−14^), which was the tissue type where most observed associations occurred across CC phenotypes (**Supplementary Table S30**). Moreover, a negative association was observed in a *KANSL1* (cluster 11707) in fibroblasts for the genu area (ρ = 0.82, *p* = 3.11 × 10^−7^). An sQTL in *MFSD13A* (cluster 7894) in the anterior cingulate showed very strong yet opposite associations for total CC thickness (ρ = 0.42, *p* = 1.12 × 10^−13^) and total CC area (ρ = −0.44, *p* = 2.98 × 10^−11^). Other notable findings across tissue types included *CRHR1* in the cortex, nucleus accumbens, and putamen, as well as *UGP2* in fibroblasts, whole blood, and the putamen. No significant results from LAVA-TWAS gene-set enrichment analyses were observed after Bonferroni correction (**Supplementary Tables S31-S32**).

**Figure 4:**
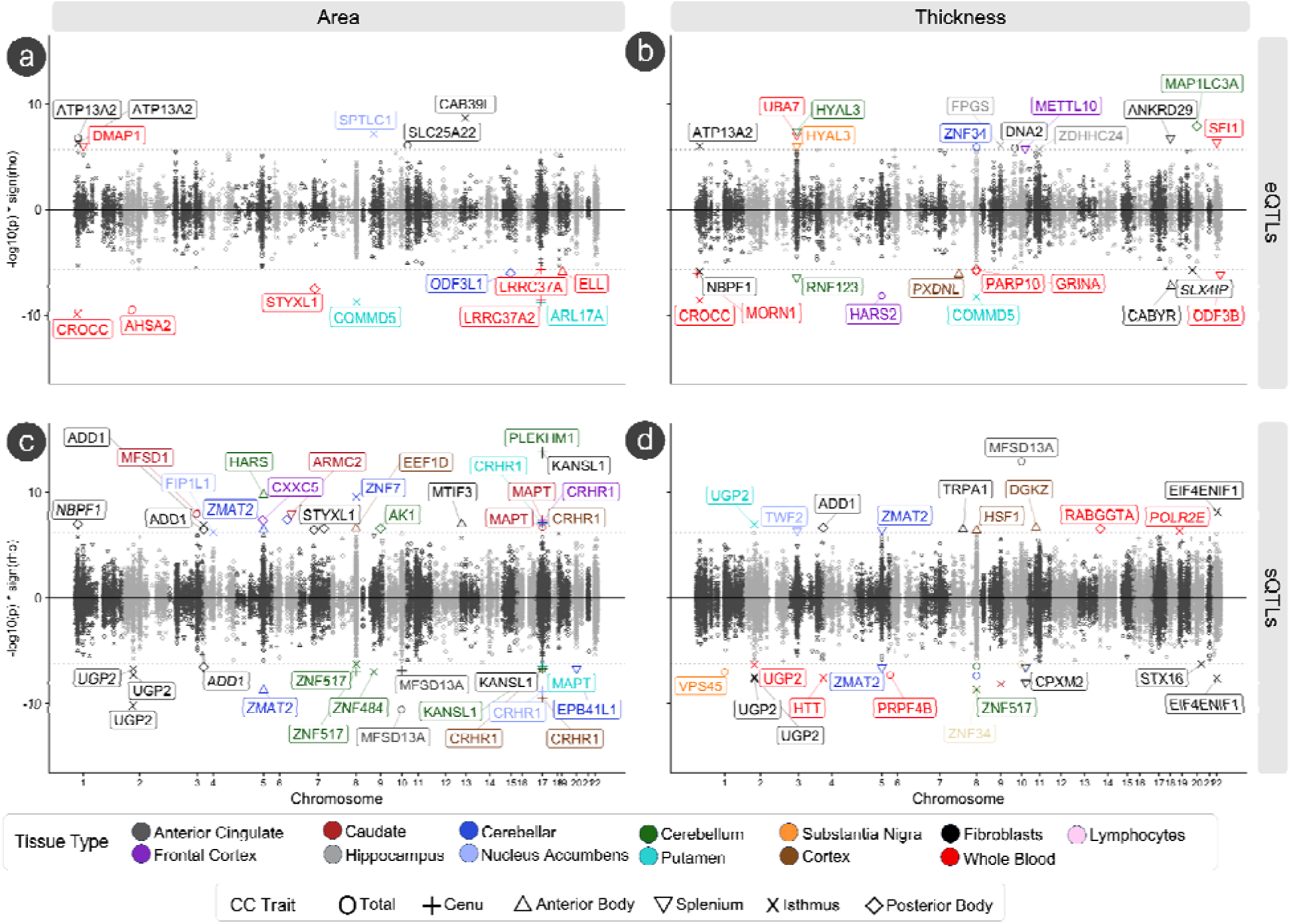
LAVA-TWAS analyses of corpus callosum traits with gene-expression (eQTLs) and splicing (sQTLs). Results of local genetic correlations between CC traits and eQTLs and sQTLs from GTEx v8 using the LAVA-TWAS framework. Associations between (A) CC area and eQTLs, (B) CC thickness and eQTLs, (C) CC are and sQTLs, and (D) CC thickness and sQTLs are shown via -log10p values scaled by the direction of association (y-axis) and chromosomal location (x-axis). All significant points are colored by tissue type and labeled by CC trait. Significance thresholds for eQTLs (*p* < 2.01 ×10^-6^) and sQTLs (*p* < 5.45 × 10^-7^) were determined by Bonferroni correction.

### Genetic overlap of corpus callosum and cerebral cortex architecture

Broadly, we observed a pattern of negative genetic correlations with area and thickness of the CC with cortical thickness across regions of the cingulate cortex, but positive genetic correlations with regions’ cortical thickness across the neocortex (**Fig. 5A**). Specifically, we observed a significant negative genetic correlation between total area with cortical thickness of the rostral anterior cingulate (*r_g_* = −0.35, *SE* = 0.06) and posterior cingulate (*r_g_* = −0.28, *SE* = 0.06). Mean thickness was negatively genetically correlated with cortical thickness of the rostral anterior cingulate (*r_g_*= −0.29, *SE* = 0.06) and posterior cingulate (*r_g_* = −0.23, *SE* = 0.05). Positive genetic correlations were observed with cortical thickness of the lingual gyrus (*r_g_*= 0.26, *SE* = 0.05) and cuneus (*r_g_* = 0.27, *SE* = 0.06). When parcellating by the Witelson scheme, negative genetic correlations were observed for area and mean thickness with cortical thickness of regions across the cortex and the cingulate, but positive genetic correlations with regions in the occipital lobe. We also observed a significant negative genetic correlation between total area of the CC with surface area of the precuneus (*r_g_*= −0.20, *SE* = 0.04). (**Supplementary Table S9-S10**).

**Figure 5:**
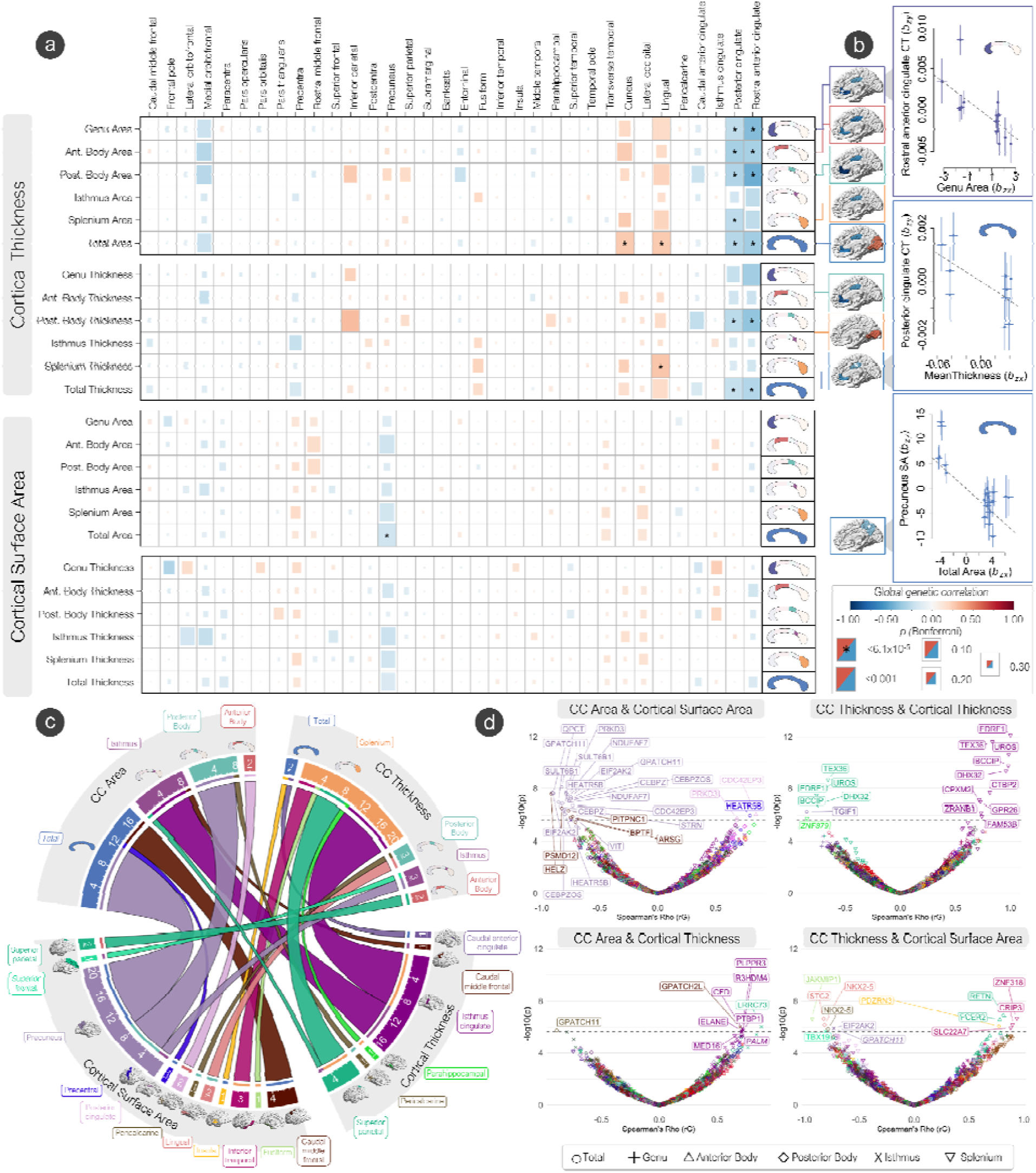
The genetic overlap of the corpus callosum and cerebral cortex. (A) Global genetic correlations (LDSC - rG) between CC phenotypes and cerebral cortex phenotypes. The Bonferroni significance threshold was set at *p* 6.1 × 10^-5^. Surface area and cortical thickness of significant cortical regions with each CC phenotype are displayed on brain plots. (B) Of the significant global genetic correlations, significant Mendelian randomization (GSMR) results ar displayed, representing the effect of CC phenotypes on cortical phenotypes free of non-genetic confounders. (C) Chord plot displaying the number of significant bivariate local genetic correlations (LAVA) between CC and cortical phenotypes. Underlined numbers represent the total number of genes shared with that phenotype. (D) Volcano plots showing degree (-log_10_ *p*-values) and direction (rG) of local genetic correlations (LAVA) between cortical and CC phenotypes. Colors represent cortical regions labeled on the chord plot in section C. Significant genes (Bonferroni significance threshold was set at *p* = 2.18 × 10^-6^) across all phenotypes are labeled.

Genetic correlations can reflect direct causation, pleiotropy, or genetic mediation. To explore potential causal relationships between CC phenotypes and morphometry of the cerebral cortex, we ran Generalized Summary-data-based Mendelian Randomization (GSMR) analyses^47^ directional effect of CC phenotypes on morphometry of the cerebral cortex, but not vice-versa. (**Fig. 5B, Supplementary Table S14**). There was a strong negative unidirectional effect of total CC area on the precuneus surface area (*b_xy_* = −0.50, *SE* = 0.13, *p* = 0.0002), implying a greater total area and thickness of the CC results in a lower surface area of the precuneus. There was also a negative unidirectional effect of total CC mean thickness and cortical thickness of the posterior cingulate (*b_xy_*= −0.02, *SE* = 0.008, *p* = 0.02), but not vice versa. When using the Witelson parcellation scheme, there was a strong negative unidirectional effect on the area of the genu on the cortical thickness of the rostral anterior cingulate (*b_xy_* = −0.001, *SE* = 0.0003, *p* = 0.003).

Local genetic correlations of area phenotypes of the CC and surface area of the cerebral cortex with LAVA^28^ showed many significant negative correlations in genes between the total area and posterior body and the precuneus SA along the 2p22.2 cytogenetic band (*QPCT, PRKD3, SULT6B1, NDUFAF7, EIF2AK2, HEATR5B, GPATCH11, CEBPZ, CEBPZOS, CDC42EP3, STRN, VIT*) (**Fig. 5C-D**). Negative genetic correlations between total CC area and caudal middle frontal gyrus SA in 5 genes along the 17q24.2 cytogenetic band (*HELZ, PSMD12, PITPNC1, ARSG, BPTF*) were also observed. Positive local genetic correlations along the 2p22.2 cytogenetic band were observed with anterior body area and the surface area of the posterior cingulate (*CDC42EP3*, *PRKD3*), as well as total area of the CC and precentral gyrus surface area (*HEATR5B*).

Many negative local genetic correlations were observed with mean thickness of the splenium and cortical thickness of the superior parietal gyrus (*TEX36, EDRF1, UROS, BCCIP, DHX32*) and the parahippocampal gyrus (*ZNF879*) along the 10q26.13-10q26.2 cytogenetic bands, while positive genetic correlations were observed with isthmus cingulate cortical thickness along the 10q26.13-10q26.2 cytogenetic bands (*EDRF1, TEX36, UROS, BCCIP, DHX32, CTBP2, CPXM2, GPR26, ZRANB1, FAM53B*).

Area of the posterior body showed a negative local genetic correlation with pericalcarine gyrus cortical thickness (*GPATCH11*). Area of the isthmus showed positive local genetic correlations with the cortical thickness of the superior parietal gyrus (*LRRC73*), caudal middle frontal gyrus (*GPATCH2L*), and isthmus cingulate (*PLPPR3, CFD, R3HDM4, PTBP1, ELANE, MED16, PALM*) along the 19p13.3 cytogenetic band.

Mean thickness of the posterior body showed negative local genetic correlations with the surface area of the lingual gyrus (*STC2, NKX2-5,* 5q35.2) and pericalcarine gyrus (*NKX2-5*). Mean thickness of the isthmus showed negative local genetic correlations with the precuneus (*EIF2AK2, GPATCH11,* 2p22.2) and superior frontal gyrus (*TBX19*) surface area. Total mean thickness of the CC showed a positive genetic correlation with surface area of the insula (*PDZRN3*). The anterior body mean thickness showed positive local genetic correlations with surface area of the superior parietal gyrus (*RETN, FCER2*). Splenium mean thickness showed positive genetic correlations with inferior temporal gyrus surface area (*ZNF318, CRIP3, SLC22A7*) along the 6p21.1 cytogenetic band.

### Genetic overlap of corpus callosum and associated neuropsychiatric phenotypes

We observed a significant genetic correlation (**Fig. 6A, Supplementary Table S11**) between total CC area and ADHD (*r_g_*= −0.11, *SE* = 0.03), bipolar disorder (BD, *r_g_* = −0.10, *SE* = 0.03), and bipolar I disorder (BD-I, *r_g_* = −0.10, *SE* = 0.03). Total mean thickness was genetically correlated with BD (*r_g_* = −0.10, *SE* = 0.03) and BD-I (*r_g_* = −0.10, *SE* = 0.03). When analyzing the regional Witelson parcellations, the area of the genu was genetically correlated with ADHD risk (*r_g_* = −0.13, *SE* = 0.03), and the mean thickness of the splenium was genetically correlated with risk for BD (*r_g_* = −0.13, *SE* = 0.03) and BD-I (*r_g_*= −0.12, *SE* = 0.03).

**Figure 6:**
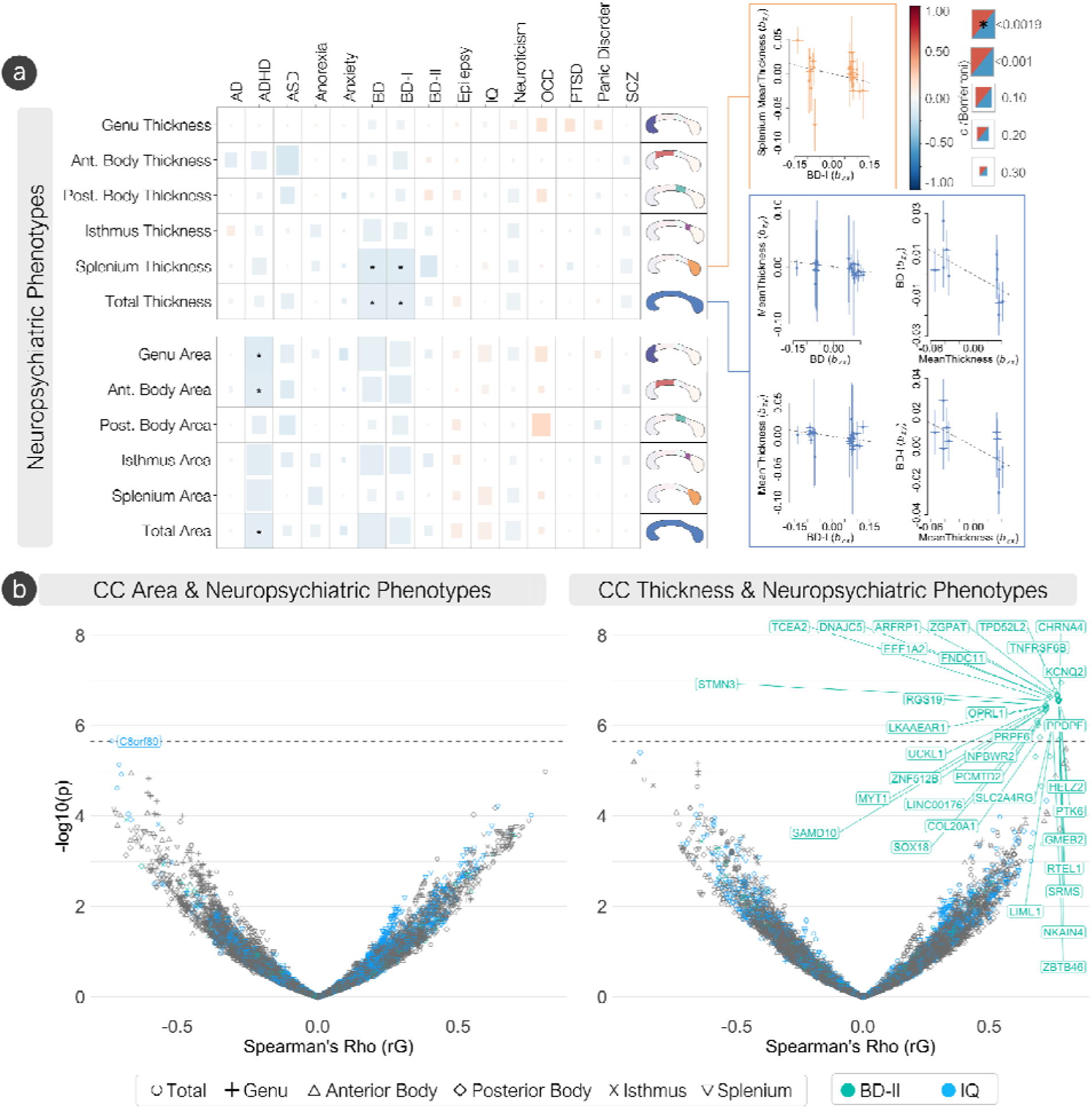
The genetic overlap of the corpus callosum and neuropsychiatric phenotypes. (A) Global genetic correlations between CC traits and neuropsychiatric phenotypes. The Bonferroni significance threshold was set at *p* = 0.0019. Of the significant global genetic correlations, significant Mendelian randomization (GSMR) results ar displayed, representing the effect of CC phenotypes on neuropsychiatric phenotypes free of non-genetic confounders. (B) Volcano plots showing degree (-log_10_ *p*-values) and direction (rG) of local genetic correlations (LAVA) between neuropsychiatric and CC phenotypes. Phenotypes with significant associations are colored (IQ an bipolar II disorder). Significant genes (Bonferroni significance threshold was set at *p* = 2.23 × 10^-6^) across all neuropsychiatric phenotypes. AD: alzheimer’s disease, ADHD: attention deficit hyperactivity disorder, ASD: autism spectrum disorder, BD: bipolar disorder, BD-I: bipolar I disorder, BD-II: bipolar II disorder, IQ: intelligence quotient, OCD: obsessive-compulsive disorder, PTSD: post-traumatic stress disorder, SCZ: schizophrenia.

GSMR analyses showed causal bidirectionality of genetic liability of BD (*b_xy_* = −0.06, *SE* = 0.02, *p* = 0.006) and BD-I (*b_xy_* = −0.05, *SE* = 0.02, *p* = 0.003) on total mean thickness of the CC, and mean thickness of the CC on BD (*b_xy_* = −0.19, *SE* = 0.08, *p* = 0.01) and BD-I (*b_xy_* = −0.23, *SE* = 0.09, *p* = 0.02). When using the Witelson parcellation, GSMR analyses showed causal directionality of genetic liability of BD-I on mean thickness of the splenium (*b_xy_*= −0.09, *SE* = 0.04, *p* = 0.01), but not *vice versa* (**Fig. 6A**, **Supplementary Table S15**).

Local genetic correlations with LAVA^28^ (**Fig. 6B**, **Supplementary Table S17**) showed 34 positive local genetic correlations between thickness of the posterior body and bipolar II disorder (BD-II) along the 20q13.33 cytogenetic band (top 5 genes being *KCNQ2, TPD52L2, TNFRSF6B, ZGPAT, ARFP1*), and one negative local genetic correlation between total CC area and IQ (*C8orf89*).

## Discussion

We performed a GWAS meta-analysis of corpus callosum morphometry using our artificial intelligence based extraction tool, *SMACC*, from 46,685 individuals using UKB and ABCD. The majority of studies investigating the genetic influence via candidate genes on CC structure and development have been conducted using various animal models and post-mortem human studies^6^. Given the difference of the human CC compared to animal models^6^, this study provides genome-wide insight into human variation and genes that influence the human CC *in vivo*.

We show the genetic architecture of the CC is highly polygenic, and specific genetic variants influence CC subregions along a rostral-caudal gradient. Five loci that were positionally mapped to genes were identified to influence both total area and mean thickness of the CC (*IQCJ-SHIP1*, *FIP1L1*, *HBEGF*, *CDKN2B-AS1*, and *FAM107B*). *IQCJ-SHIP1* had the strongest effect across total area and mean thickness, implicating mechanisms such as conduction of action potentials in myelinated cells via organizing molecular complexes at the nodes of Ranvier and axon initial segments, calcium mediated responses, as well as axon outgrowth and guidance^48^. The strongest locus for total area was mapped to the *STRN* gene. *STRN* has been heavily implicated in the Wnt signaling pathway, which controls the expression of genes that are essential for cell proliferation, survival, differentiation, and migration via transcription factors^49–51^. The *HBEGF* gene was the strongest locus for total mean thickness, implicating mechanisms in early development. *HBEGF* expression is localized in the ventricular zone and cortical layers during development^52^, and has been implicated in regulating cell migration via chemoattractive mechanisms^52^. Significant enrichment of heritability of total mean thickness in various histone marks from chromatin data (ATAC-seq) of the fetal brain and cortex derived primary cultured neurospheres, significant tissue expression in the brain 19-weeks post conception, as well as enrichment of gene sets involving regulation of histone modification, suggests genetic variants in regions of open chromatin and transcriptional activity regulation in early development are key mechanisms underlying CC morphometry. When histones are acetylated, they become more negatively charged. This negative charge repels the negatively charged DNA, causing the DNA to be “pushed away” from the histones. This loosening of the DNA-histone complex makes it easier for transcription factors to access the DNA and initiate transcription^53^.

Parcellation of the CC into the five regions defined by the Witelson scheme allowed for further refinement and genetic understanding of its morphometry in a rostral-caudal gradient. Our results provide insight as to which molecular mechanisms influence this functionally defined gradient (i.e. prefrontal, premotor/supplementary motor, primary motor, primary sensory, and parietal/temporal/occipital)^24^. An overlap of genetic loci along the most anterior (genu and anterior body, *SHTN1*) and most posterior (isthmus and splenium, *IQCJ-SCHIP1*) regions of the CC, along with splenium heritability enrichment of in histone chromatin marks of the fetal brain and dorsolateral prefrontal cortex, implicates regulation of neuron migration and action potential conduction. But the overlap of the *FOXO3* along the area of the posterior body and isthmus implicates IL-9 signaling and *FOXO*-mediated transcription responsible for triggering apoptosis^54^. Only the posterior body and isthmus showed heritability enrichment in immune cells including myeloid cells and innate lymphocytes. The thinning of the CC (along the posterior body and isthmus) occurs in a functional gradient connecting the somatosensory and parietal association areas of the brain^6,55,56^. This follows activity dependent pruning by functional area^6^, where somatosensory circuits are pruned in early development in an experience dependent context^57^. As immune cells are increasingly being recognized as key players in brain maturation and neurodevelopment^58^, our results suggest IL-9 mediating a neuroprotective effect in the CC during the cell dieback phase^58,59^, and may play a significant role in posterior CC morphometry. LAVA-TWAS results showed another potential mechanism of isthmus pruning via expression of *ATP13A2* in fibroblasts, and splicing of genes involved in NF-κB signaling^60^. *ATP13A2* is involved in lysosomal-mediated apoptosis^61^, suggesting such regulation of fibroblast mediated growth of callosal projections^62^. This is also supported by the current discovery of enrichment of genes related to isthmus area in the “*regulation of chaperone mediated autophagy pathway*”, which may influence isthmus morphometry.

The topographic organization of the CC correlates with the homotopic bilateral regions of the cortex it is known to connect^5^. A variety of genetically regulated principal mechanisms influence CC neuronal and glial proliferation, neuronal migration and specification, midline patterning, axonal growth and guidance, and post-guidance refinement to homotopic analogs in the cortex^63,64^. Our results suggest potential genetic mechanisms contributing to callosal-cortical organization. We show an overall negative global genetic correlation of CC phenotypes with the cortical thickness of the cingulate and surface area of the posterior parietal cortices, including a unidirectional negative effect of genu area on rostral anterior cingulate thickness, and total area on precuneus surface area free of any non-genetic confounders. Positive global genetic correlations of total CC area and splenium thickness with cortical thickness in the occipital cortex were also observed. Local genetic correlations of the CC were observed throughout the cerebral cortex, most pronounced with total CC area and splenium thickness. Notable findings included numerous genes in the chr2p22 cytogenetic band showing negative correlations between total CC and posterior body area with precuneus surface area, including the significant *STRN* gene observed across all CC phenotypes, further implicating the Wnt signaling pathway and dendritic calcium signaling in the context of neurodevelopment^65,66^. Within this cytogenetic band, *HEATR5B* was also positively genetically associated with precentral gyrus surface area. Opposing genetic effects were observed between splenium thickness with isthmus cingulate thickness (i.e. positive) vs. superior parietal cortex thickness (i.e. negative) in genes in the chr10q26.13 cytogenetic band. Clinical phenotypes associated with the central nervous system due to copy number variations of chr10q26.13 include abnormal cranium development, global developmental delay and learning difficulties, and neuropsychiatric manifestations including ADHD, impulsivity or autistic behaviors^67–69^. This provides a novel testable hypothesis for functional follow up studies, as alterations in the isthmus cingulate and superior parietal cortex have been observed in large-scale studies of various neurodevelopmental disorders^70^. Positive genetic associations in the chr19p13.3 cytogenetic band were observed between the isthmus area and isthmus cingulate cortical thickness, which has been implicated with microcephaly, ventriculomegaly and developmental delay^71,72^.

Our results demonstrate opposing genetic relationships between CC phenotypes and thickness of the cingulate cortex (negative) vs the neocortex (positive), which suggests a strong genetic component underlying the development of the CC via pioneer axons and chemotaxis. Developmentally, pioneer axons emerge in the cingulate and project their axons across the midline using guidance cues. A large portion of these callosal projections are pruned and myelinated in an activity dependent manner, such that axonal remodeling is highly dependent on correlated neural activity in the cortex^6,73–75^. The strongest local genetic correlation supporting this finding was observed between total mean thickness of the CC and rostral anterior cingulate thickness on *TGIF1*. As *TGIF1* is implicated in holoprosencephaly (i.e. where the brain fails to develop two hemispheres), forebrain development via alterations in the Sonic Hedgehog (SHH) pathway, and disruption of axonal guidance via chemoattractive mechanisms^76,77^, these results provide a potential genetic localization for functional follow-up. The isthmus cingulate, in relation to the isthmus and splenium, was the only cingulate region showing positive local genetic correlations, providing further evidence of distinct molecular mechanisms (e.g. immune-mediated apoptosis and regulation of callosal projections) compared to the rest of the corpus callosum underlying its structure and development.

Abnormalities of the CC have also been associated with various neurological/neuropsychiatric disorders^6^. The negative global genetic correlations observed in CC area with ADHD and CC thickness with bipolar disorder, indicate that the allelic differences resulting in smaller CC area and thickness are partly shared with those resulting in a greater risk for ADHD and bipolar disorder, respectively. Positive local genetic correlations heavily implicated the 20q13.33 cytogenetic band in the relationship between posterior body thickness and bipolar II disorder, providing a plausible neurobiological mechanism underlying an observed genetic risk at this locus^78–80^, and an observed morphological difference in the CC^13,81,82^. A negative genetic correlation at the *C8orf89* locus, known to have biased expression in the testis^83,84^, was observed between total CC area and IQ. Strong evidence suggests the high similarity in gene expression and proteome between the brain and testes are due to involvement in the speciation process^85^.

In summary, this work identifies genome-wide significant loci of morphometry of the overall corpus callosum and its sectors, convergence on biological functions, tissues and cell types, as well as the genetic overlap with the cerebral cortex and neuropsychiatric conditions.

## Methods

### Artificial intelligence corpus callosum extraction and segmentation with SMACC

#### Data Preprocessing

All UKB participants completed a 31-minute neuroimaging protocol using a Siemens Skyra 3 Tesla scanner and a 32-channel head coil in one of three MRI scanning locations. All 3D structural T1-weighted brain scans were acquired using the following parameters: 3D MPRAGE, sagittal orientation, in-plane acceleration factor = 2, TI/TR = 880/2000 ms, voxel resolution = 1 x 1 x 1 mm, acquisition matrix = 208 x 256 x 256 mm. All scans were pre-scan normalized using an on-scanner bias correction filter. More details of the imaging protocols may be found in the following reference papers^86,87^.

All ABCD participants completed a neuroimaging protocol in one of three scanner types at 21 different sites^88^. The Siemens Prisma had the following parameters for the T1-weighted scans: TI/TR = 1060/2500 ms, TE = 2.88 ms, voxel resolution = 1 x 1 x 1 mm, acquisition matrix = 176 x 256 x 256, flip angle = 8 degrees. The Philips Achieva Ingenia had a TI/TR = 1060/6.31 ms, voxel resolution = 1 x 1 x 1 mm, acquisition matrix = 225 x 256 x 256 mm and a flip angle = 8 degrees. The GE MR750 had a TI/TR = 1060/2500 ms, TE = 2 ms, voxel resolution = 1 x 1 x 1 mm, acquisition matrix = 208 x 256 x 256, and a flip angle = 8 degrees.

All T1w MRIs were registered to MNI152^89–91^ 1mm space with 6 degrees of freedom using FSL’s *flirt*^92^ command.

#### SMACC development and UNet training

Mid-sagittal T1w, T2w, and FLAIR images from UK Biobank^21^, PING^93^, HCP^94^, and ADNI1^95^ were used for training the UNet model for CC segmentation. Individual study scanner parameters can be found in their respective references. The demographic information for the datasets used to create the UNet model is shown in **Supplementary Table 31**. Augmentation of image data is a common procedure in deep learning to prevent model overfitting and improve model accuracy^96^. All the images were downsampled by a factor of 2, 3, 4 and 5 along the sagittal axis and then upsampled back to original size using MRtrix’s *mrgrid* command to include low resolution images in the training^97^. To include lower resolution T1w images resembling older or clinical data in training, all the images were harmonized using a fully unsupervised deep-learning framework based on a generative adversarial network (GAN)^98^ to a subject from the ICBM dataset^90^. Images were also rotated clockwise in increments of 15 degrees and then resized to 256*256. Black boxes were randomly added to the images to imitate partial agenesis cases. **Supplementary Figure 1** shows some T1w augmented images that were the input training images for the UNet model.

#### UNet Implementation

A Tensorflow implementation of UNet^99^ was trained on 80% of the images for 250 epochs until the difference between the intersection over union (IOU) after consecutive iterations was less than 1×10^−4^. The U-Net architecture is structured with a contracting pathway and an expansive pathway. The contracting pathway repeatedly performs two 3×3 convolutions (without padding), with each convolution followed by a rectified linear unit (ReLU) and a 2×2 max pooling operation. At each stage in the expansive pathway the feature map is upsampled followed by a 2×2 convolution which reduces the feature channels by half. Then, the corresponding cropped feature map from the contracting pathway is concatenated, and two 3×3 convolutions are applied, with each one followed by a ReLU. We used the following training parameters: 1×10^−4^ learning rate and an Adam optimizer^100^. The rest of the data was used for validation. The midsagittal CC (midCC) was initially segmented using image processing techniques^101^ on subjects from ADNI1 (N=1032, 54-91 years), PING (N=1178, 3-21 years), HCP (N=963, 22-37 years) and UKB (N=190, 45-81 years). These masks were then visually verified and manually edited by neuroanatomical experts which served as the ground truth. To evaluate the model, the area of overlap between the predicted segmentation and the ground truth was calculated.

#### CC shape metrics extracted with SMACC

SMACC provides outputs of global and regional shape metrics extracted from the corpus callosum segmentation, including area, thickness, length, perimeter and curvature. The regional shape metrics were based on a 5 compartment version of the Witelson atlas^23,24^. The Witelson atlas is composed of the (1) genu, (2) anterior midbody, (3) posterior midbody, (4) isthmus, and (5) splenium. The metrics used for the GWAS analysis were area and mean thickness of the total CC and all of the parcellations of the Witelson atlas. The thickness is defined as the distance in the inferior-superior direction between the top and bottom of the contour and at every point along the length of the segment, then averaged across the region of interest. The total area is the summation of the number of voxels with intensity value greater than 0.5 in the segmentation.

#### Corpus callosum segmentation quality control (QC) with SMACC

To ensure that segmentations were of appropriate quality without having to manually assess all output images, which eventually may scale to hundreds of thousands of scans, we included an automated quality control (QC) assessment into SMACC. The regional and global metrics were used as inputs to the machine learning models detailed below for automatic binary classification of segmentations as Pass or Fail. CC segmentations from SMACC were manually assessed across multiple datasets by neuroanatomical experts. This included data from UKB (N=12,902, aged 45-81 years), ADNI1 (N=724, aged 54-91 years), PING (N=857, 3-21 years) and HCP (N=615, 22-37 years), all of which served as the ground truth for QC model building. All data was split 80/20 for training/testing.

Figure 7 gives the overview and the flow of *SMACC*. Several architectures including a 3-layer sequential neural network with 42 neurons, 22 in the second layer, and 11 in the third layer; a wide & deep neural network with 80 neurons in the first 3 layers and 40 in the last 3 layers, XGBoost classifier and an ensemble model were tested to classify the segmentations from the UNet as pass or fail. The ensemble model consisted of XGBoost, k-nearest neighbors (KNN), support vector classifier (SVC), logistic regression, and a random forest classifier. The results from all the classifiers in the ensemble model were combined using a majority voting classifier. All the models were compared using metrics including precision, recall, F1 score and Area Under the Curve (AUC). **Supplementary Table 34** shows the performance of different models based on the shape metrics extracted from the CC segmentations.

**Figure 7:**
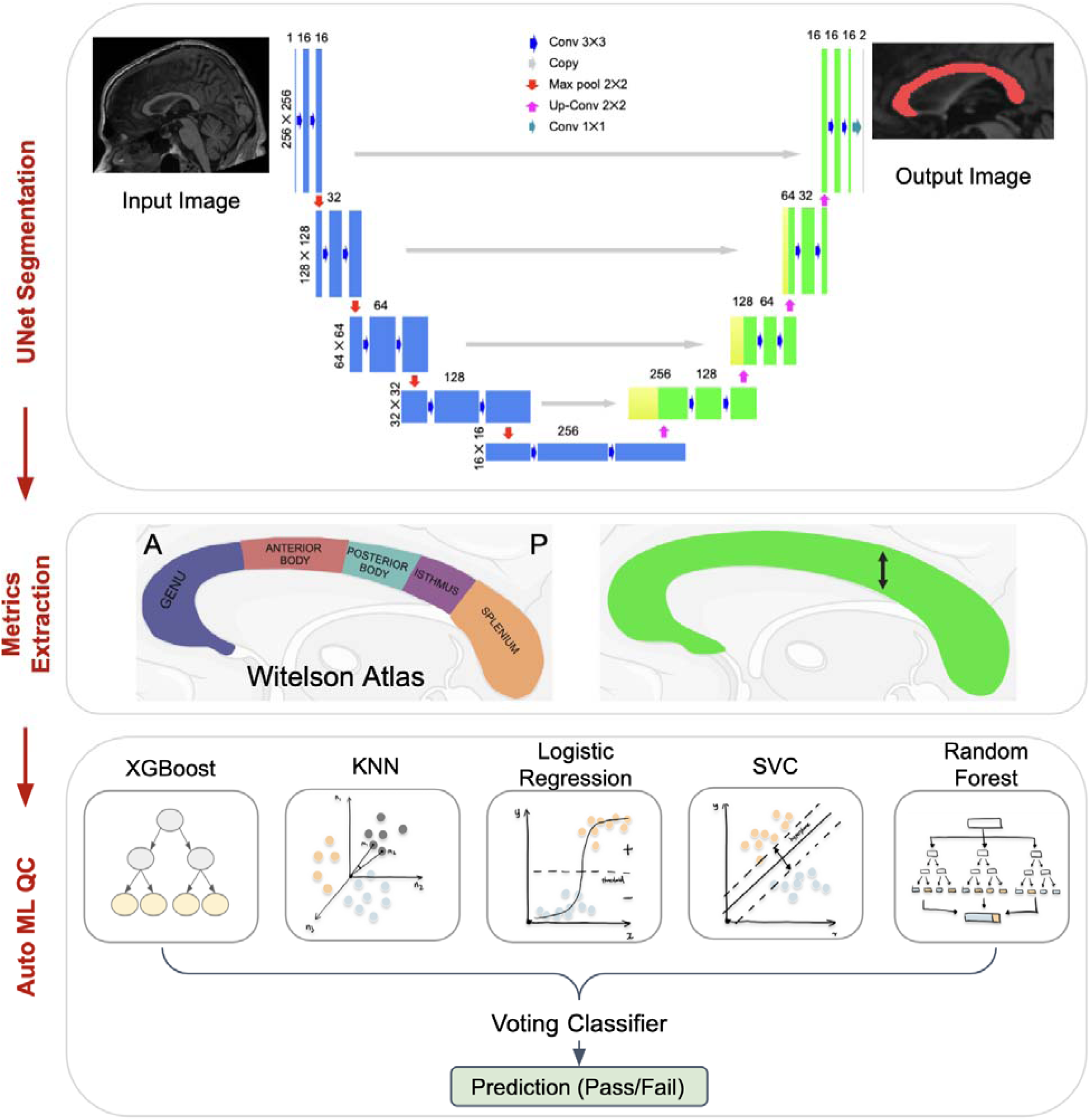
Segment, Measure, and AutoQC the midsagittal CC (SMACC) pipeline. The midsagittal slice from a participant registered to MNI space with 6 degrees of freedom serves as an input to the UNet architecture used for the midsagittal corpus callosum segmentation. The Witelson atlas was used for segmenting the CC into five different regions. Global and subregion metrics (thickness and area-shown in green) were extracted from the segmentation. The thickness (black arrow) is defined as the distance in the inferior-superior direction between the top and bottom of the contour, after reorientation to standard space, at every point along the length of the segment, then average across the region of interest. These metrics serve as input for the ensemble machine learning model used for labeling CC segmentations as having passed or failed quality control (QC). Abbreviations: Montreal Neurological Institute - MNI, CC - corpus callosum, ML - Machine Learning, KNN - K Nearest Neighbors, SVC - Support Vector Classifier

#### SMACC vs FreeSurfer

##### Comparing SMACC and FreeSurfer via Dice scores with respect to manual masks

For assessing the accuracy of the *SMACC* compared to the ground truth and compared to the commonly used tool FreeSurfer^102^, we ran the *SMACC* pipeline on 30 subjects from the Hangzhou Normal University (HNU) test-retest dataset^103,104^. Each subject in this dataset was scanned with a full brain T1w MRI 10 times within a period of 40 days, for a total of 300 scans. All 300 scans had also been manually segmented by a neuroanatomical expert to serve as the ground truth. Segmentations from *SMACC* and FreeSurfer v7.1 were compared to manual segmentations using the Dice overlap coefficient. The average Dice coefficient between automated CC masks from *SMACC* and ground truth segmentations was 0.94 across all scans. The average Dice score between FreeSurfer CC segmentations and manual masks was 0.82. The Dice score was consistently higher for all the subjects using *SMACC*. **Supplementary Figure 2 and Supplementary Table 35**, show a few midCC segmentations obtained from *SMACC* compared to FreeSurfer.

#### ICC for SMACC

To assess test-retest reliability of *SMACC* the intraclass correlation (ICC) scores were calculated. Average ICC values for thickness and area of the Witelson parcellations and the total CC were greater than 0.9 and are shown in **Supplementary Figure 3**.

### Study cohorts

#### U.K. Biobank

The UK Biobank (UKB) is a large population level cohort study conducting longitudinal deep phenotyping of around 500,000 participants in the United Kingdom (UK) aged between 40-69 at recruitment. All participants provided informed consent to participate. The North West Centre for Research Ethics Committee (11/NW/0382) granted ethics approval for the UK Biobank study^21^. We used genotype data from UKB released in May 2018. The data was collected from 489,212 individuals, and 488,377 of those individuals passed quality control checks by UKB. The genotypes were then imputed using two reference panels: the Haplotype Reference Consortium (HRC) reference panel and a combined reference panel of the UK10K and 1000 Genomes projects Phase 3 (1000G) panels^21^. There were 8,422,770 SNPs following quality control (QC) of the data which included having a genotyping call rate (SNPs missing in individuals) of greater than 95%, removing variants with a minor allele frequency less than 0.01 (1%), removing variants with Hardy-Weinberg equilibrium p-values less than 1e-6, and removing individuals with greater than three standard deviations away from the mean heterozygosity rate. To determine European ancestry in UKB, the ENIGMA MDS protocol (https://enigma.ini.usc.edu/protocols/genetics-protocols/) was completed using 10 components. The mean and standard deviations of the first and second genetic components of individuals who were classified as Utah residents with Northern and Western European ancestry from the CEPH collection (CEU) from the HapMap 3 release were then calculated. Individuals in UKB who were within a distance of 0.0101 on components 1 and 2 were classified as of European ancestry (N = 41,979). The MDS plot of individuals included in the analysis overlaid over the HapMap 3 population is available in **Supplementary Figure 4**.

#### ABCD

The Adolescent Behavioral Cognitive Development (ABCD) study is the largest study in the United States (USA) following adolescent children starting from 9 years of age through adolescence with deep phenotyping including neuroimaging and genotyping using the Smokescreen™ Genotyping array consisting over over 300,000 SNPs^88,105,106^. Only neuroimaging from baseline (ages 9-10) were used. Following imputation using the ENIGMA protocol^107^ with the European 1000 Genomes Phase 3 Version 5 reference panel, phased using Eagle version 2.3^108^, and the QC process as described in the UKB cohort, a total of 4,706 European ancestry children, and 5,683,360 SNPs were included. To determine European ancestry in ABCD, the methods described for the UKB were completed. The MDS plot of individuals included in the analysis overlaid over the HapMap 3 population is available in **Supplementary Figure 5**.

We also analyzed non-European ancestry individuals to examine the generalization of the observed effects across ancestries. Using the aforementioned methods, we included 1504 individuals from the UKB and 5536 individuals from ABCD.

### GWAS meta-analysis of corpus callosum morphometry

Genome-wide association analysis (GWAS) for UKB and ABCD separately for all CC phenotypes were completed via a linear whole-genome ridge regression model using REGENIE, allowing for the control of genetic relatedness^109^. Covariates included age, sex, age*sex interaction, and the first 10 genetic principal components. A two-step REGENIE analysis was completed with the following parameters. For step 1, the entire dataset was used with a block size of 1000 and leave-one-out-chromosome validation^109^. Step 2 was completed with a threshold for minor allele count of 5, a block size of 1000, and otherwise default parameters.

A meta-analysis of GWAS summary statistics of all CC derived metrics in UKB and ABCD were conducted using METAL software and the random-metal extension^29,30^, based on the random-effects model. A random-effects model was chosen since the effect sizes of SNPs on the corpus callosum has the potential to be different between the UKB and ABCD cohorts due to age. White matter volume is known to increase through childhood and start decreasing in middle adulthood^110^, which may result in different genetic effect sizes being observed. We opted to conduct a meta-analysis instead of using a two-stage discovery-replication approach because Skol et al. have shown that this method is more powerful, despite using more stringent significance levels for multiple correction^111^, and is common practice in the literature^112–114^. Percent variance (*R*^2^) explained by each significant SNP was calculated using the approach described in Rietvield et al^115^. The *R^2^* of each variant *j* was calculated via:

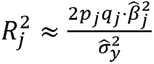

where *p_j_* and *q_j_* are the minor and major allele frequencies, *β̂* is the estimated effect of the variant within the meta-analysis and □□2 is the estimated variance of the trait (for which we used the pooled variance of the trait across UKB and ABCD. In order to determine the number of independent traits, matrix spectral decomposition was computed using matSpD in R on the phenotypic correlations between CC traits using the method proposed by Li and Ji^116,117^. This resulted in 8.16 effective independent variables, and a significance threshold of *p* = 5 × 10^-8^/8.16 = 6.13 × 10^-9^. Meta-analyses were also completed for non-European individuals.

### Heritability and genetic correlations within and between cohorts

To determine SNP heritability (*h^2^_SNP_*) tagged from SNPs used in the analysis, we used the GREML approach implemented in GCTA^38,39^, while adjusting for the same covariates as in the GWAS. The SNP heritability (*h^2^_SNP_*) from LDSC,^27^ was also computed, which estimates heritability casually explained by common reference SNPs. Genetic correlations between the UKB and ABCD cohorts for area and thickness of each parcellation of the CC defined by the Witelson scheme, and total CC were completed using LDSC^27^. Between cohort heterogeneity of *h^2^_SNP_* should not be considered unusual, as the genetic influence observed on the corpus callosum has the potential to be different between the UKB and ABCD cohorts due to age - white matter volume is known to increase through childhood and start decreasing in middle adulthood^110^, as well as the smaller sample size in ABCD making it harder for LDSC to detect polygenic effects^118^.

### Gene-mapping and gene enrichment analyses

Genetic variants (SNPs) were mapped to genes using information about genomic position, expression quantitative trait loci (eQTL) information, and 3D chromatin interaction mapping as implemented in FUMA v1.5.2 with the experiment-wide significance threshold (*p* = 6.13 × 10^-9^)^119^. Pathway enrichment analyses using the results from the full meta-analyses with no pre-selection of genes via MAGMA v1.08^40^ gene-set analysis in FUMA. Genes located in the MHC region were excluded (hg19: chromosome 6: 26Mb - 34Mb). There were 19,021 gene sets from MSigDB v7.0^120^ (Curated gene sets: 5500, GO terms: 9996), and 9 other data resources including KEGG, Reactome and Biocarta (https://www.gsea-msigdb.org/gsea/msigdb/collection_details.jsp#C2). MAGMA uses gene-based P-values to identify genes that are more strongly associated with a phenotype than would be expected by chance. MAGMA then applies a competitive test to compare the association of genes in a gene set to the association of genes outside of the gene set. This allows MAGMA to identify gene sets that are enriched for association signals. MAGMA corrects for a number of confounding factors, such as gene length and size of the gene set, to ensure that the results are not due to chance. A gene-based association analysis (GWGAS) in MAGMA was completed using the full summary statistics for each trait from METAL. Corrections for multiple comparisons were completed using the Bonferroni approach.

To determine whether genes associated with CC morphometry cluster into biological functions, tissue types, or specific cell types, we used the full results of the meta-analyzed genome-wide association studies (GWAS) rather than prioritizing genes. Pathway analysis as described above was completed.

We performed gene-property and gene-set analysis using the MAGMA software on 54 tissue types from the GTEx v8 database and BrainSpan^42,43^, which includes 29 samples from individuals representing 29 different ages of brains, as well as 11 general developmental stages.

Single cell RNA-sequencing data sets used in the cell-type specific analyses included the human developmental and adult brain samples from the PsychENCODE consortium^121^, human brain samples of the middle temporal gyrus and lateral geniculate nucleus from the Allen Brain Atlas^122^, human brain samples using DroNc-seq^123^, two datasets of human prefrontal cortex brain samples across developmental stages which show per cell type average across different ages, and per cell type per age average expression^124^, two datasets of human brain samples with and without fetal tissue^125^, human brain samples from the temporal cortex^126^, and human samples from the ventral midbrain from 6-11 week old embryos^127^. A 3-step workflow is implemented in FUMA to determine association between cell-type specific expression and CC morphometry-gene association supported by multiple independent datasets, which has been extensively described^45^. All tests were corrected using the Bonferroni approach.

### Partitioned heritability of meta-analysis results by cell and tissue type with LDSC

Partitioned heritability analysis was completed to estimate the amount of heritability explained by annotated regions of the genome^41,44^. We tested for enrichment of CC *h^2^* of variants located in multiple tissues and cell types using the LDSC-SEG approach, with all analyses being corrected for the FDR^44^. Annotations indicating specific gene expression in multiple tissues/cell types from the Genotype-Tissue Expression (GTEx) project and Franke lab were downloaded from https://alkesgroup.broadinstitute.org/LDSCORE/LDSC_SEG_ldscores/. We also downloaded 489 tissue-specific chromatin-based annotations from narrow peaks for six epigenetic marks from the Roadmap Epigenomics and ENCODE projects^128,129^. These annotations were downloaded from the URL mentioned above. This would allow us to either verify or identify new findings from the gene expression analysis from an independent source using a different type of data. Finding new patterns of chromatin enrichment can help us to understand how genes are regulated. For example, if we find that a particular epigenetic mark is enriched in a region of the genome that is associated with a specific gene in a specific tissue type, this could suggest that the gene is regulated by that epigenetic mark in that specific tissue type. Gene expression data from the Immunological Genome (ImmGen) project^130^, which contains microarray data on 292 immune cell types from mice, was used to test immune cell-type-specific enrichments. Data was downloaded from the aforementioned link.

### LAVA - TWAS

We used the LAVA-TWAS framework to investigate the relationship between CC traits and gene expression in brain tissues, fibroblasts, lymphocytes, and whole blood from the GTEx consortium (v8)^83^ in all protein coding genes, as it has ability to model the uncertainty of eQTL effects compared to other commonly used TWAS approaches, which have been shown to be prone to high type-I errors (false positives), and provides a directly interpretable effect size in the rG estimate^46^. Analyses were performed on all protein coding genes (N = 18,380) between all CC phenotypes and eQTLs/sQTLs for each tissue. Genotype data from the European sample of the 1000 Genomes (phase 3) project^131^ was used to estimate SNP LD for LAVA. For each eQTL/sQTL that had a significant genetic signal for both the CC phenotype and cortical phenotype (univariate *p*-values less than 1 × 10^-4^), the local bivariate genetic correlation between the two was estimated and tested. All LAVA-TWAS results were corrected using the Bonferroni approach. Following TWAS, trait specific enrichment analysis via a Fisher’s exact test of the top 1% of genes, to evaluate overrepresentation in 7,246 MSigDB v6.2^132^ gene sets and gain insight into biological pathways, was conducted. Gene sets were subset such that they must have consisted of at least one of the top 1% of genes, to avoid testing gene-sets with no significantly associated genes. All enrichment testing for eQTLs and sQTLs was performed with Bonferroni correction.

### Global and local genetic correlations with cortical morphometry and mendelian randomization

The CC develops in such a manner that callosal projections are over-produced then refined during development. The majority of cortical projections are refined during postnatal stages and are under the influence of guidance cues^6^. As many genes are responsible for callosal axon guidance, we sought to investigate the genetic relationship between our derived CC traits and the genetic architecture of the human cerebral cortex^6^. We used LDSC to determine the global genetic correlation between area and thickness of the total and parcellated regions of the corpus callosum, and the GWAS summary statistics of each globally corrected region-of-interest of the cerebral cortex from the ENIGMA-3 GWAS^133^. We performed bi-directional Mendelian Randomization analyses to investigate if significant genetic correlations observed could be driven by genetic causal relationships between an exposure (e.g., area and thickness of different regions of the CC) and outcome (e.g., regional surface area & cortical thickness). Analyses were performed with summary statistics using GSMR^47^. All analyses were corrected using the Bonferroni approach. To capture potential local shared genetic effects across the genome, we ran LAVA^28^ for all protein coding genes (N = 18,380) between all CC phenotypes and surface area and cortical thickness of regions in the ENIGMA3 GWAS. Genotype data from the European sample of the 1000 Genomes (phase 3) project^131^ was used to estimate SNP LD for LAVA. For each gene that had a significant genetic signal for both the CC phenotype and cortical phenotype (univariate *p*-values less than 1 × 10^-4^), the local bivariate genetic correlation between the two was estimated and tested. All results were corrected using the Bonferroni approach.

### Global and local genetic correlations with neuropsychiatric conditions and mendelian randomization

Abnormalities of the corpus callosum have also been heavily implicated in several neurological and neuropsychiatric conditions such as autism spectrum disorders (ASDs), ADHD, bipolar disorder, schizophrenia, visual impairments and epilepsy^9,56,134–141^. We used LDSC to determine the global genetic correlation between area and thickness of the total and parcellated regions of the corpus callosum, and 15 neuropsychiatric traits. Mendelian randomization analysis, and local genetic correlations were run as done for the brain cortical phenotypes.

## Data Availability

This work is a meta-analysis. Upon publication, the full meta-analytic summary statistics will be made available in ENIGMA-Vis^142^.

## Code Availability

The code and model used to extract the corpus callosum and its metrics is available at https://github.com/USC-LoBeS/smacc/.

## Acknowledgements

This work was supported by the National Institutes of Health (Grant Nos. R01 MH1340004 and R01 AG059874 [to NJ], National Science Foundation Graduate Research Fellowship Program (Grant No. 2020290241 [to RRB], the Adolescent Brain Cognitive Development (ABCD) Study (https://abcdstudy.org), and UK Biobank (Resource Application No. 11559). SEM was supported by NHMRC grants APP1172917 and APP1158127.

## Ethics Declarations

