## Supplementary Information for "The Genetic Architecture of the Human Corpus Callosum and its Subregions"

Ravi R. Bhatt,^1*^ Shruti P. Gadewar,^1*^ Ankush Shetty,^1^ Iyad Ba Gari,^1^ Elizabeth Haddad,^1^ Shayan Javid,^1^ Abhinaav Ramesh,^1^ Elnaz [Nourollahimoghadam](https://arxiv.org/search/q-bio?searchtype=author&query=Nourollahimoghadam%2C+E),^1^ Alyssa H. Zhu,^1^ Christaan de Leeuw,^2^ Paul M. Thompson,^1^ Sarah E. Medland,^3^ Neda Jahanshad^1^

^1^Imaging Genetics Center, Mark and Mary Stevens Neuroimaging and Informatics Institute, Keck School of Medicine, University of Southern California, Marina del Rey, CA, USA

^2^Department of Complex Trait Genetics, Centre for Neurogenomics and Cognitive Research, VU University, Amsterdam, The Netherlands

^3^Psychiatric Genetics, QIMR Berghofer Medical Research Institute, Brisbane 4006, Australia

*Co-first authors

**Supplementary Figure 1:** **Data augmentation techniques** - **A.** A T1w MR image **B**. Downsampled MR images by a factor of 2, 3, 4 and 5. **C**. MR images rotated in increments of 15 degrees. **D**. Black boxes of various width X height ((100 X 60), (60 X 100), (50 X 30), (30 X 50)) were added at random locations in the midCC slice to imitate partial agenesis cases.
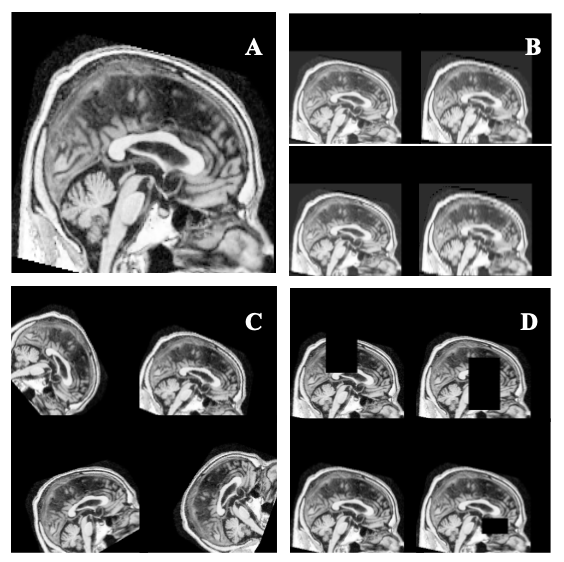

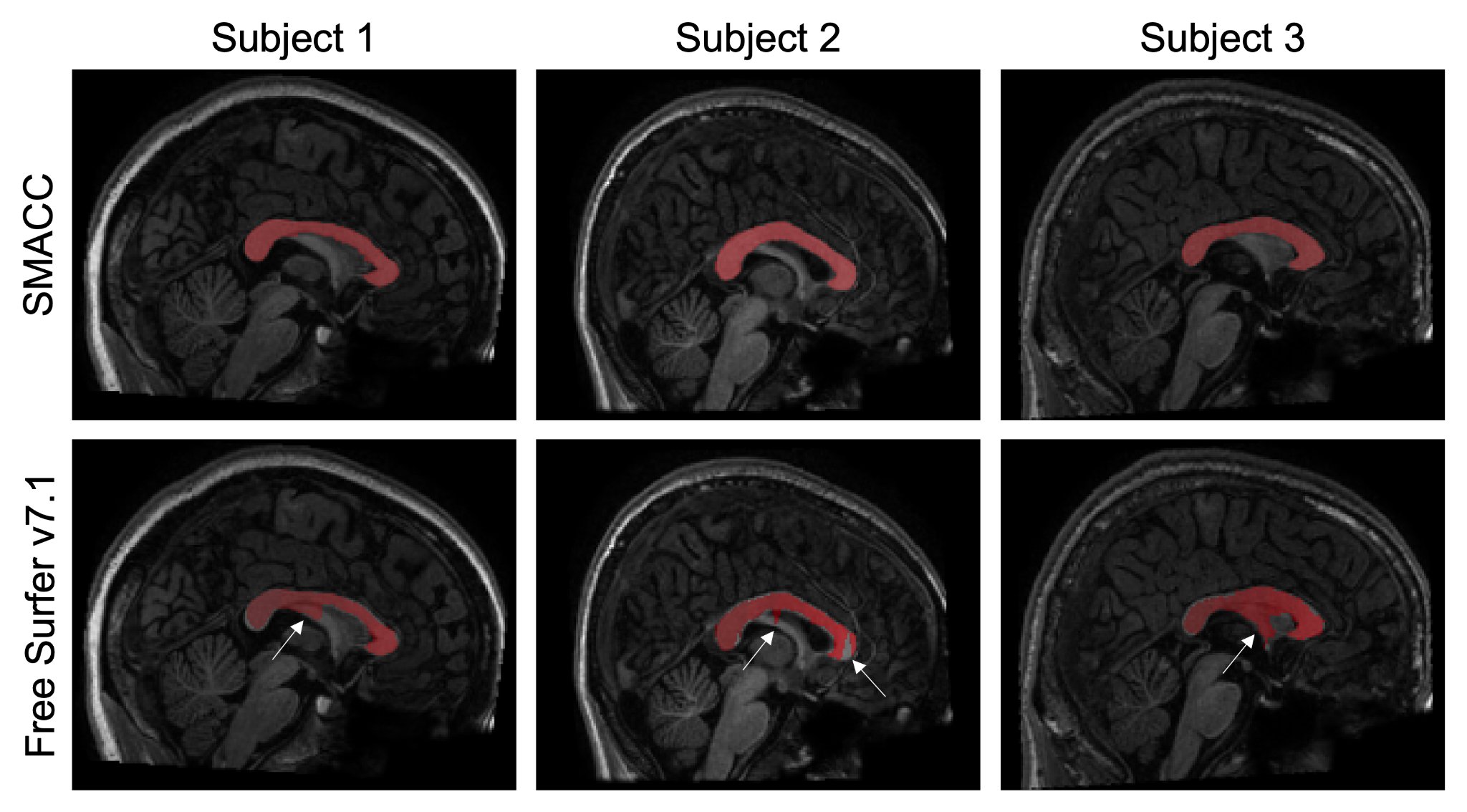

**Supplementary Figure 2**: **Midsagittal corpus callosum segmentation via SMACC:** Examples of midsagittal corpus callosum (midCC) segmentation from three different participants (left to right) using our SMACC tool^1^ (top) and FreeSurfer (bottom) in Hangzhou Normal University (HNU) dataset^2^.

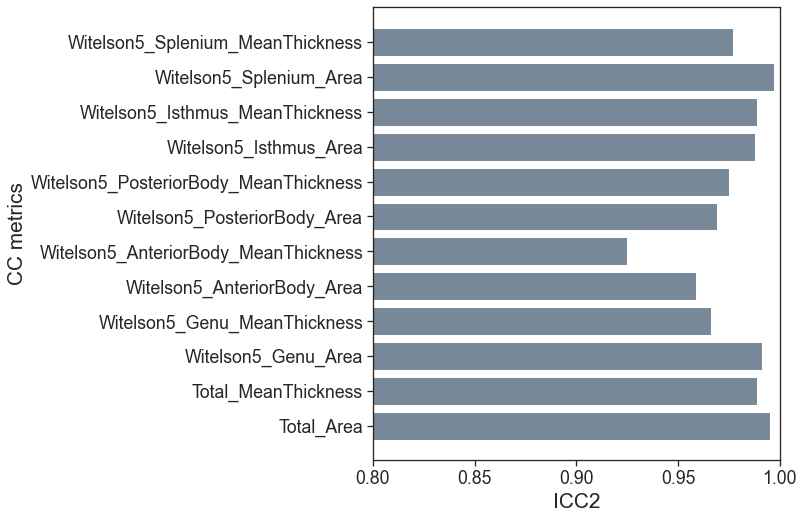

**Supplementary Figure 3**: **Reliability of SMACC metrics in a test-retest dataset:** Average intraclass correlation (ICC) coefficients for different CC metrics between sessions for all subjects in Hangzhou Normal University (HNU) dataset^2^. Higher ICC shows that our segmentations across all the sessions for a subject are very similar and hence highly reliable.

**Supplementary Figure 4:** **UK Biobank MDS in European Ancestry Individuals:** European individuals (red points) in UK Biobank^3^ as defined via MDS. UK Biobank sample is overlaid with the HapMap3 data release^4^. CEU individuals represent Utah residents with Northern and Western European ancestry from the Centre d′Etudes du Polymorphisme Humain (CEPH) collection. TSI individuals represent Toscani in Italy.
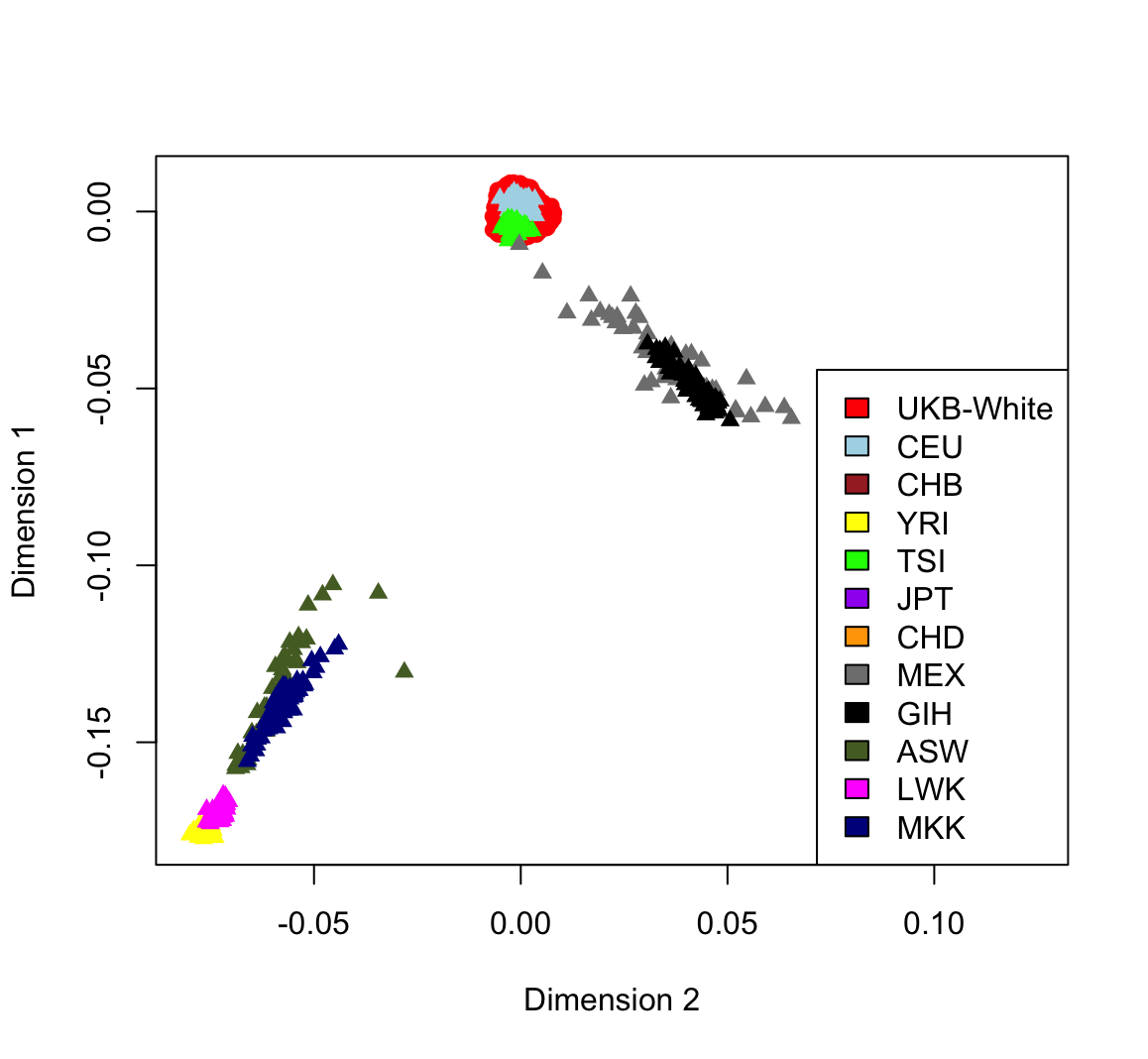

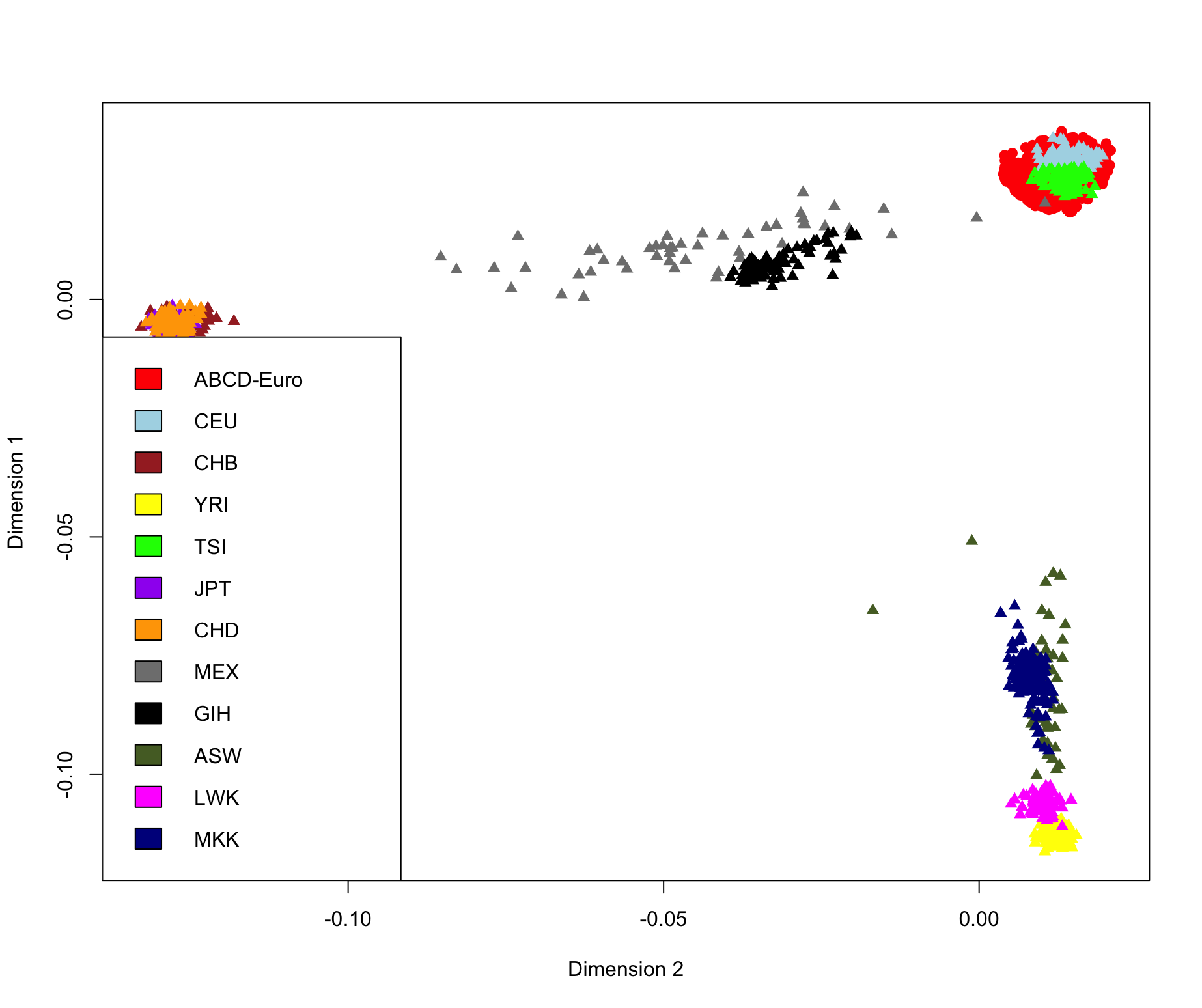

**Supplementary Figure 5**: **Adolescent Behavioral Cognitive Development study MDS in European Ancestry Individuals:** European Individuals (red points) in the Adolescent Behavioral Cognitive Development study (ABCD) as defined via MDS^5^. The ABCD sample is overlaid with the HapMap3 data release^4^. CEU individuals represent Utah residents with Northern and Western European ancestry from the Centre d′Etudes du Polymorphisme Humain (CEPH) collection. TSI individuals represent Toscani in Italy.

**Regional association plots**: Regional association plots of the meta-analyzed SNP results for each corpus callosum trait. These plots are provided in the **Supplementary Data 1**.
