## Extended Data for "The Genetic Architecture of the Human Corpus Callosum and its Subregions"

Locus 1, SDHB, Total Area, rs12073028

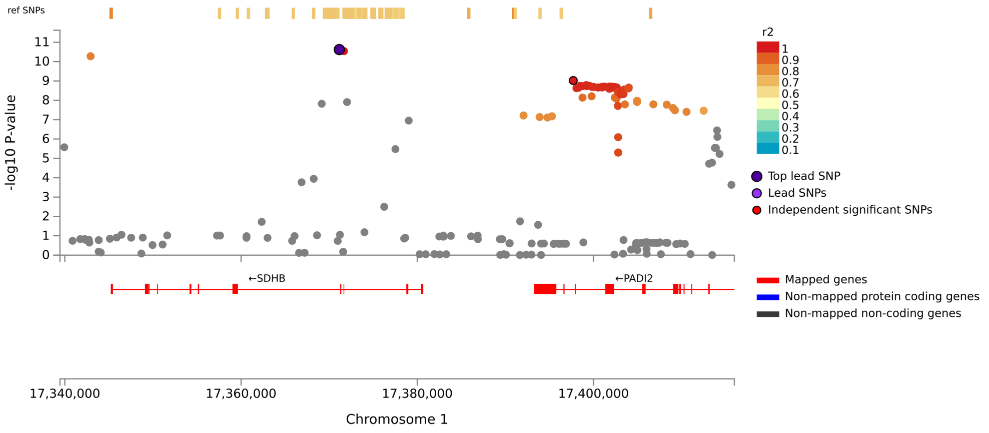


Locus 2, SDCCAG8, Total Area, rs2994330

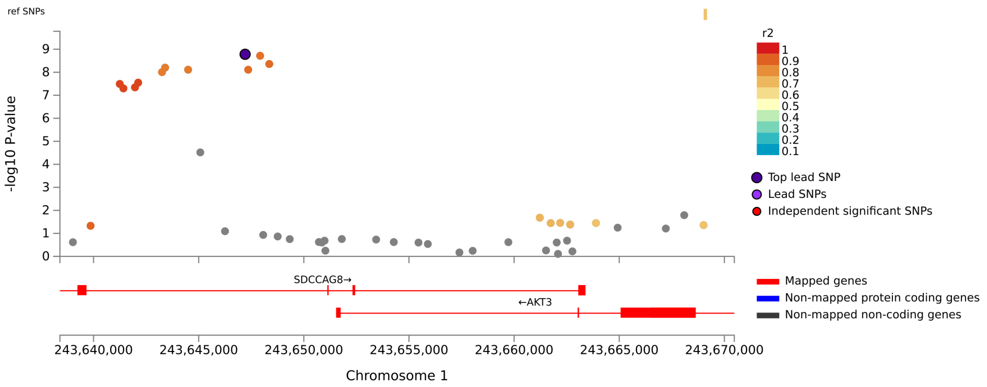

Locus 3, STRN, Total Area, rs7561572

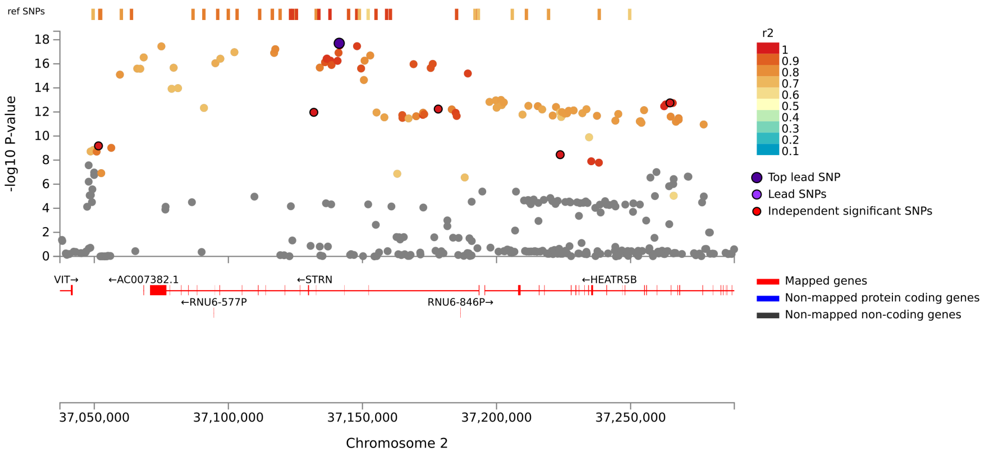


Locus 4, AC016727.1, Total Area, rs778760

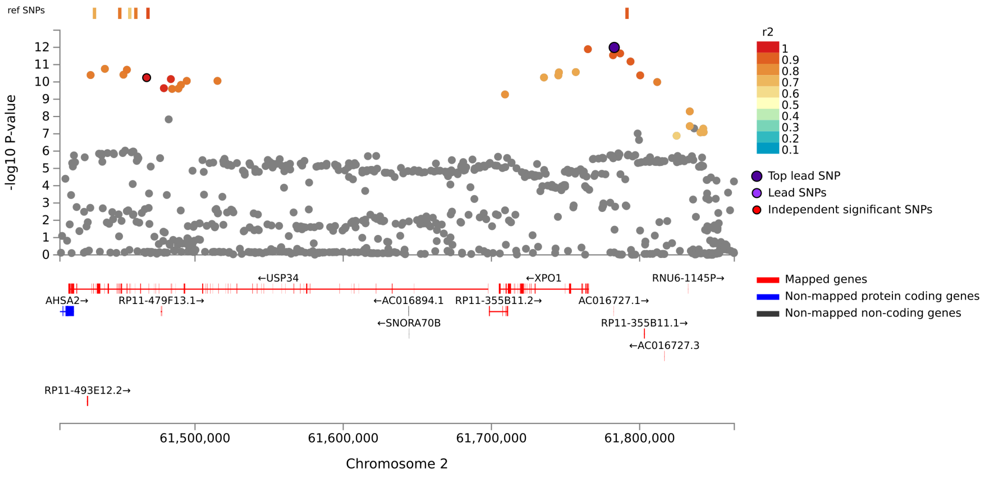

Locus 5, FAM171B, Total Area, rs17750683

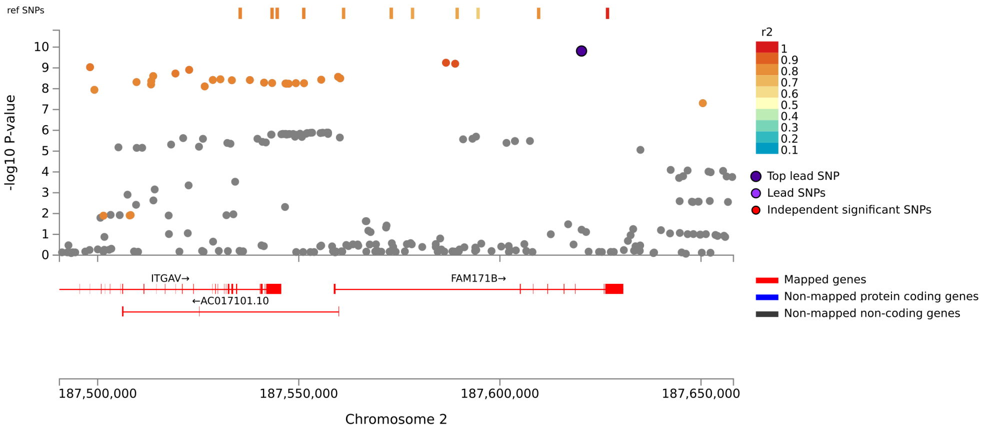


Locus 6, RP11-190P13.2, Total Area, rs4831182

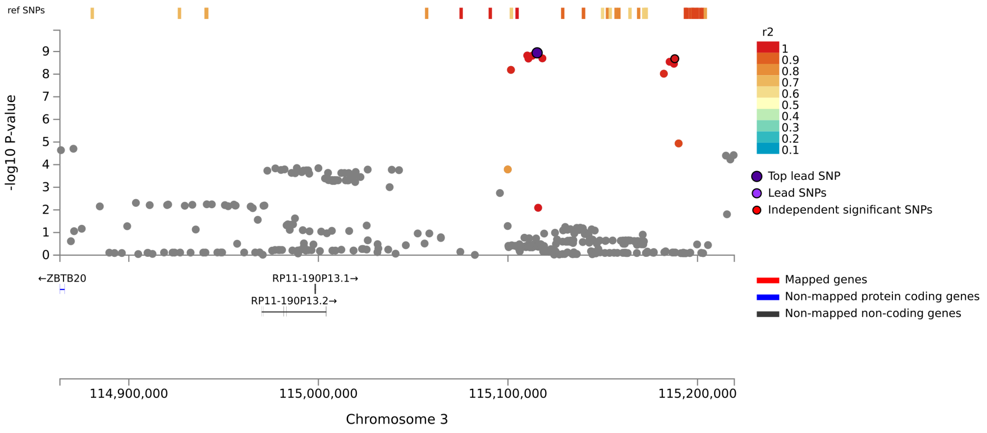

Locus 7, IQCJ-SCHIP1:IQCJ, Total Area, rs11717303

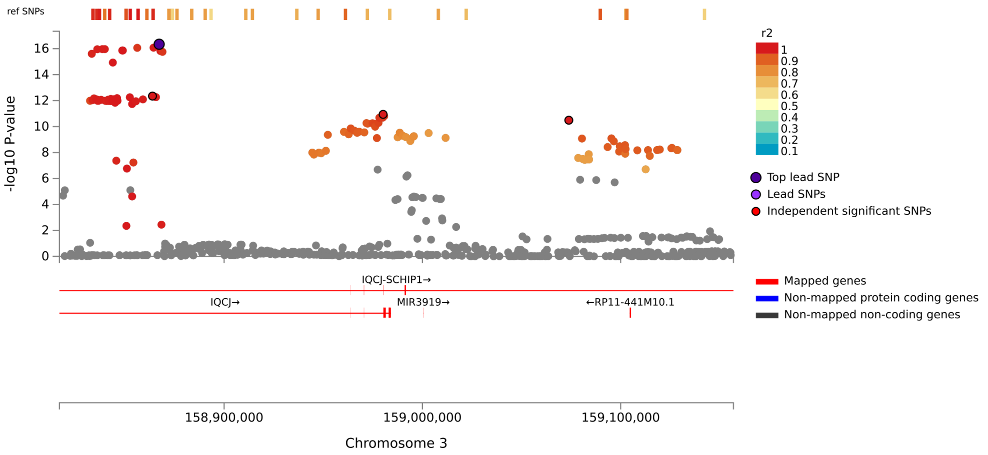


Locus 8, TNIK, Total Area, rs2035913

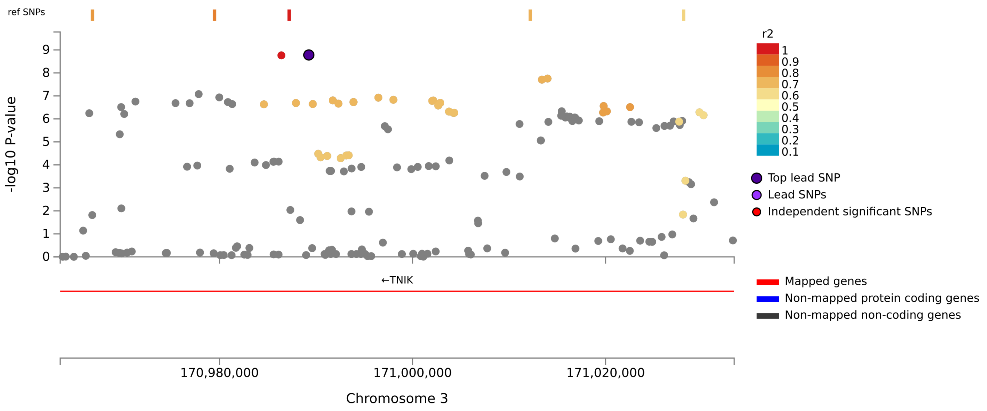

Locus 9, FIP1L1, Total Area, rs6835429

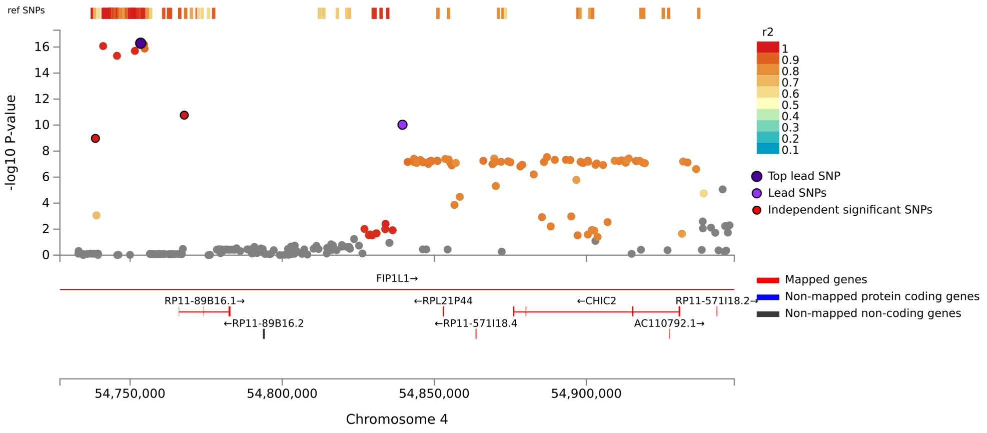


Locus 10, CTD-2316B1.1, Total Area, rs2919904

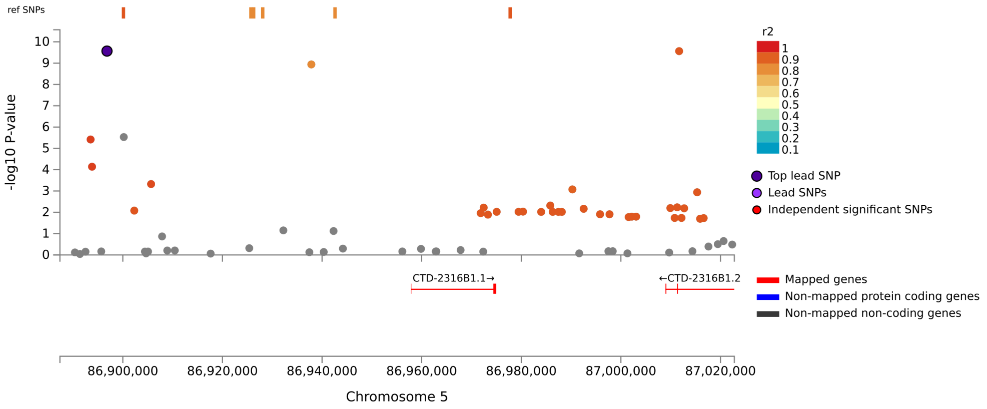

Locus 11, CTB-118N6.2, Total Area, rs17140192

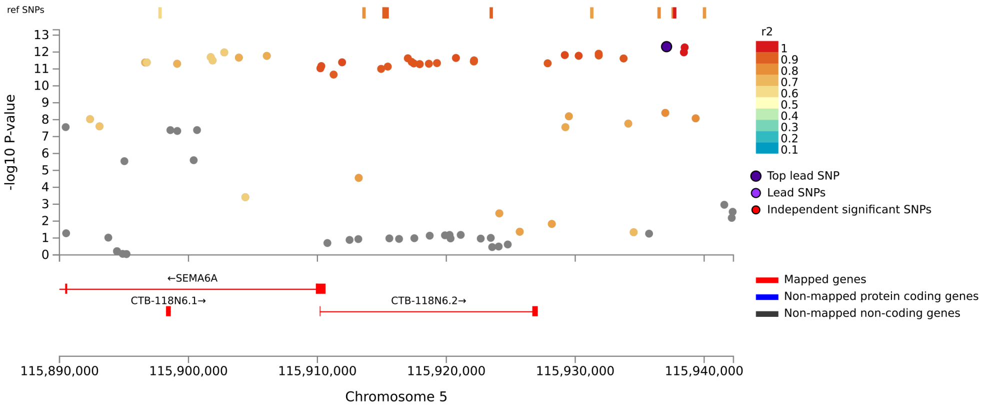


Locus 12, HBEGF, Total Area, rs4913081

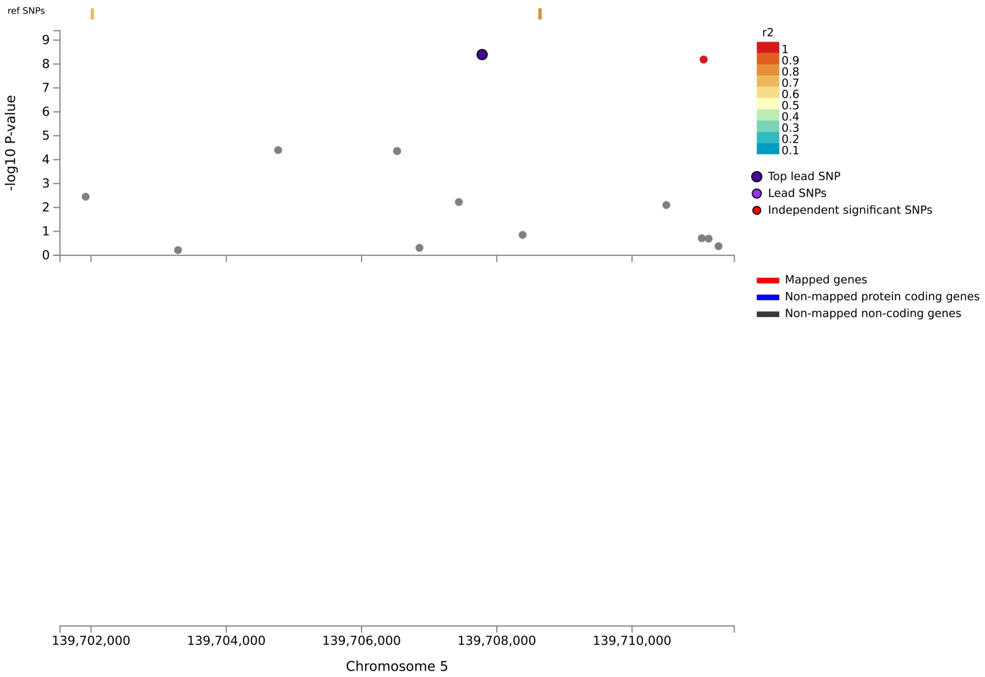

Locus 13, FOXO3, Total Area, rs9486902

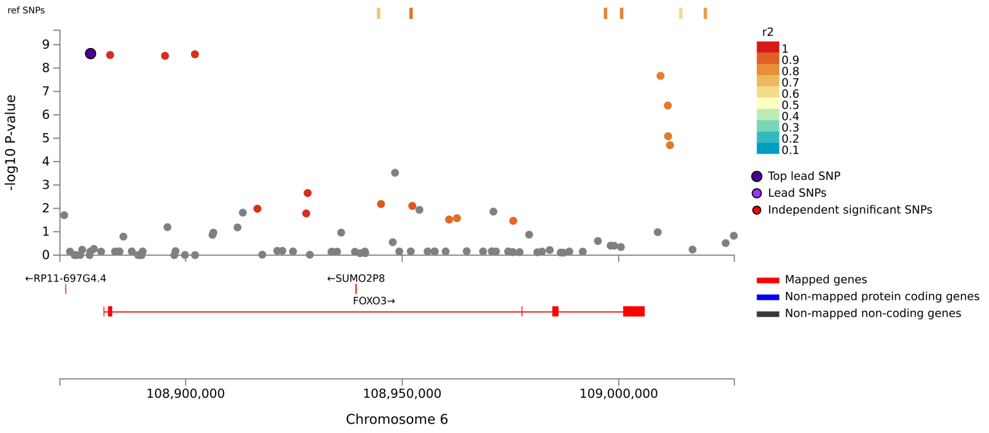


Locus 14, SNORA73, Total Area, rs76928645

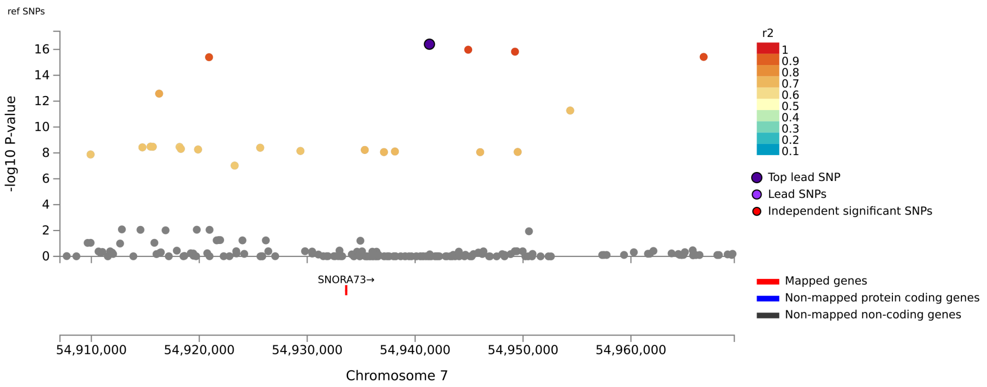

#### Locus 15, RP1-16A9.1, Total Area, rs2936679

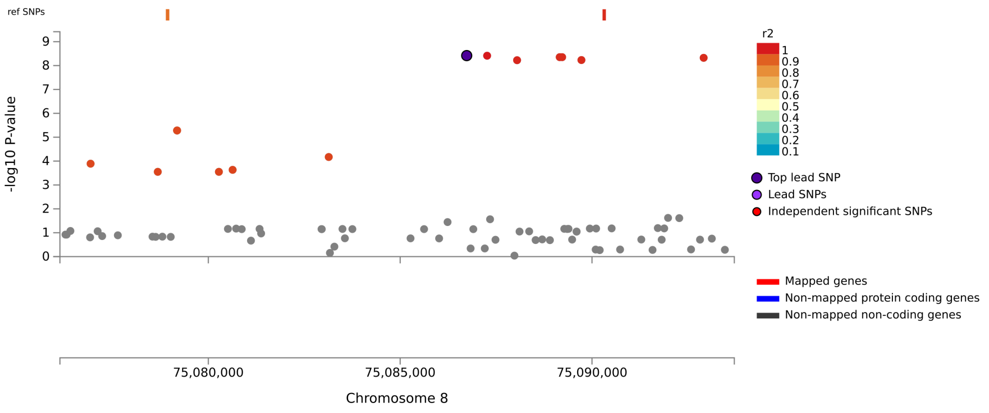


#### Locus 16, SLC45A4, Total Area, rs3739241

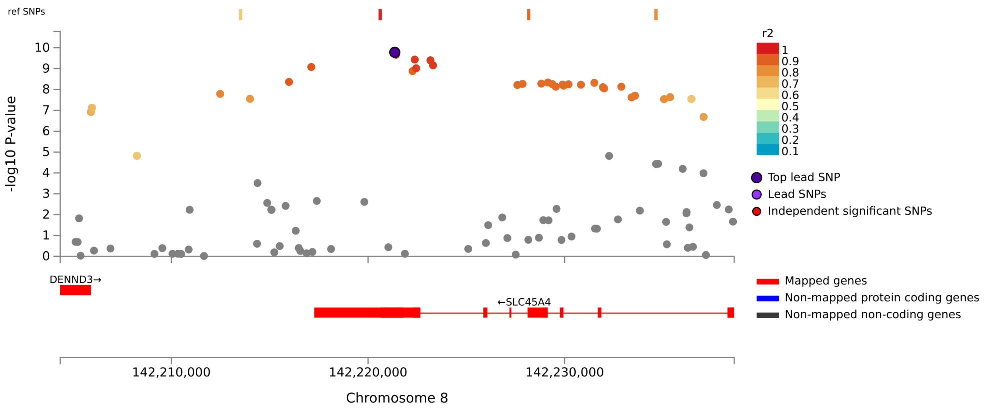

#### Locus 17, PLEC, Total Area, rs55646585

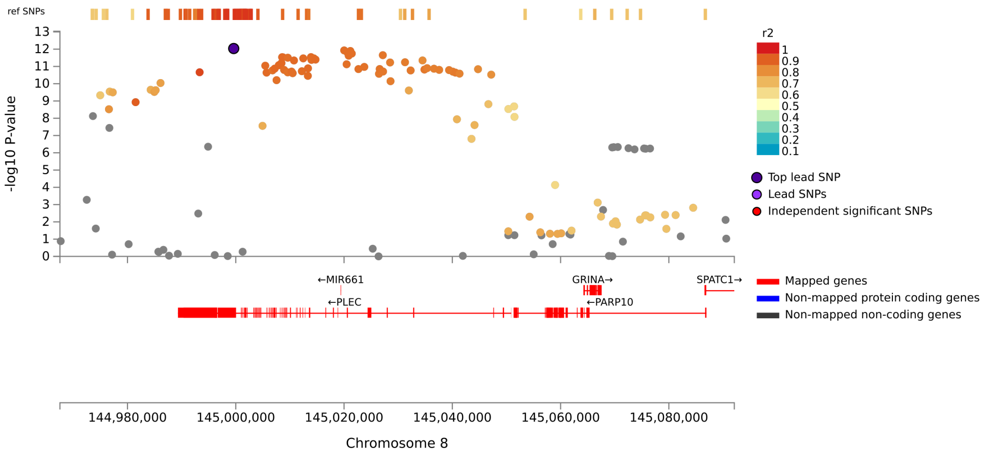


#### Locus 18, CDKN2B-AS1, Total Area, rs4451405

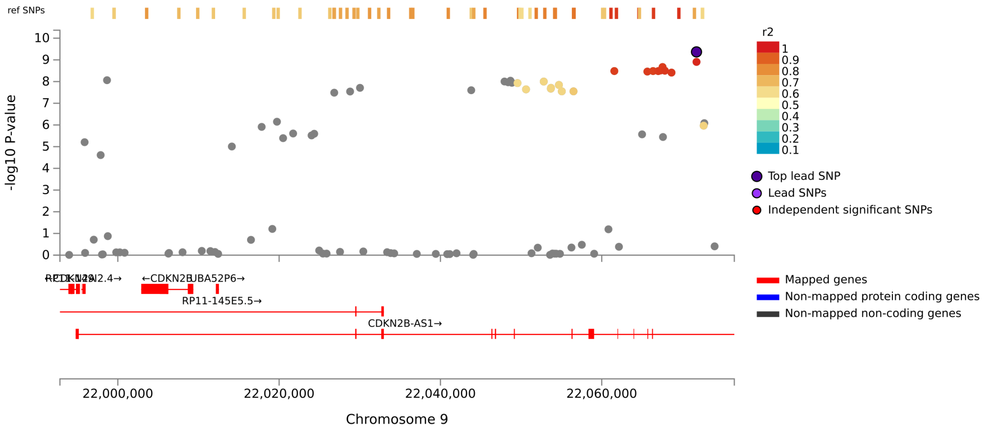

Locus 19, BICD2, Total Area, rs10992447

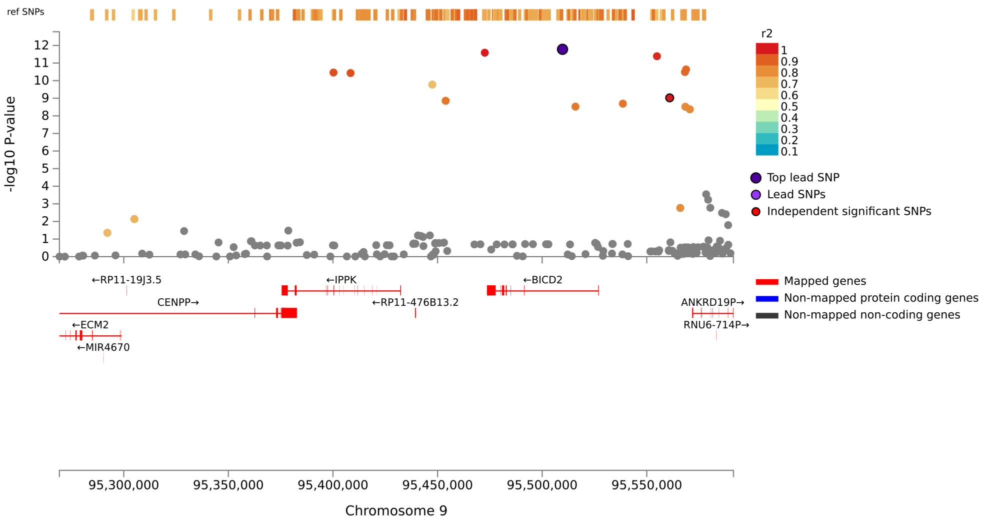


Locus 20, FAM107B, Total Area, rs1410071

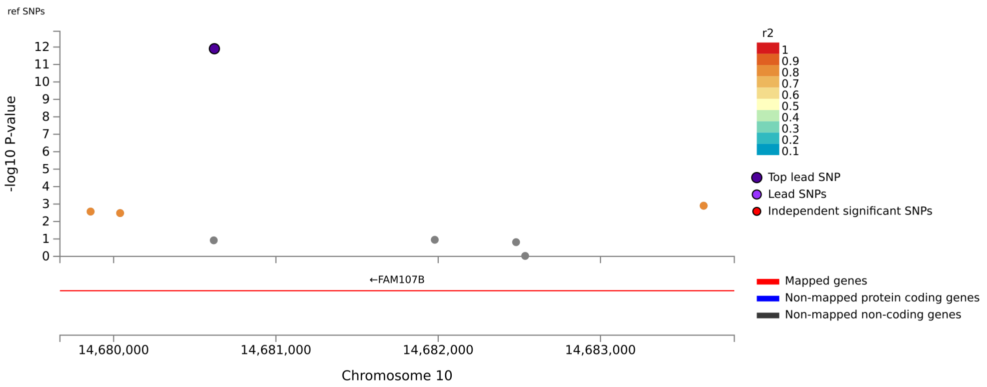

Locus 21, KIAA1598, Total Area, rs1122688

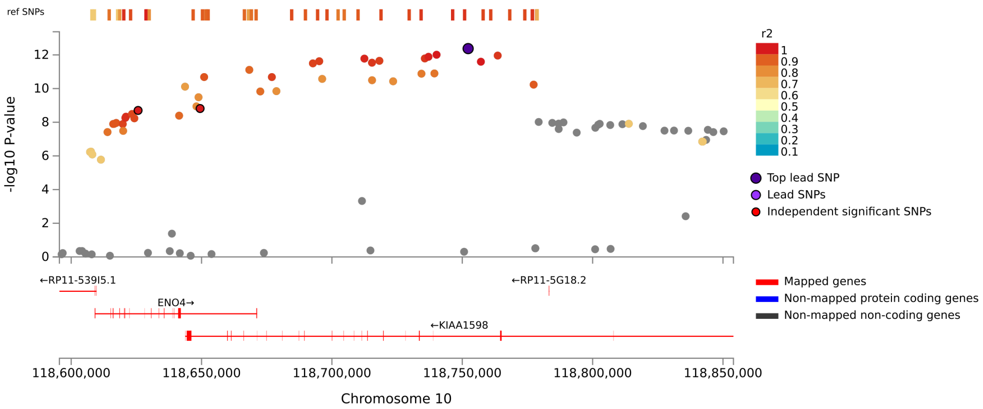


Locus 22, BRSK2, Total Area, rs7947308

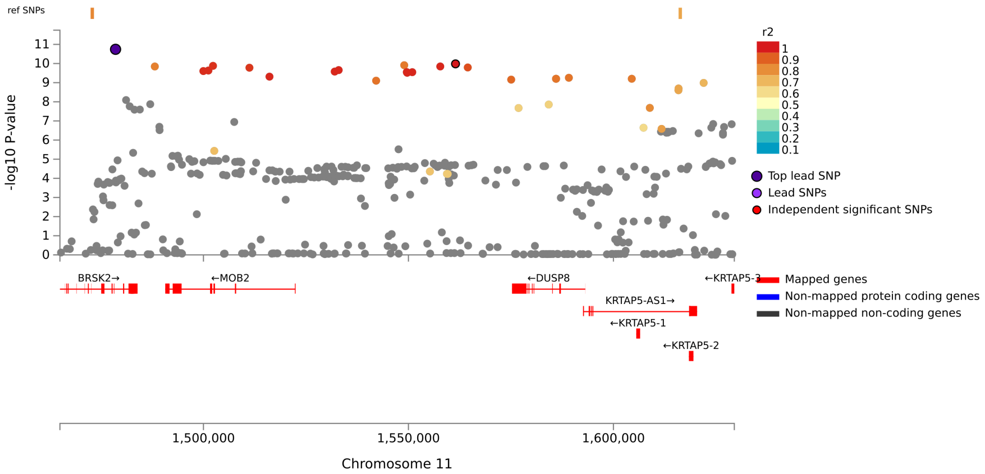

Locus 23, NAV2, Total Area, rs2625320

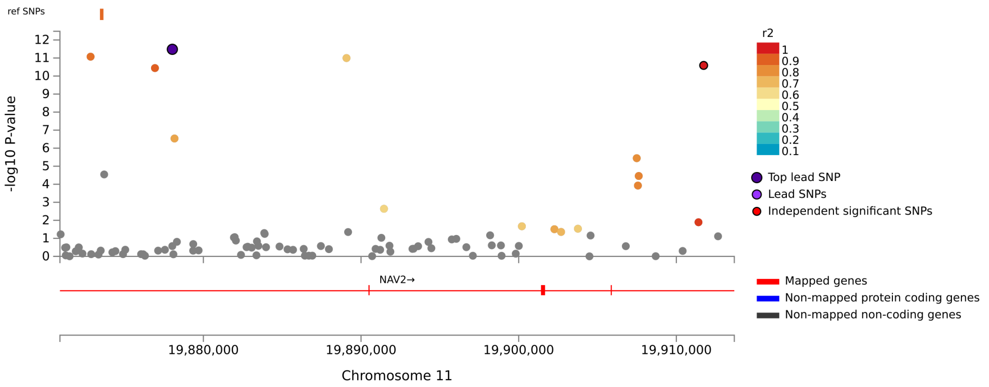


Locus 24, PPP2R5E, Total Area, rs11435884

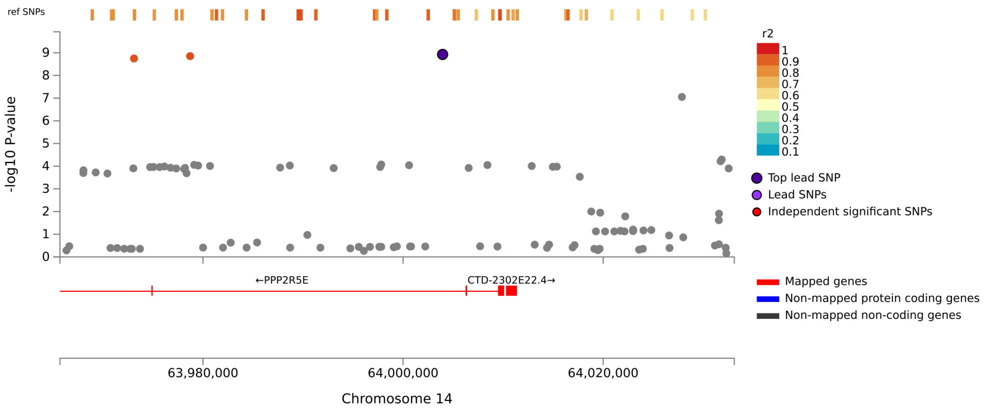

#### Locus 25, KPNA2, Total Area, rs62086903

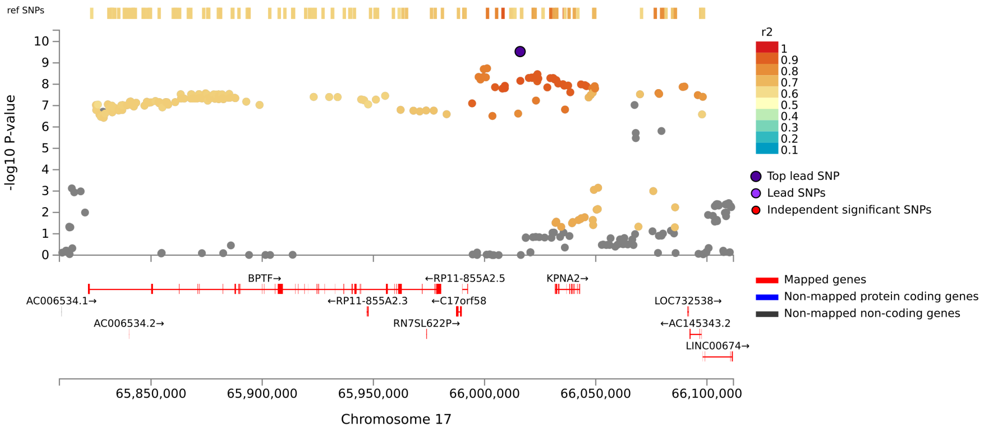


#### Locus 26, TTC39C:RP11-799B12.2, Total Area, rs8083625

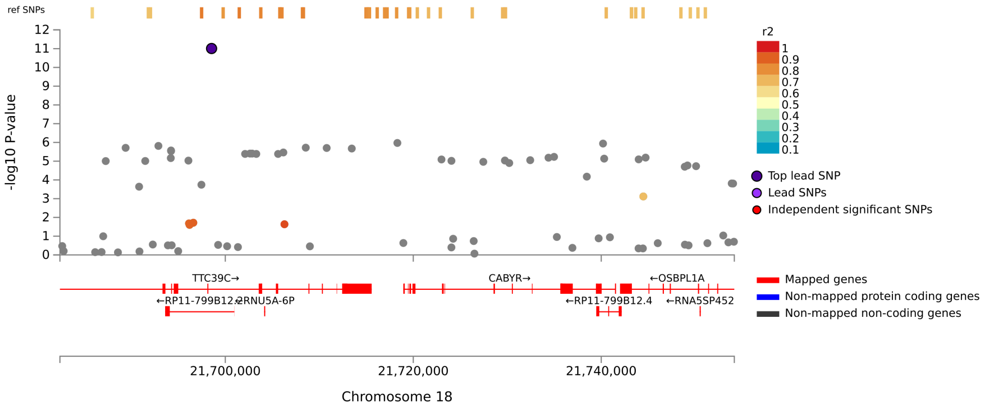

Locus 27, GAL3ST1, Total Area, rs2267158

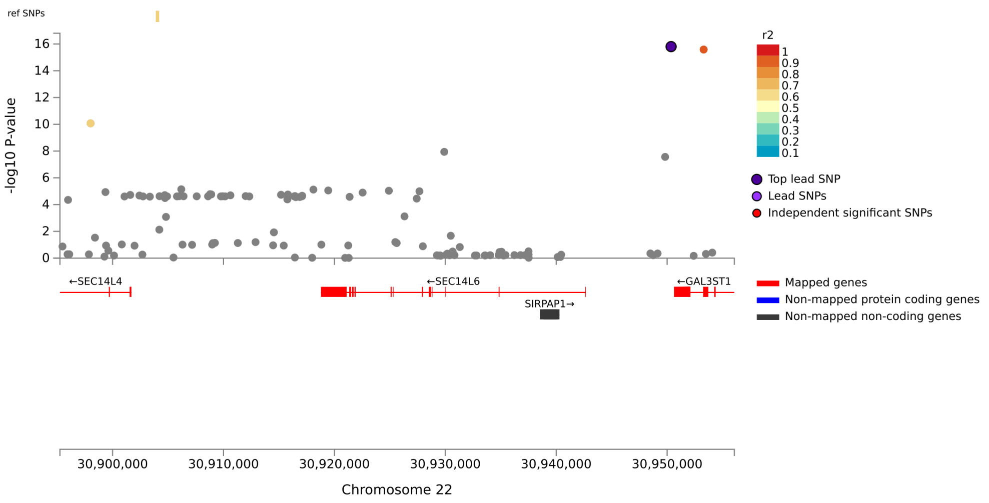


Locus 28, PICK1:RP5-1039K5.13, Total Area, rs760975

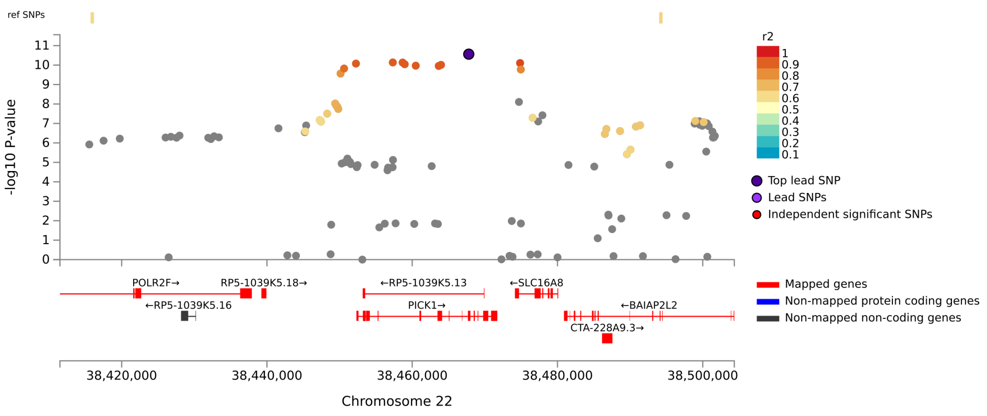

Locus 1, Y_RNA, Total Mean Thickness, rs9427220

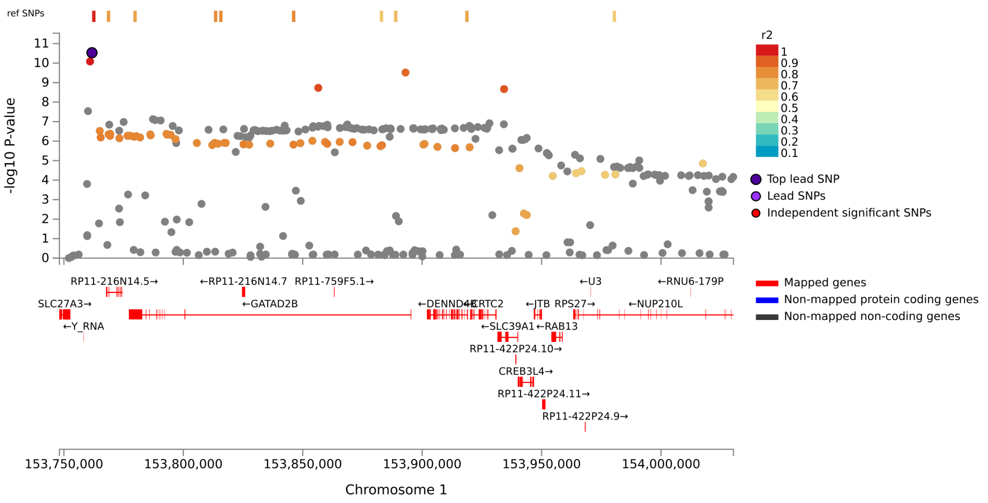


Locus 2, STRN, Total Mean Thickness, rs17496249

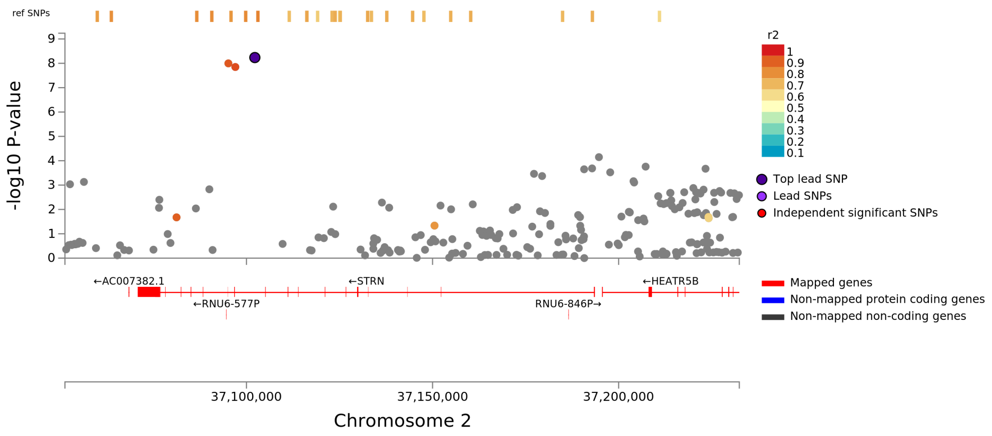

Locus 3, ITGAV:AC017101.10, Total Mean Thickness, rs7589470

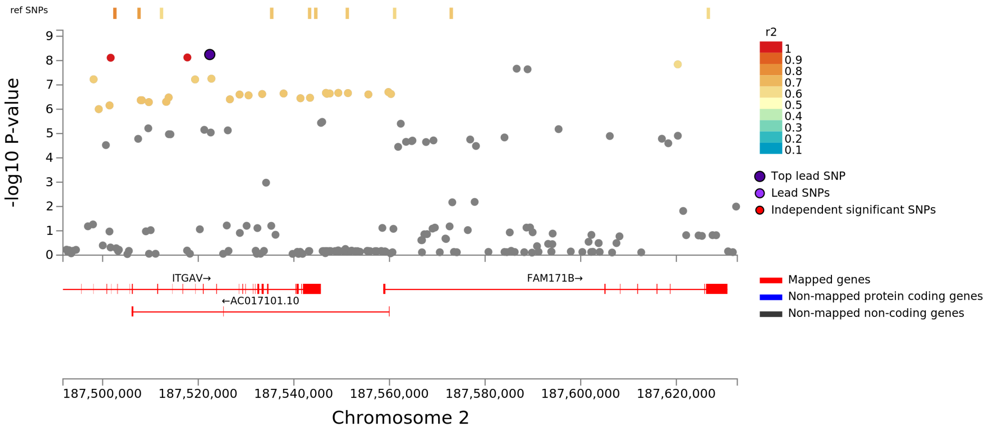


Locus 4, IQCJ-SCHIP1:IQCJ, Total Mean Thickness, rs12632564

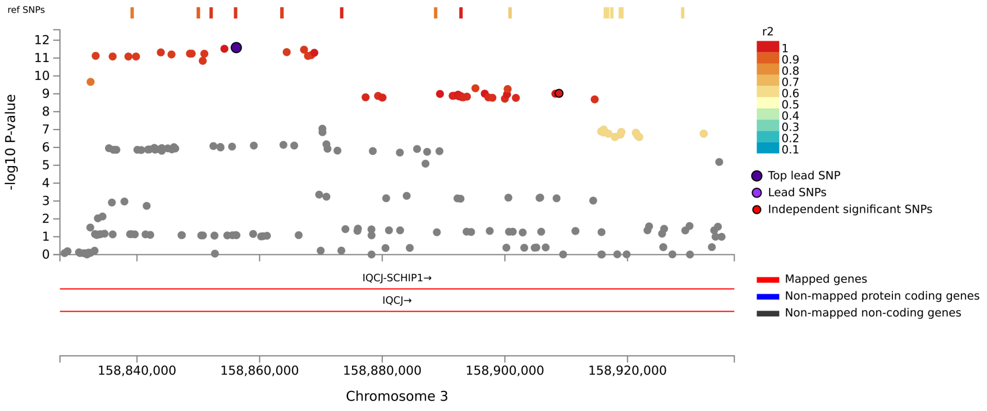

#### Locus 5, FIP1L1, Total Mean Thickness, rs6849897

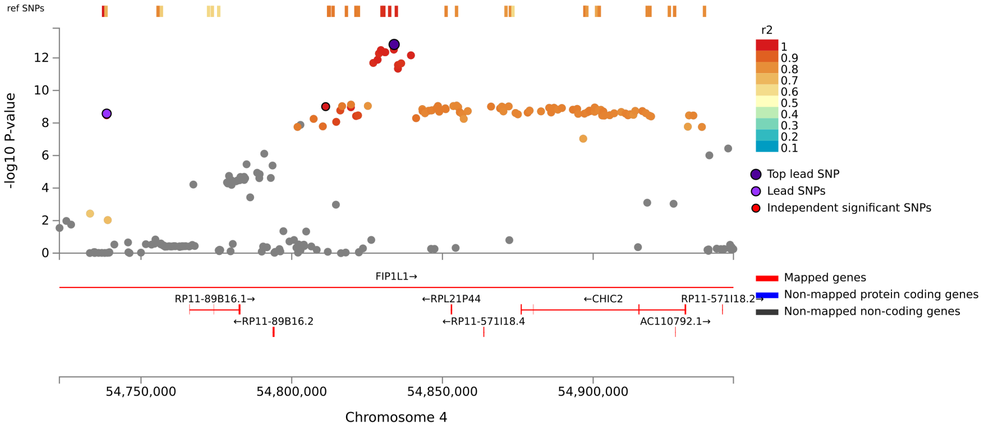


#### Locus 6, HBEGF, Total Mean Thickness, rs4150211

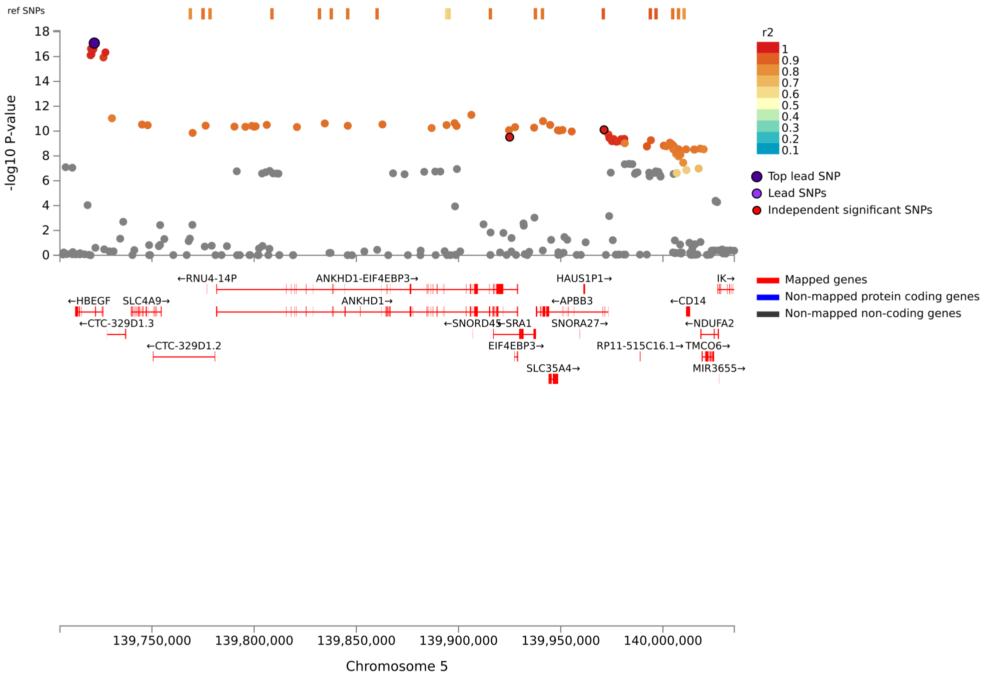

Locus 7, PARP10, Total Mean Thickness, rs62530285

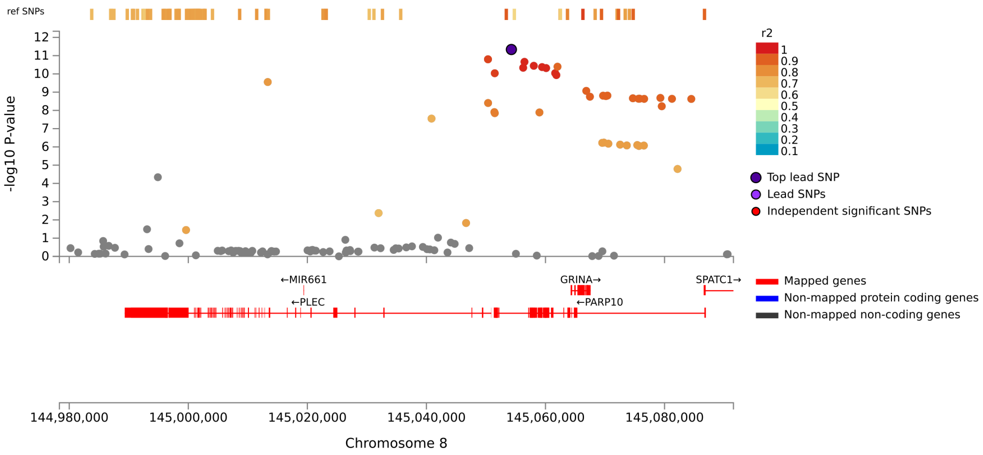


Locus 8, CDKN2B-AS1, Total Mean Thickness, rs1412832

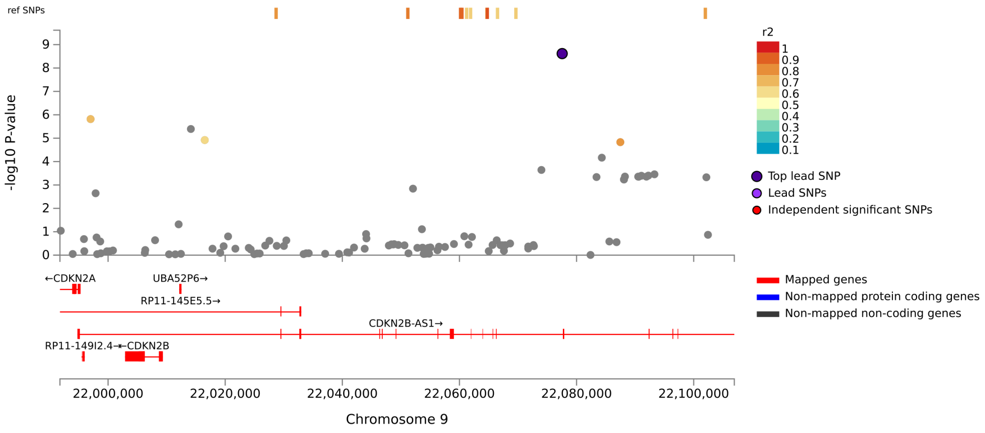

Locus 9, FAM107B, Total Mean Thickness, rs878733

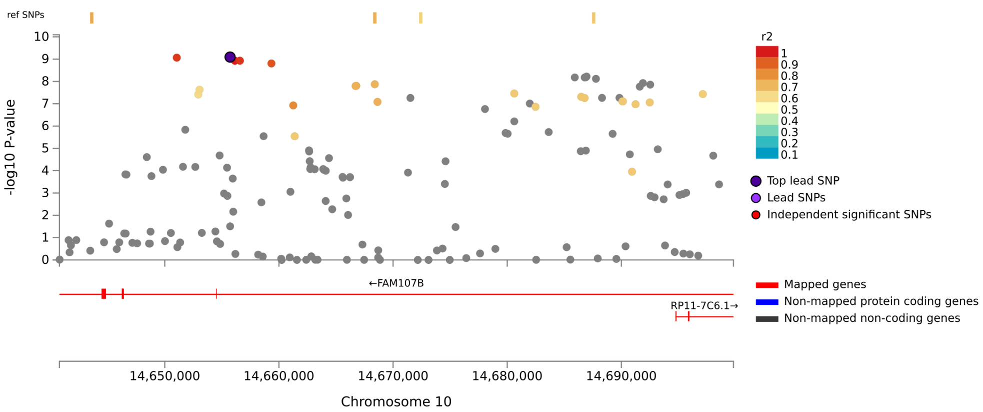


Locus 10, C16orf95, Total Mean Thickness, rs4843550

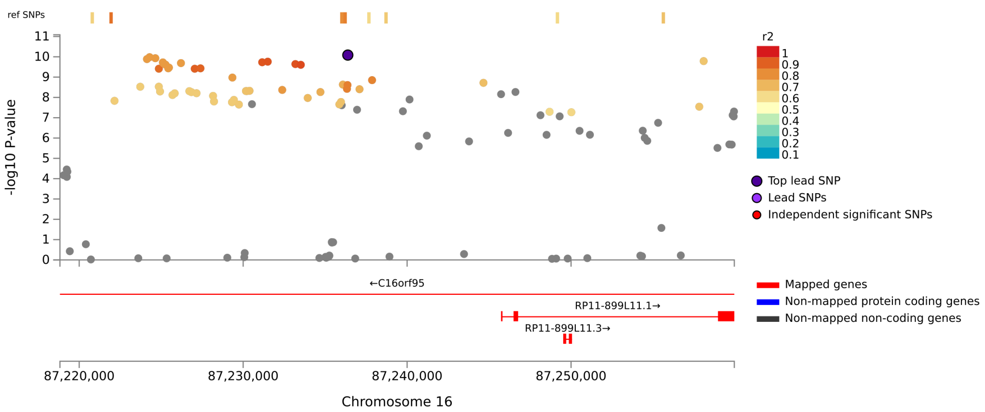

### Locus 11, CABYR, Total Mean Thickness, rs752797

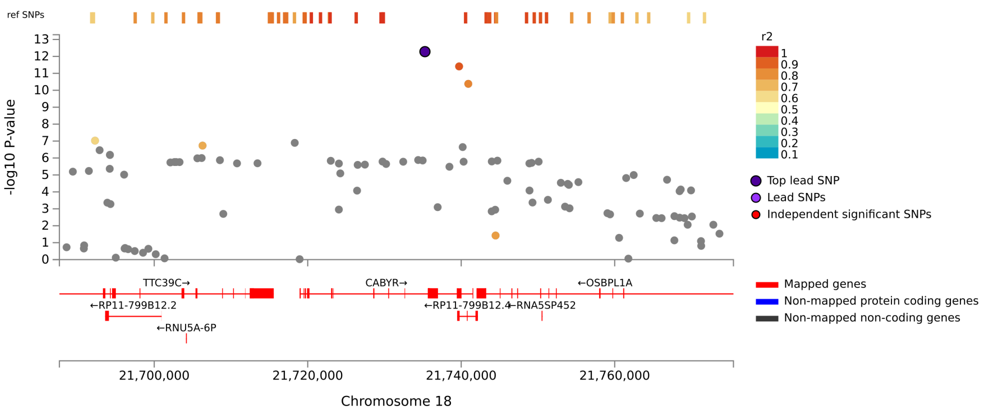

Locus 1, ZZZ3, Genu Area, rs12026939

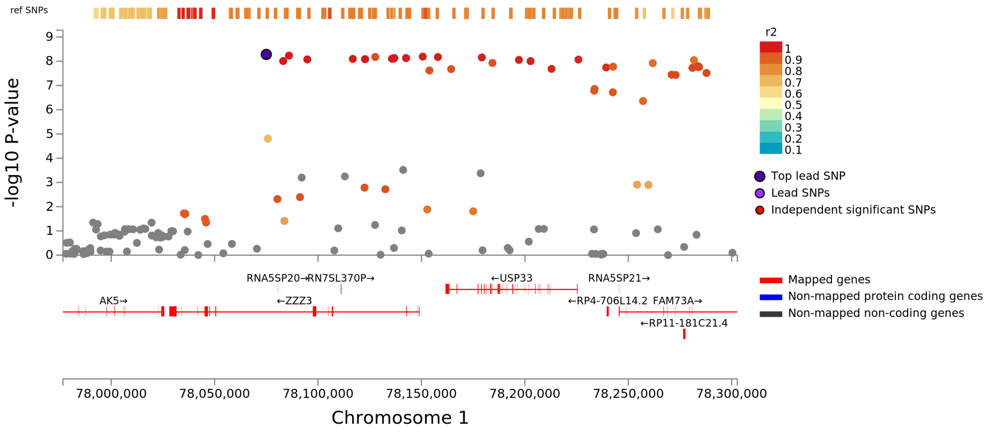


Locus 2, STX6, Genu Area, rs35306826

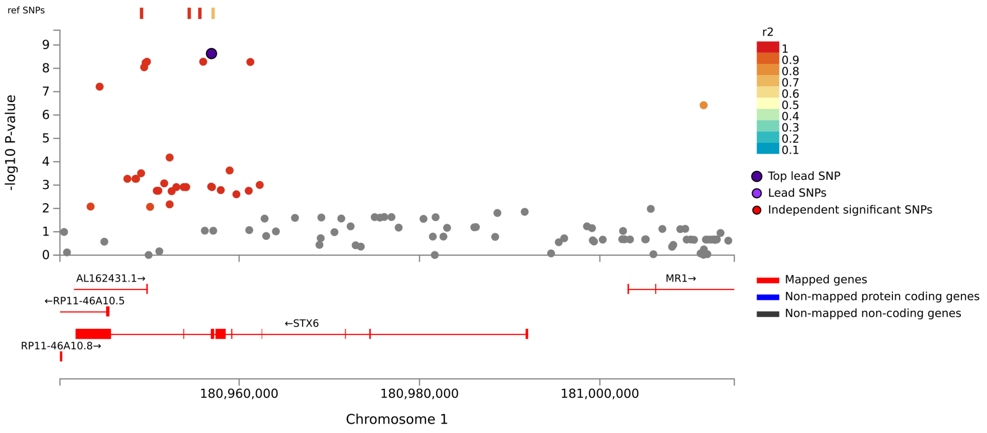

Locus 3, STRN, Genu Area, rs62132550

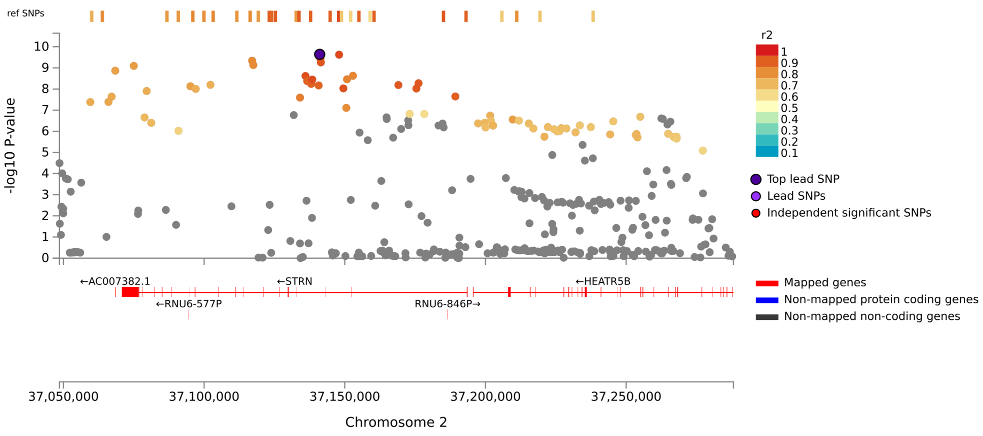


Locus 4, AC016727.1, Genu Area, rs778766

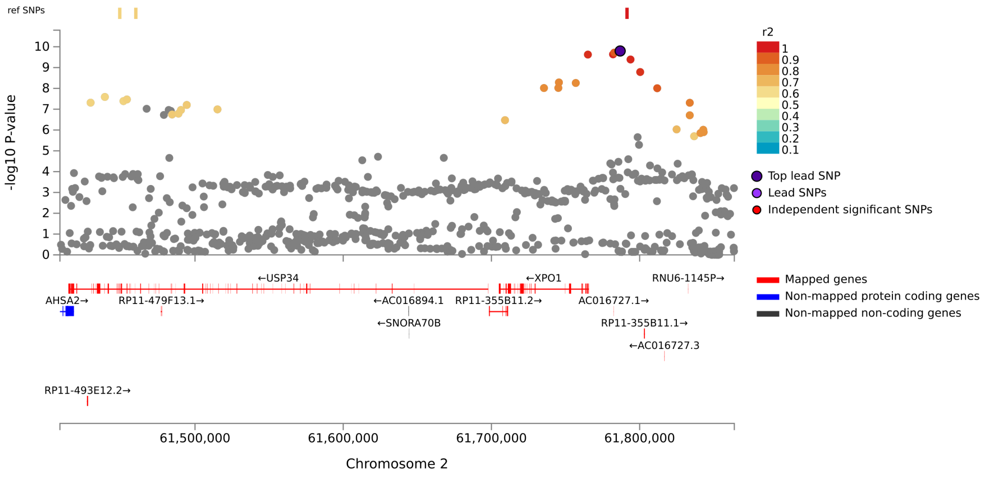

Locus 5, FAM171B, Genu Area, rs3898135

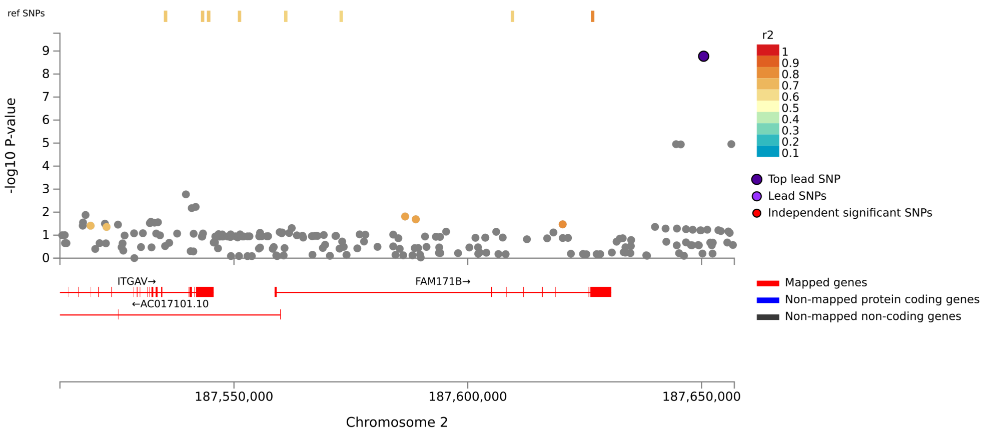


Locus 6, RPL37A, Genu Area, rs62180637

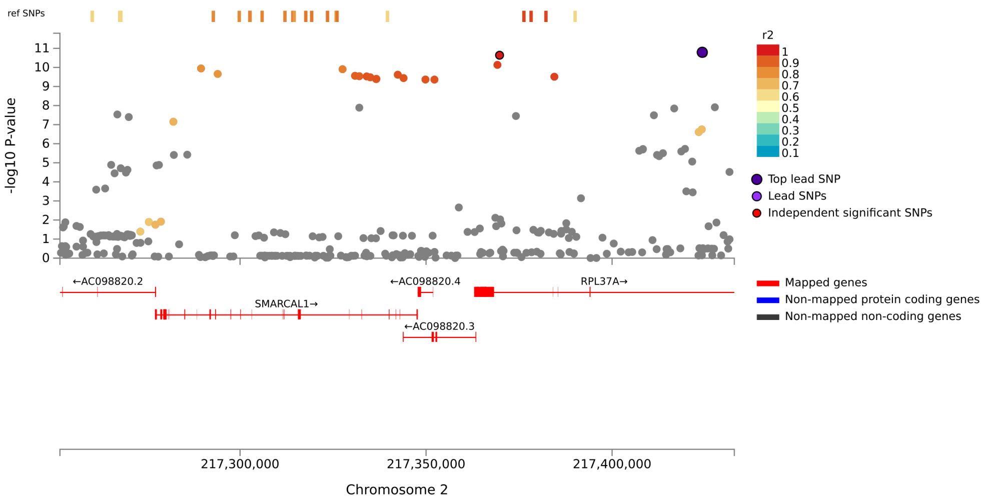

#### Locus 7, IQCJ-SCHIP1:IQCJ, Genu Area, rs12632564

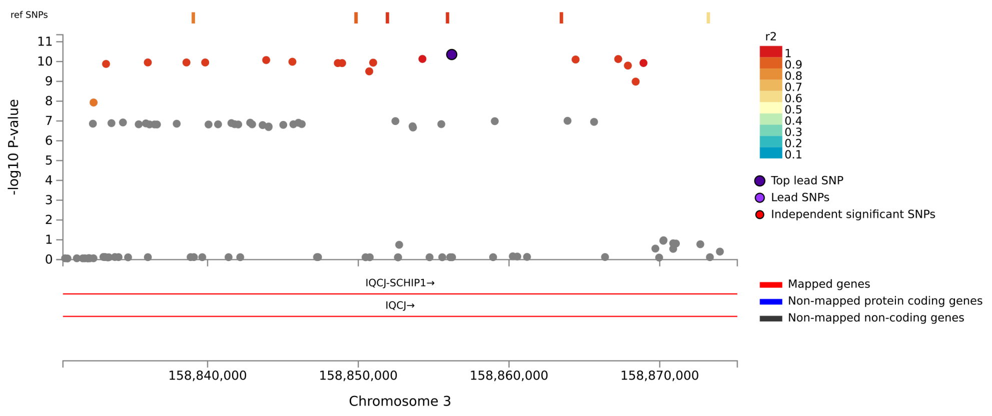


#### Locus 8, CTC-448D22.1, Genu Area, rs721065

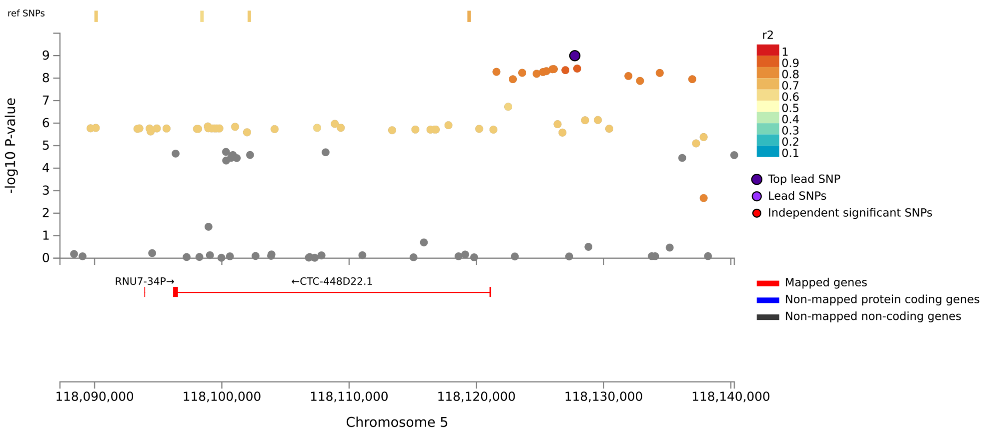

#### Locus 9, SNORA73, Genu Area, rs76928645

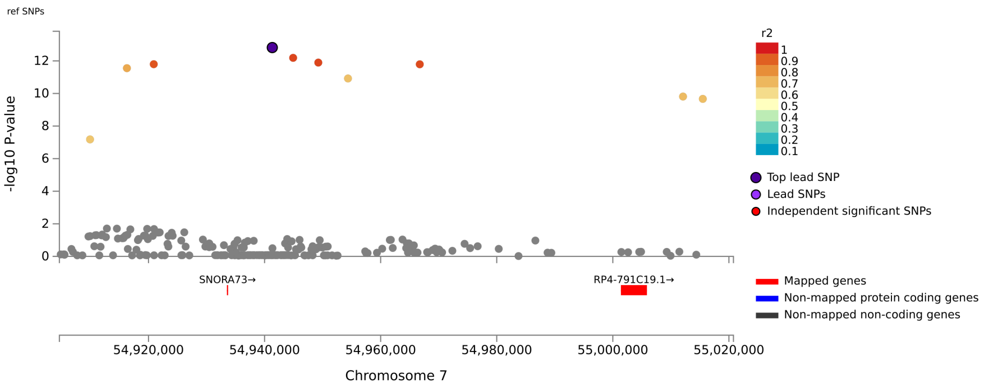


#### Locus 10, AC003084.2, Genu Area, rs7784849

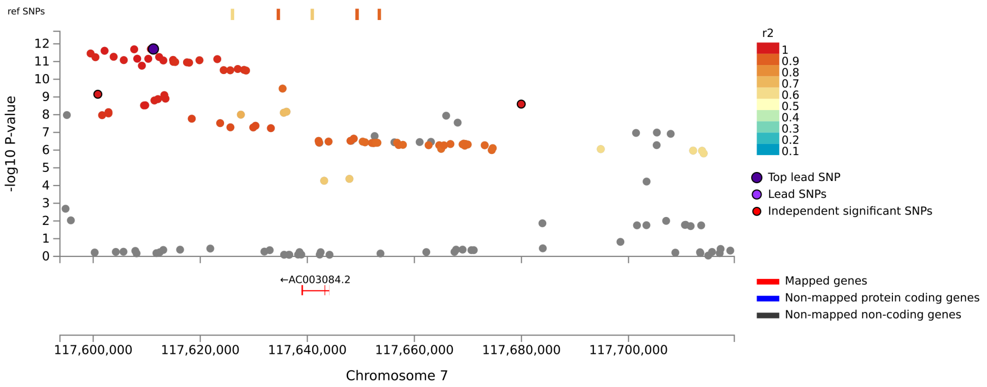

#### Locus 11, PLEC, Genu Area, rs56401356

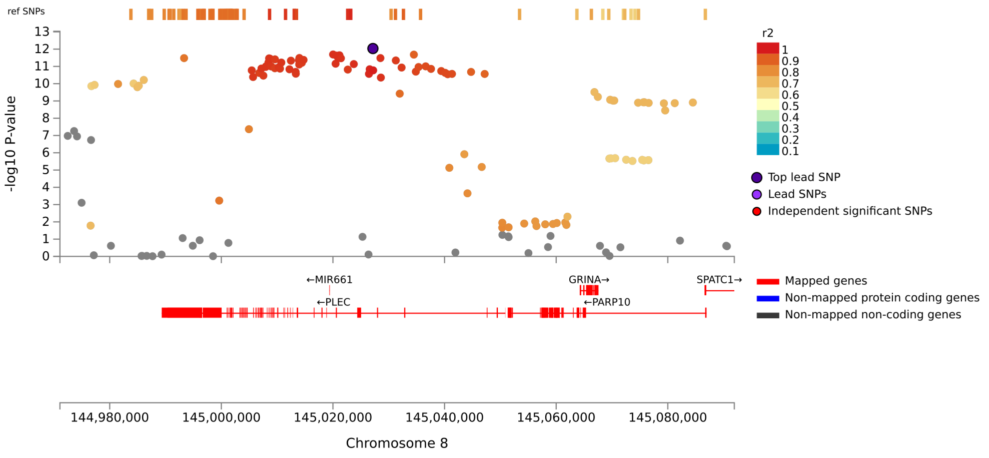


#### Locus 12, CDKN2B-AS1, Genu Area, rs1412832

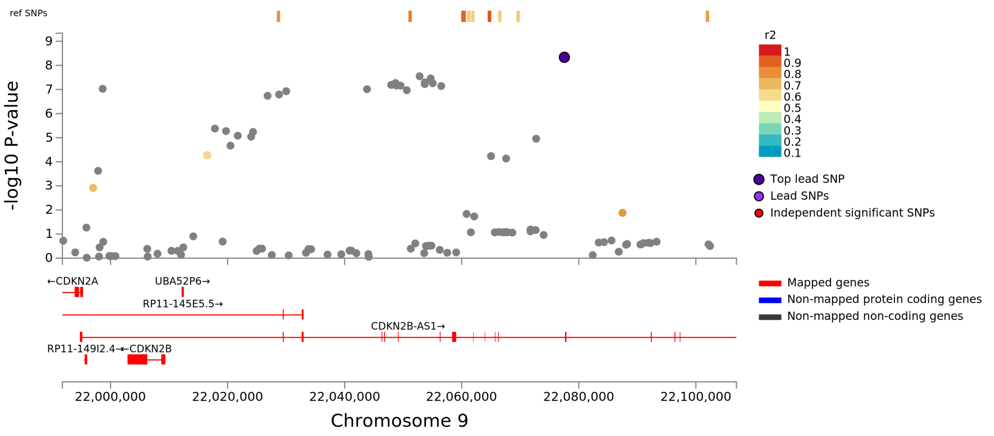

Locus 13, KIAA1598, Genu Area, rs1122688

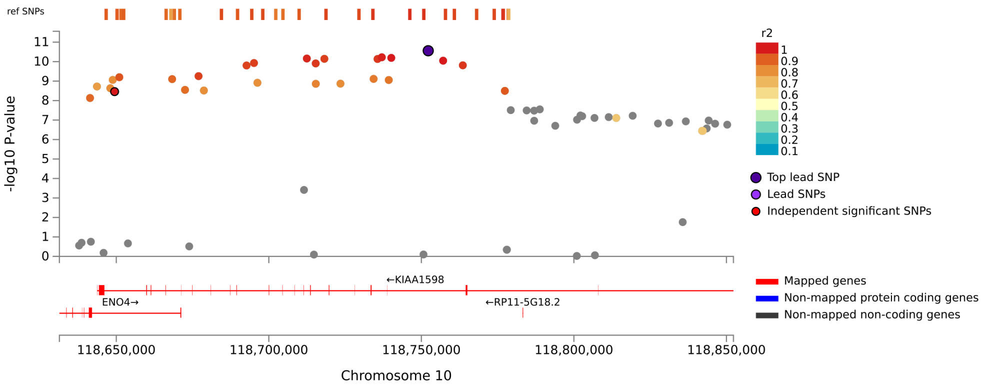


Locus 14, NAV2, Genu Area, rs2625302

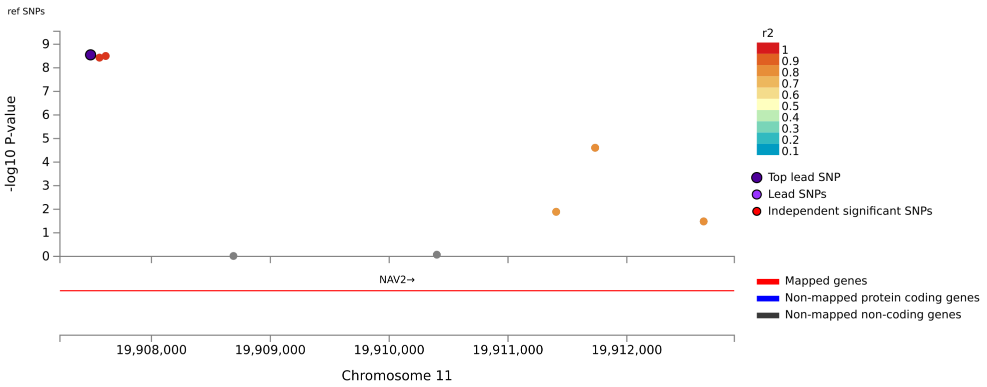

Locus 15, CELF1, Genu Area, rs11039266

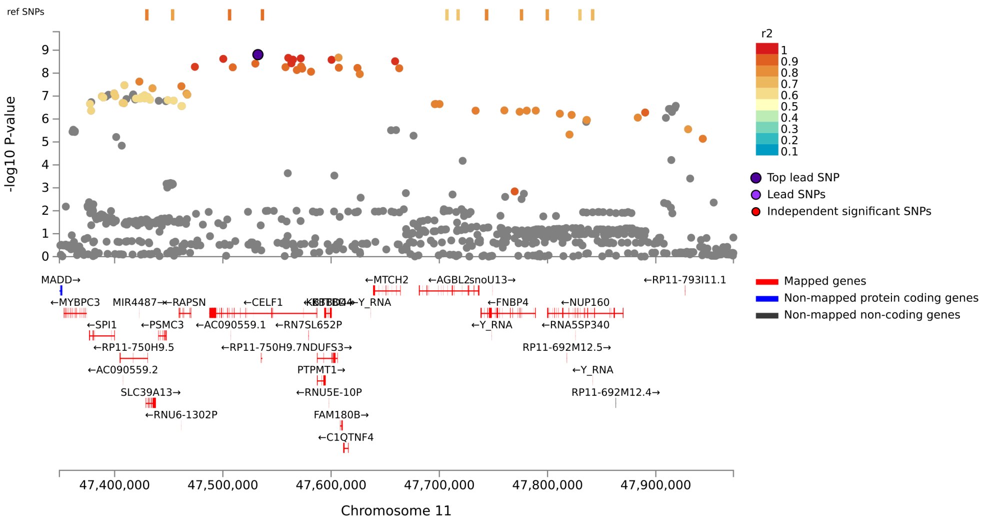


Locus 16, NAV3, Genu Area, rs10506772

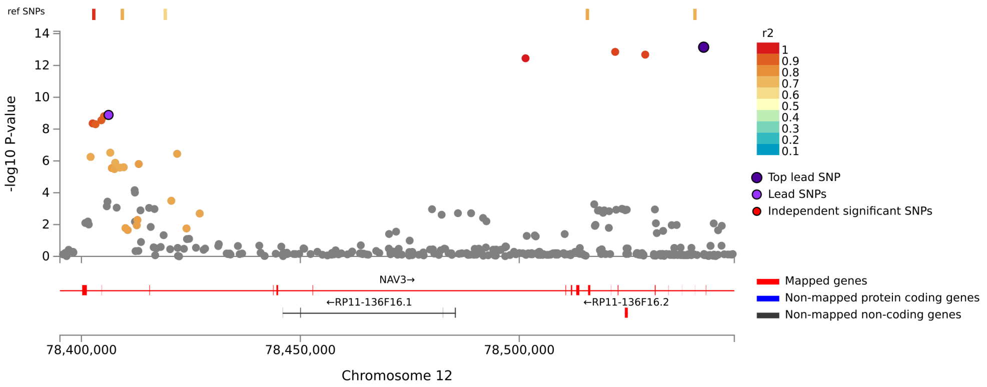

Locus 17, KANSL1, Genu Area, rs2532402

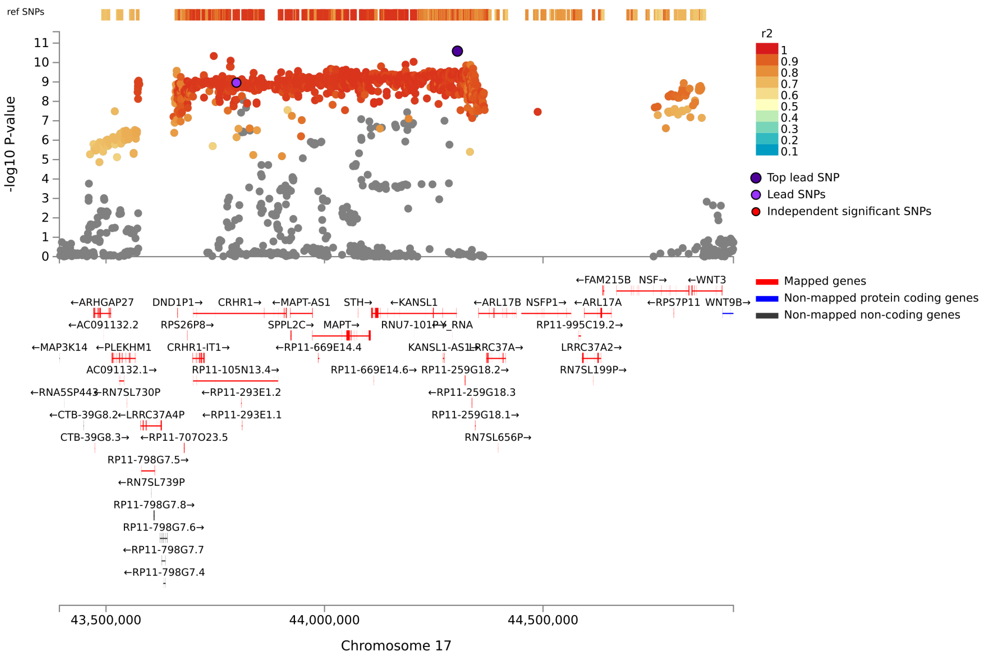


Locus 18, GAL3ST1, Genu Area, rs2267161

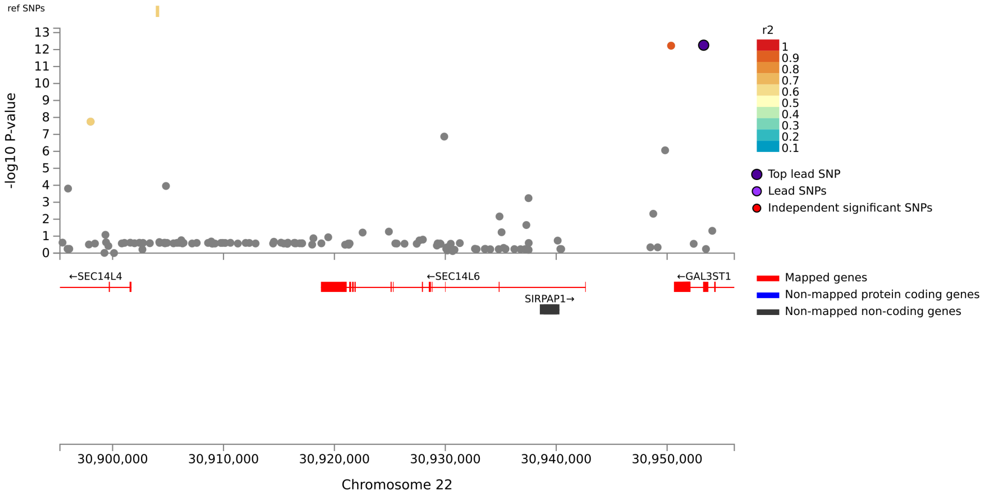

### Locus 19, PICK1:RP5-1039K5.13, Genu Area, rs738443

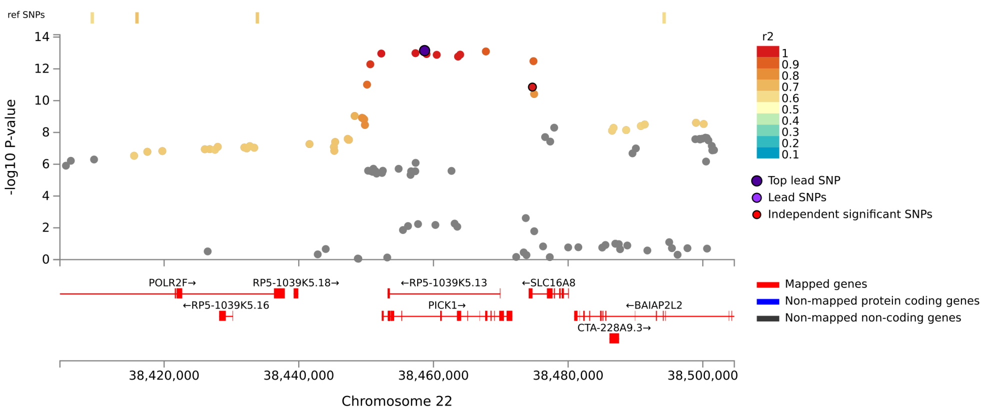

Locus 1, MAST4, Genu Mean Thickness, rs147629510

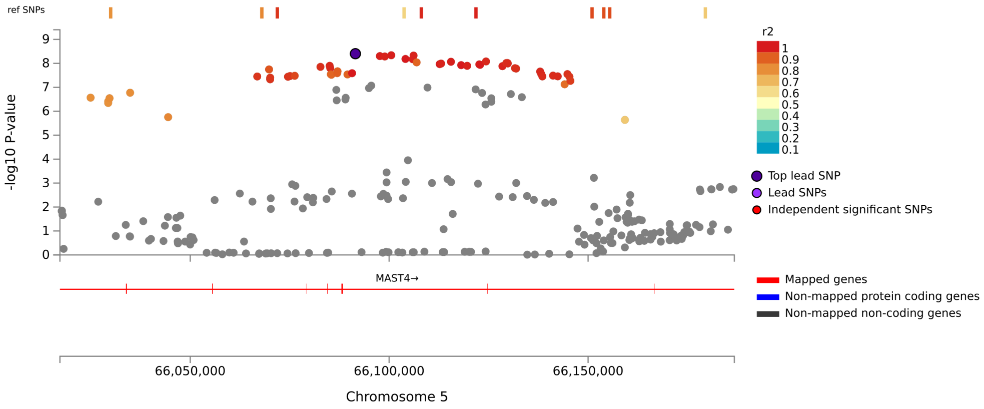


Locus 2, AC003084.2, Genu Mean Thickness, rs916784

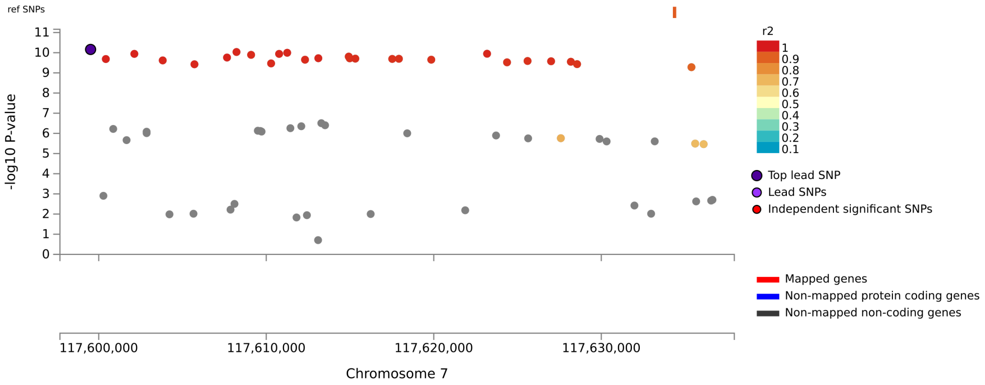

Locus 3, CHMP7, Genu Mean Thickness, rs7459962

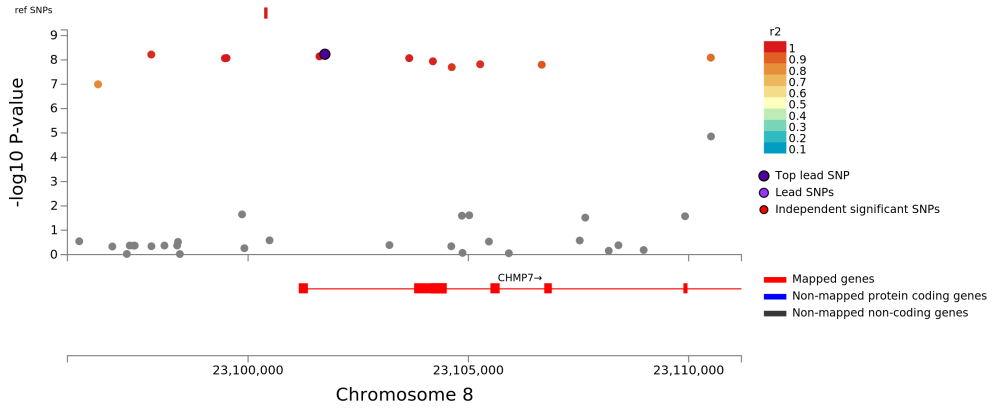

Locus 1, SDHB, Anterior Body Area, rs3754507

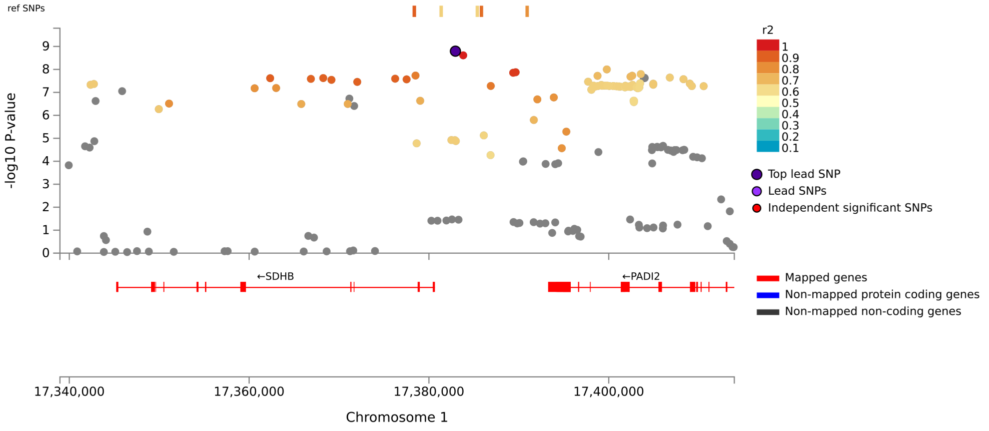


Locus 2, SOAT1, Anterior Body Area, rs3753526

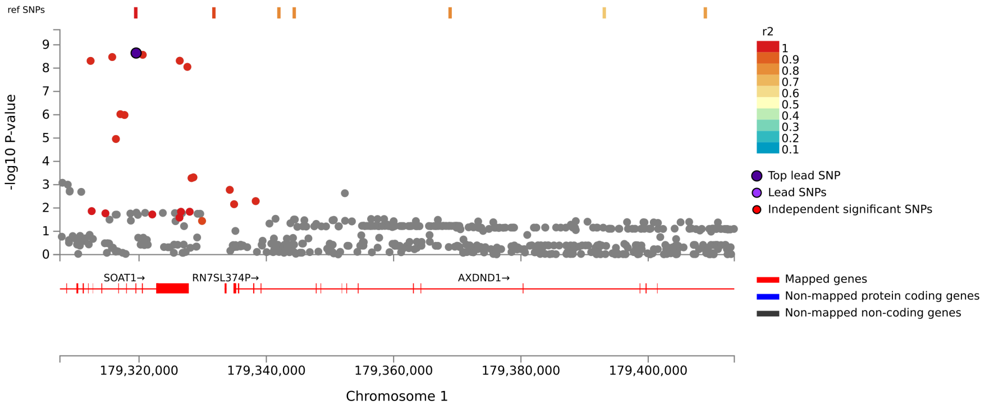

#### Locus 3, STRN, Anterior Body Area, rs7561572

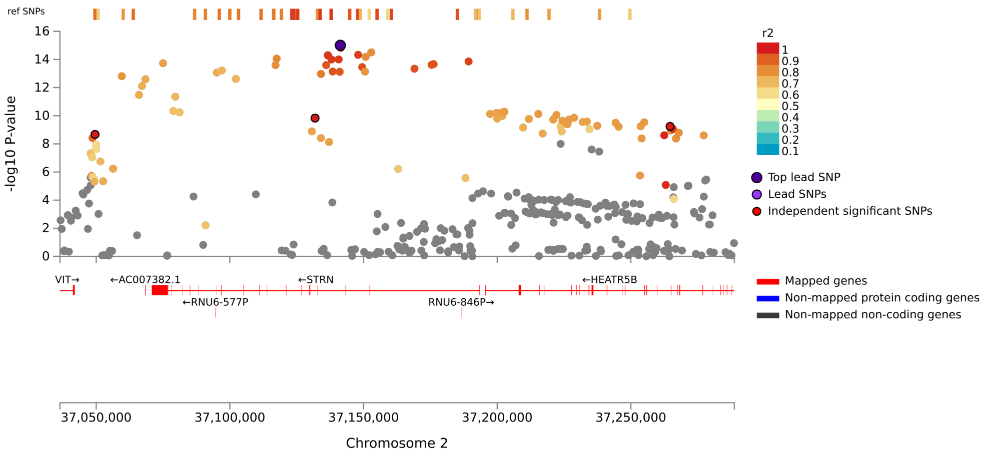


#### Locus 4, FIP1L1, Anterior Body Area, rs1466831

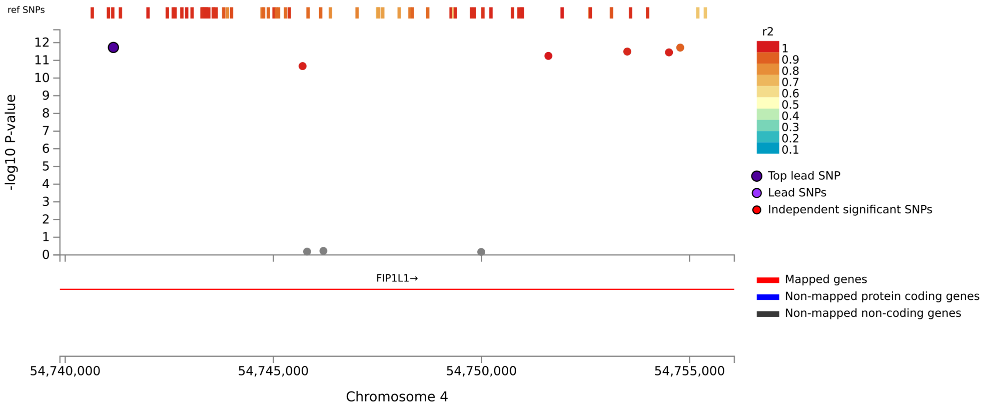

Locus 5, RNU6-727P, Anterior Body Area, rs71637276

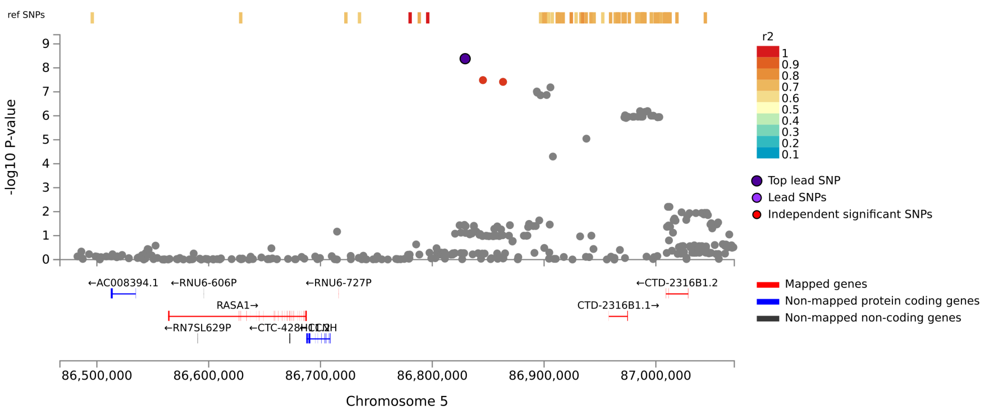


Locus 6, FOXO3, Anterior Body Area, rs1268163

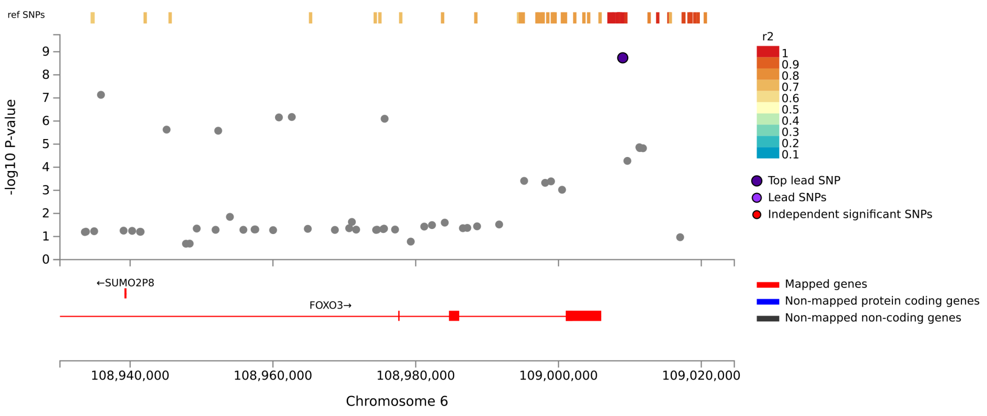

Locus 7, SNORA73, Anterior Body Area, rs74504435

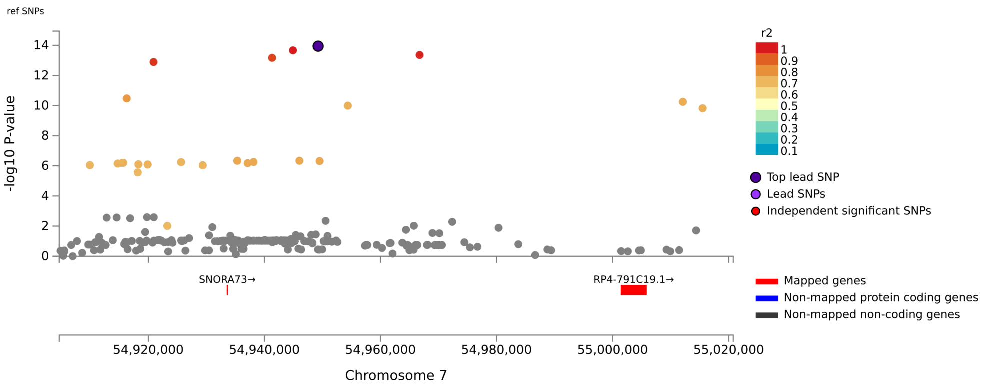


Locus 8, CPED1, Anterior Body Area, rs17356657

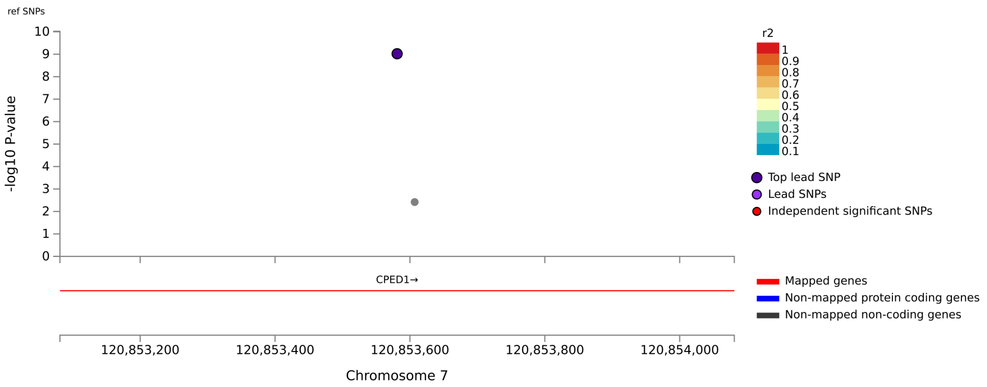

Locus 9, PLEC, Anterior Body Area, rs55646585

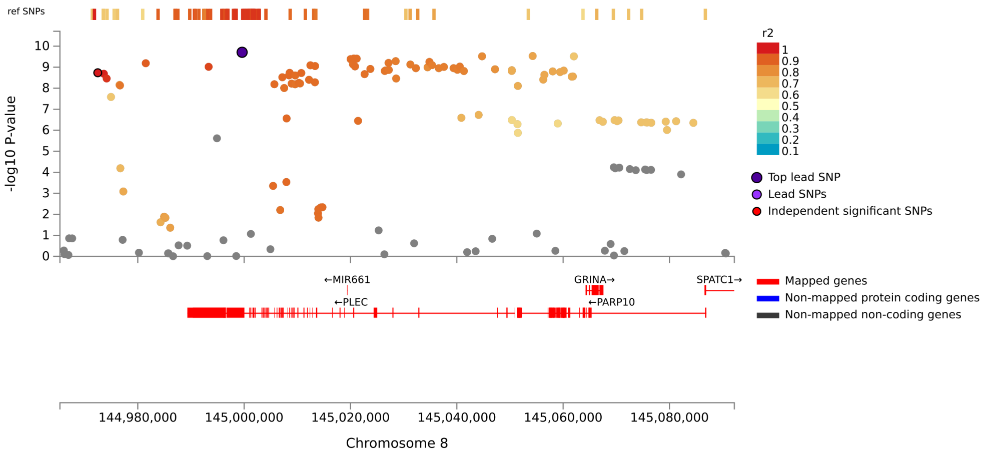


Locus 10, FAM107B, Anterior Body Area, rs10906729

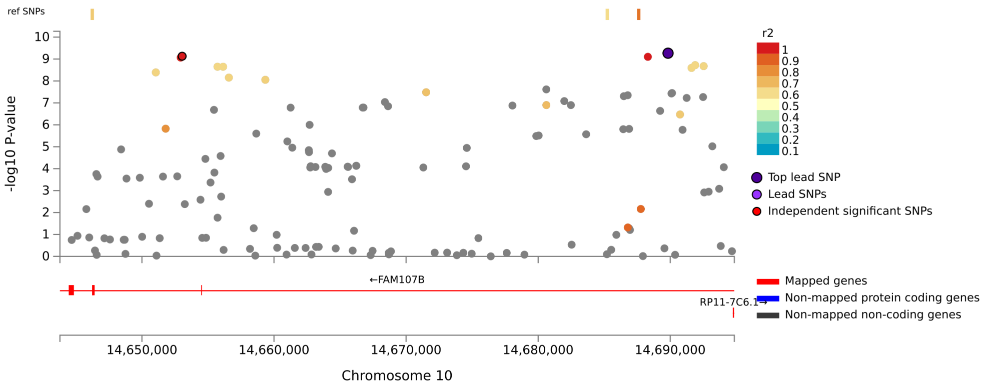

#### Locus 11, KIAA1598, Anterior Body Area, rs1122688

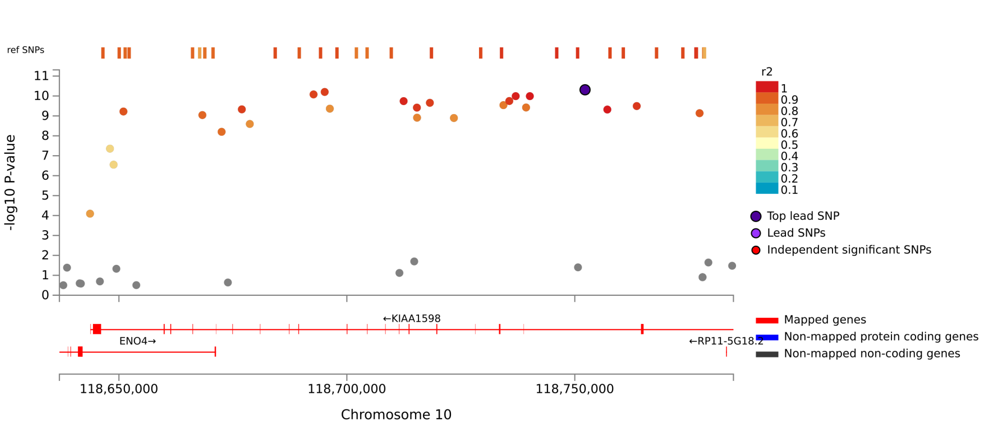


#### Locus 12, DUSP8, Anterior Body Area, rs7127282

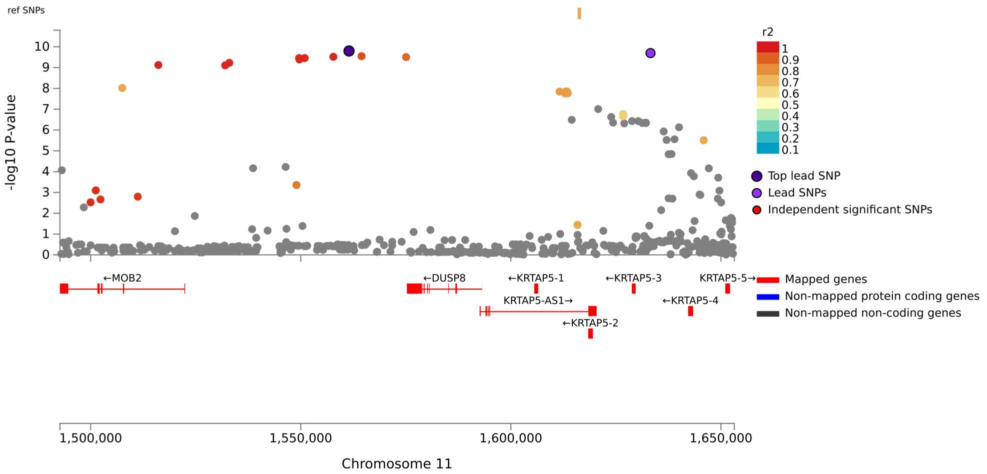

Locus 13, RP1-34H18.1, Anterior Body Area, rs696451

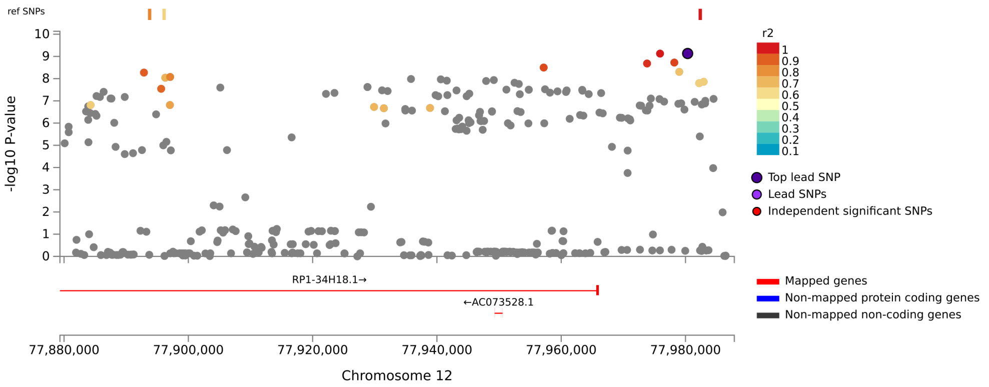


Locus 14, CABYR, Anterior Body Area, rs752797

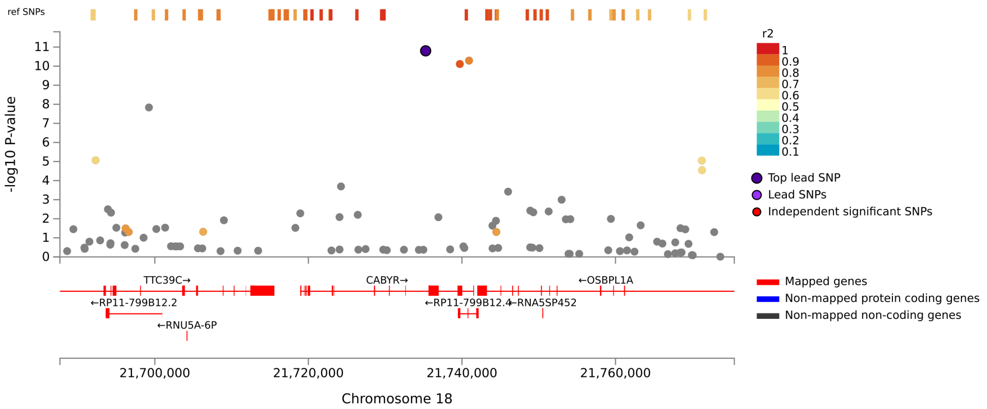

Locus 1, STRN, Anterior Body Mean Thickness, rs7561572

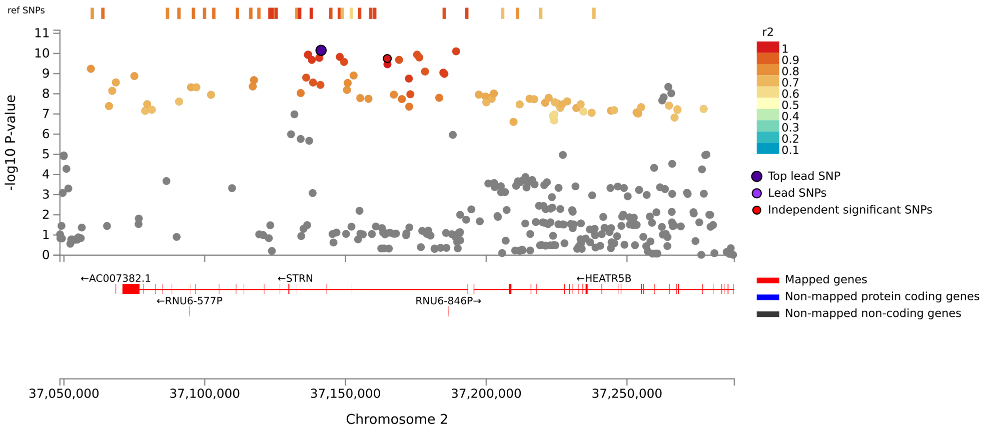


Locus 2, FIP1L1, Anterior Body Mean Thickness, rs1466831

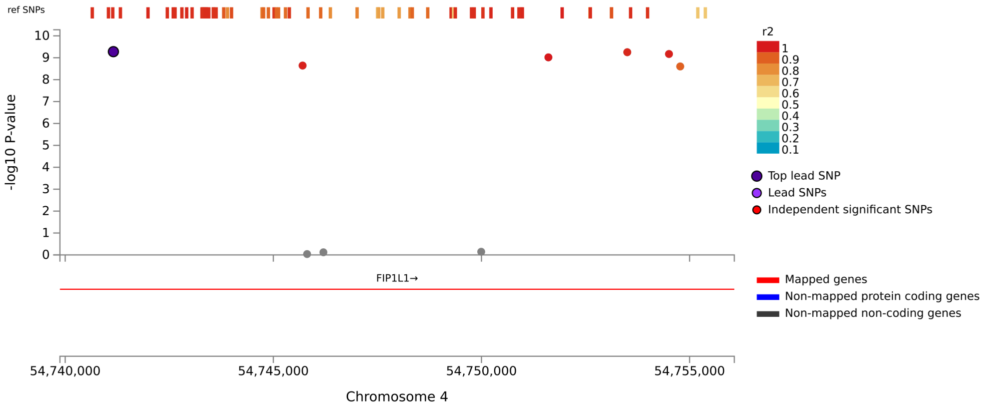

Locus 3, HBEGF, Anterior Body Mean Thickness, rs4150212

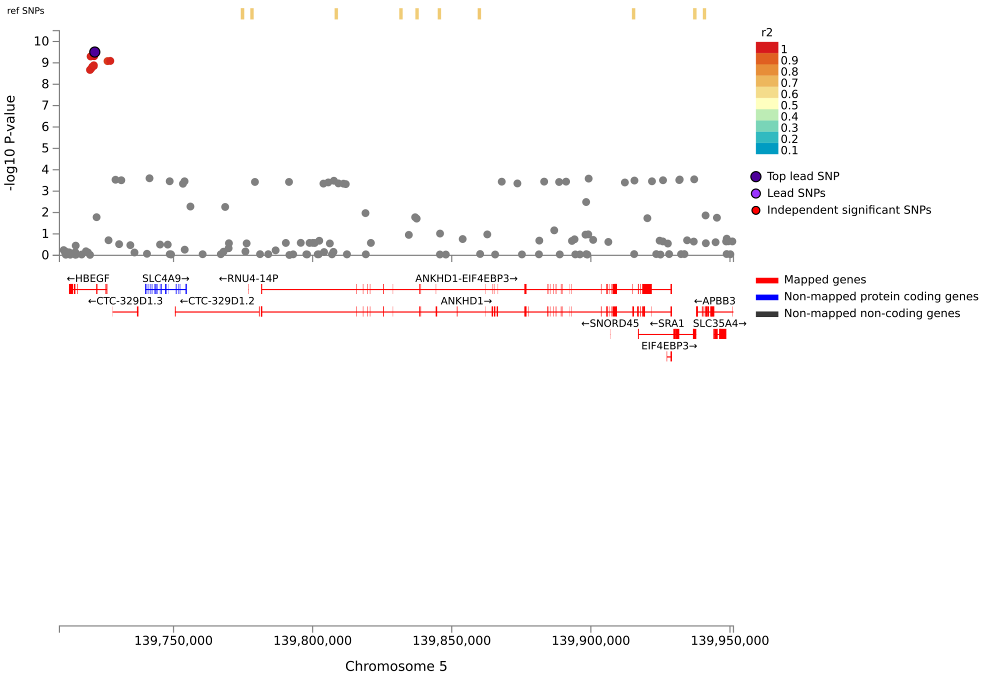


Locus 4, SNORA73, Anterior Body Mean Thickness, rs76928645

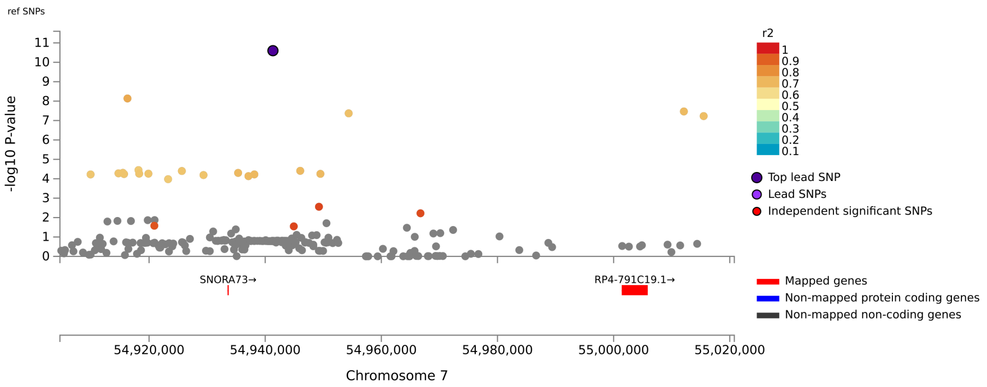

Locus 5, PLEC, Anterior Body Mean Thickness, rs7464572

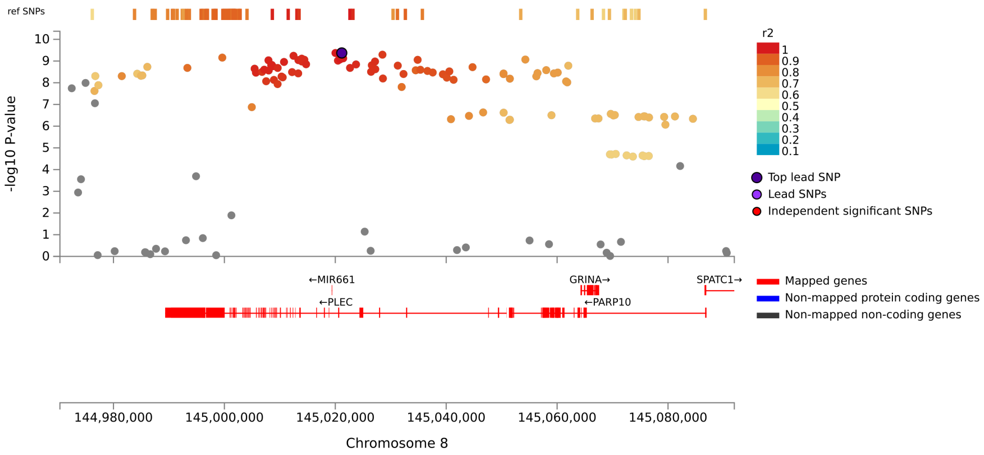


Locus 6, CABYR, Anterior Body Mean Thickness, rs752797

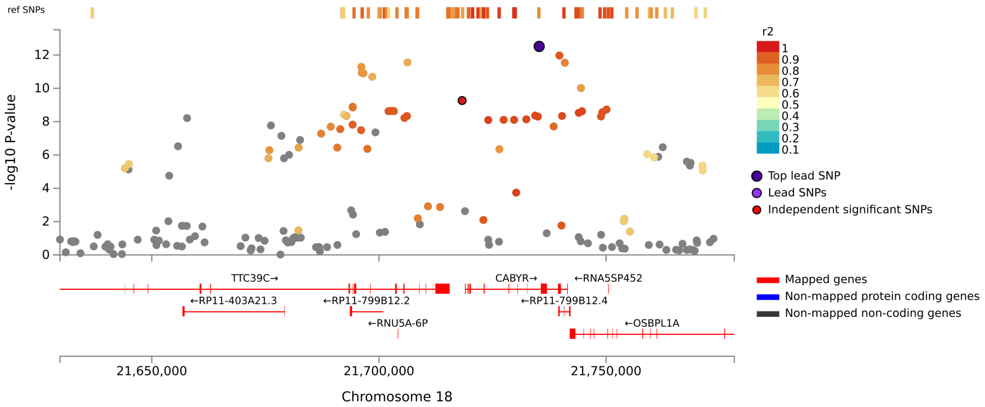

Locus 1, STRN, Posterior Body Area, rs1861435

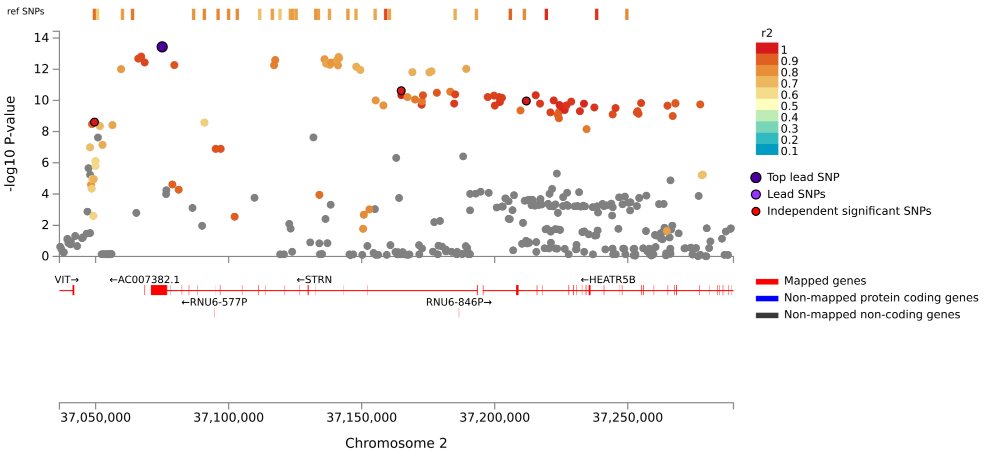


Locus 2, RP11-88I21.2, Posterior Body Area, rs35962269

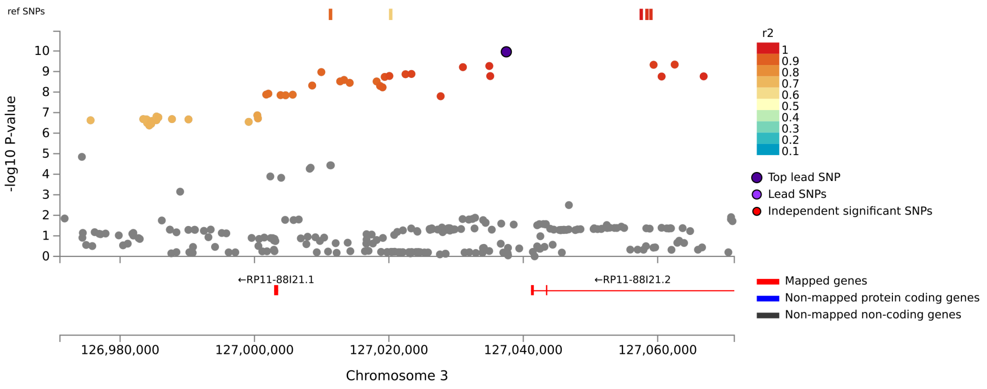

Locus 3, IQCJ-SCHIP1:IQCJ, Posterior Body Area, rs6792979

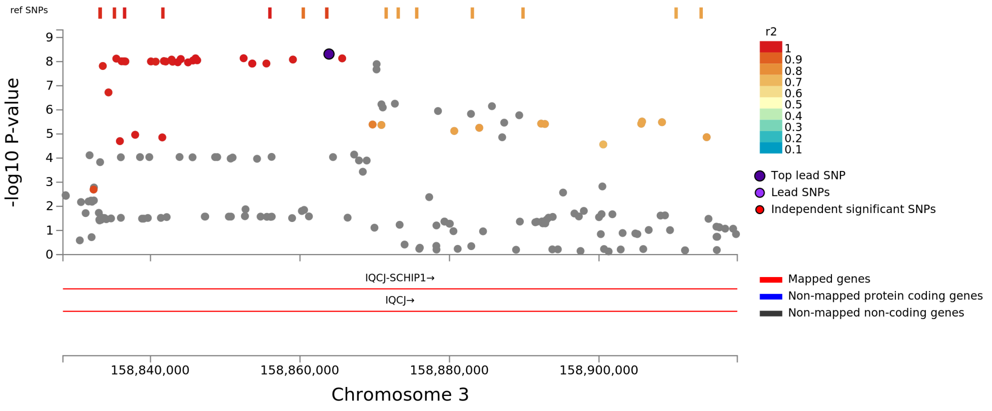


Locus 4, FIP1L1, Posterior Body Area, rs6835429

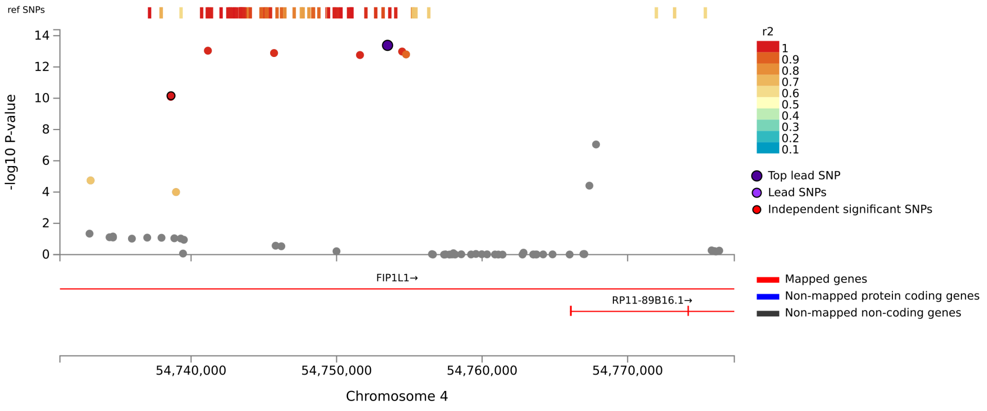

Locus 5, HBEGF, Posterior Body Area, rs4150214

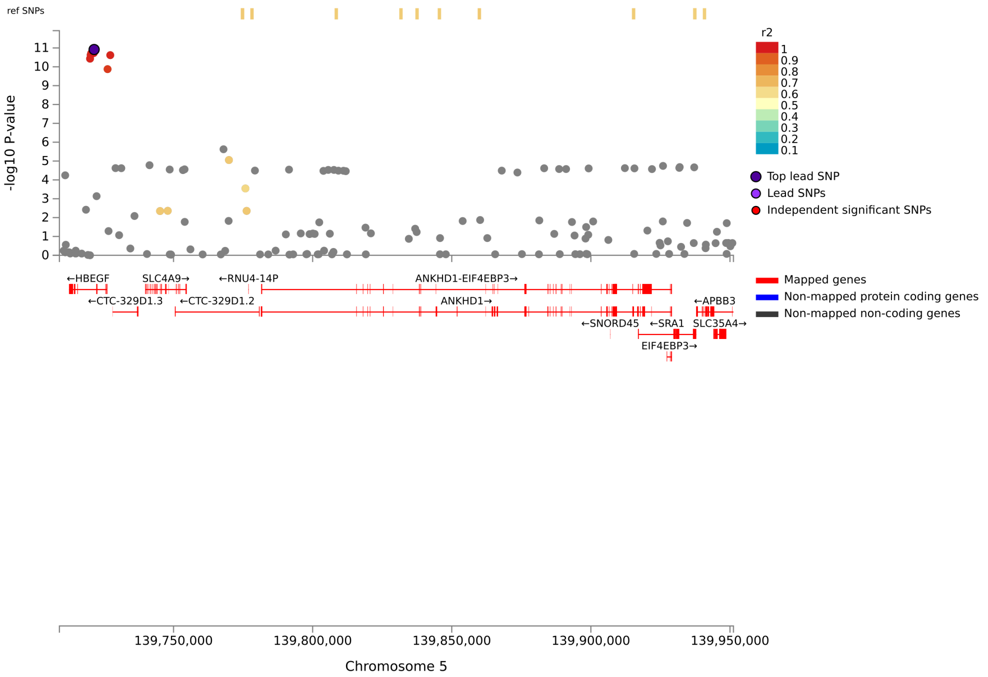


Locus 6, FOXO3, Posterior Body Area, rs1268163

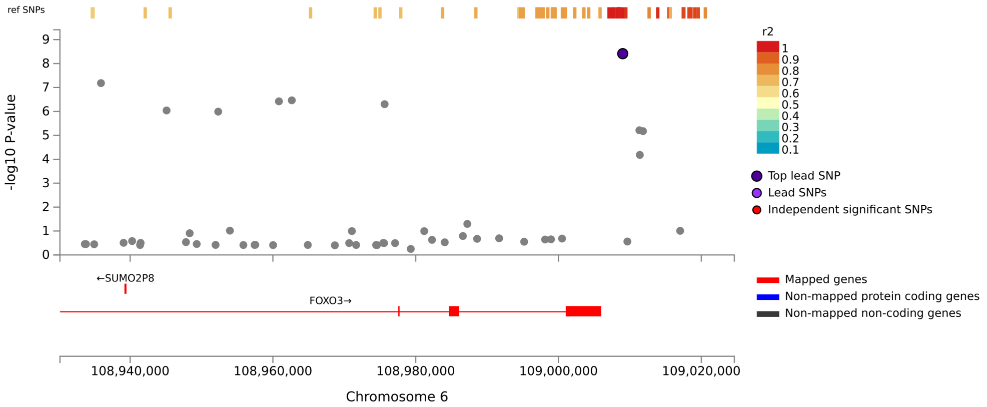

Locus 7, RP1-34H18.1, Posterior Body Area, rs4761385

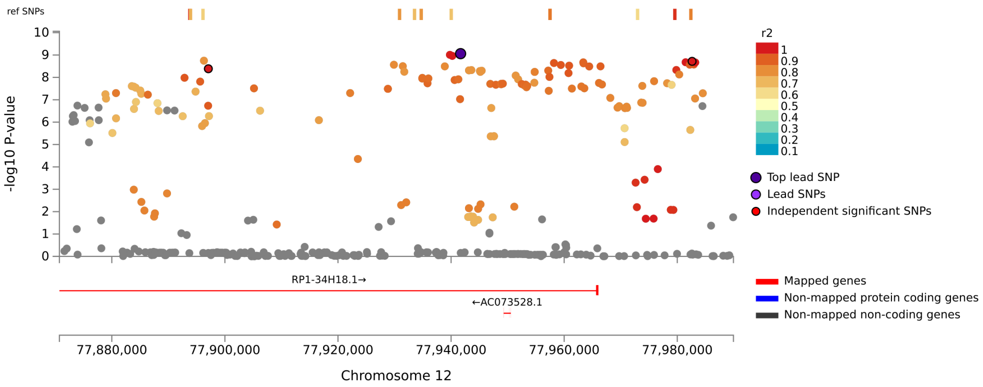


Locus 8, TTC39C, Posterior Body Area, rs12967609

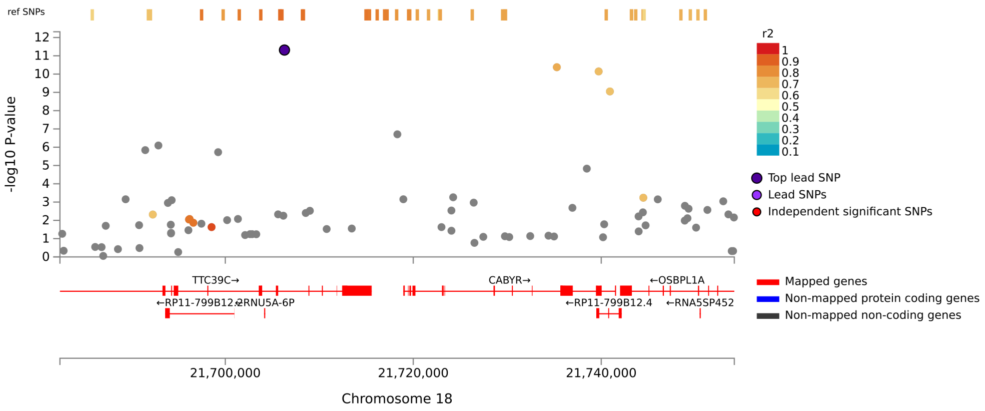

Locus 1, FIP1L1, Posterior Body Mean Thickness, rs6835429

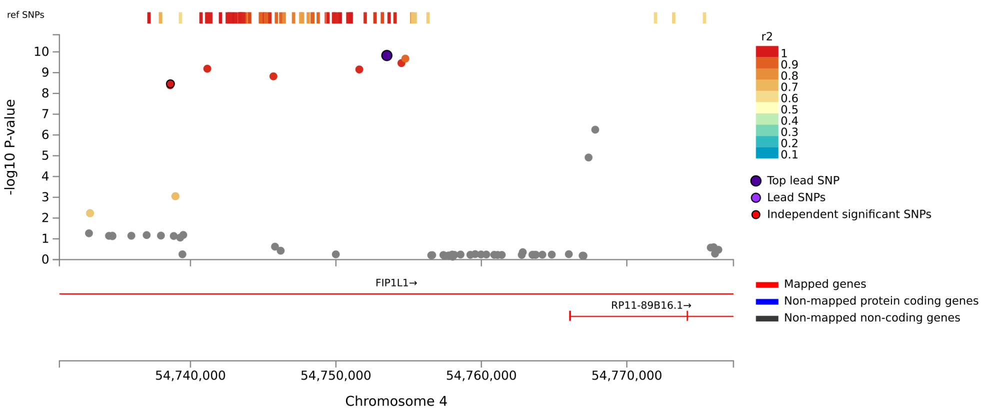


Locus 2, PARP10, Posterior Body Mean Thickness, rs11136344

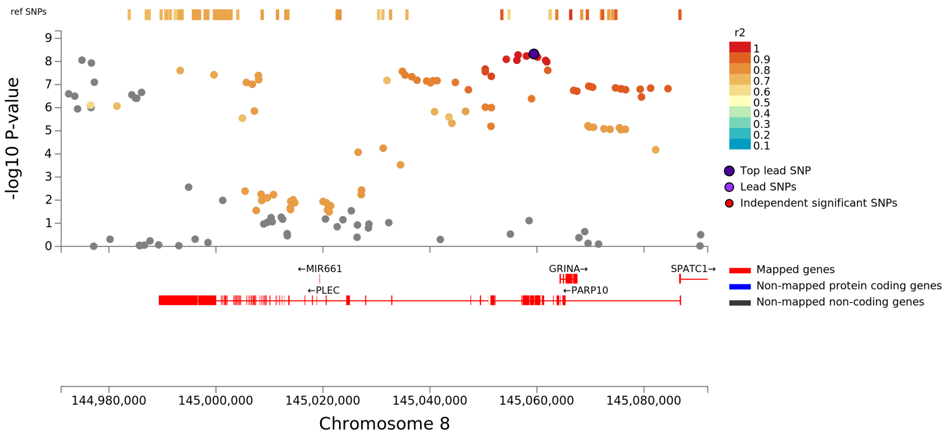

### Locus 3, C16orf95, Posterior Body Mean Thickness, rs11863620

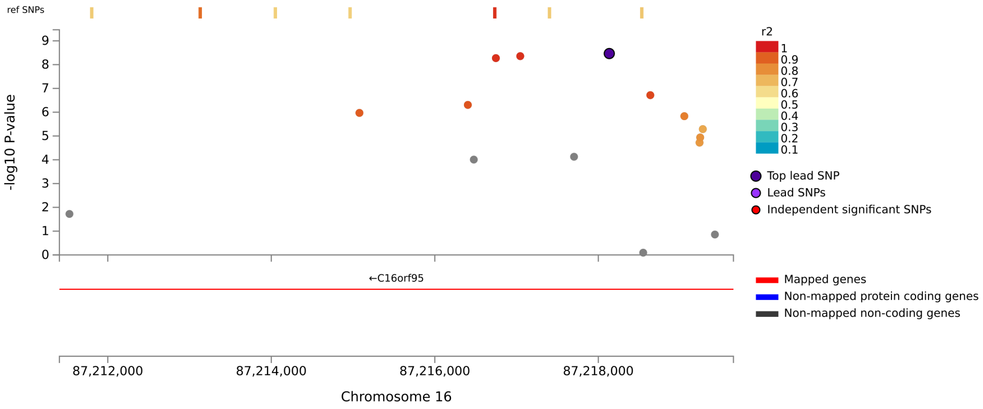


### Locus 4, TTC39C, Posterior Body Mean Thickness, rs12967609

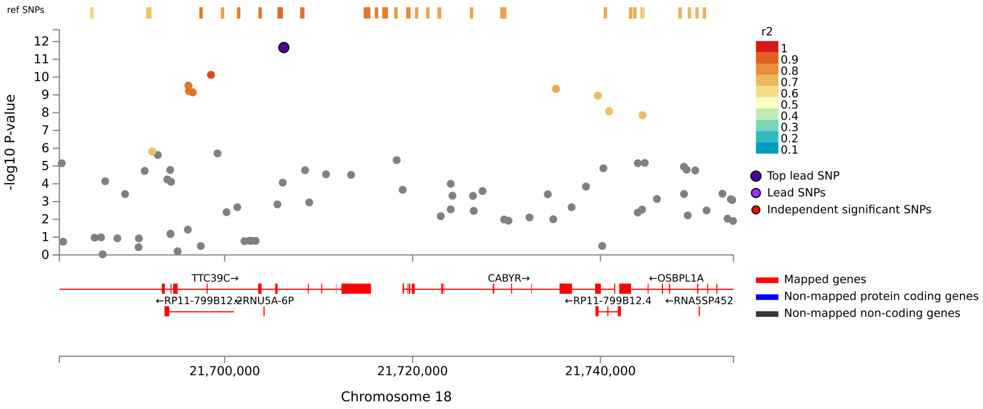

Locus 1, RP1-37C10.3, Isthmus Area, rs6682671

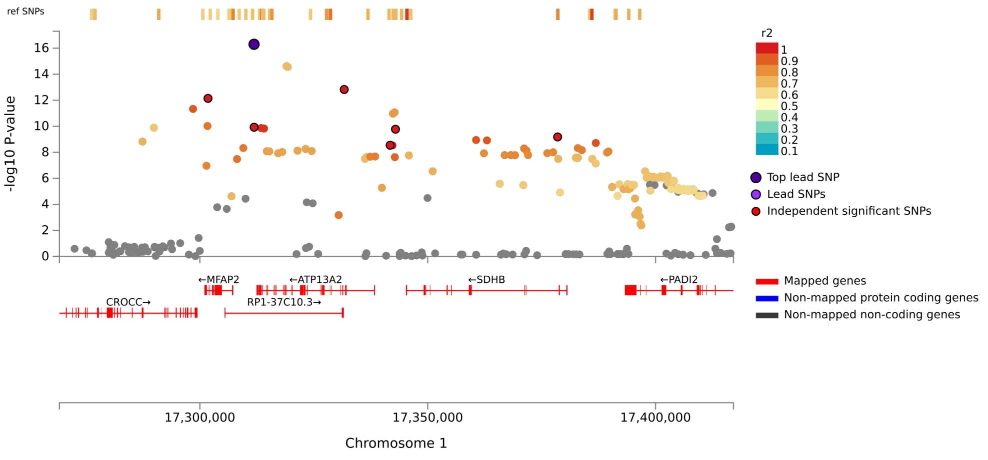


Locus 2, STRN, Isthmus Area, rs10193295

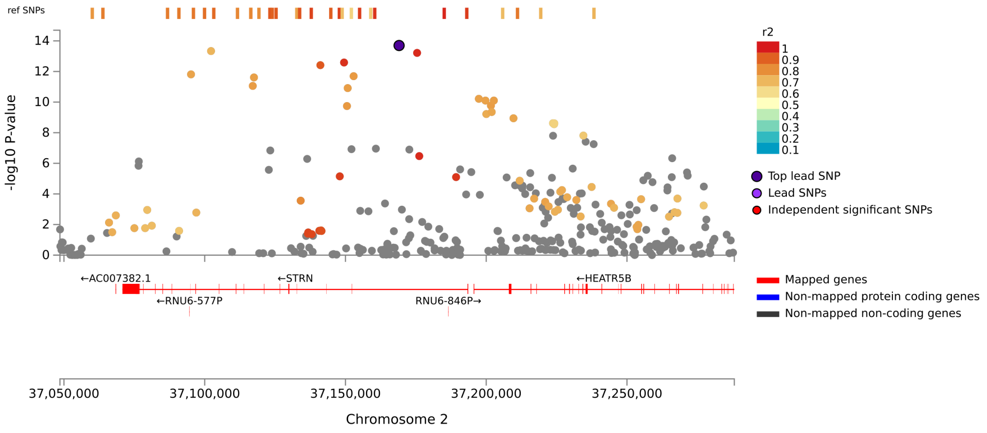

Locus 3, CCDC75P1, Isthmus Area, rs9860128

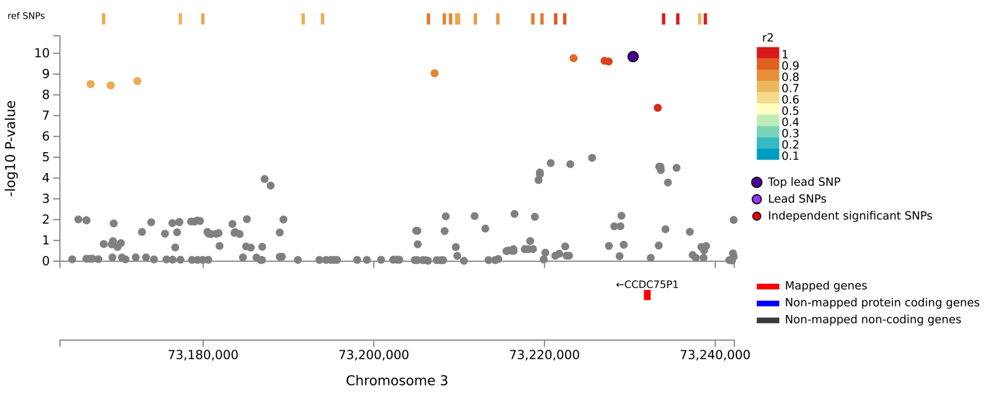


Locus 4, IQCJ-SCHIP1:IQCJ, Isthmus Area, rs11717303

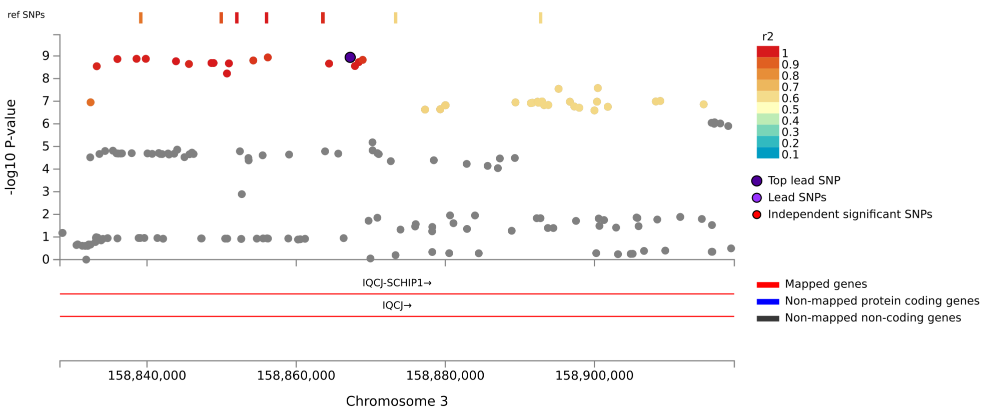

Locus 5, TNIK, Isthmus Area, rs2035913

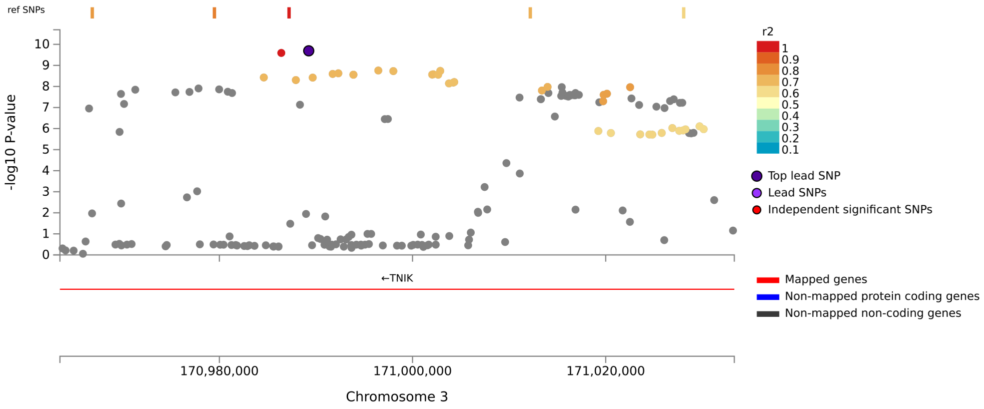


Locus 6, ADD1, Isthmus Area, rs12645803

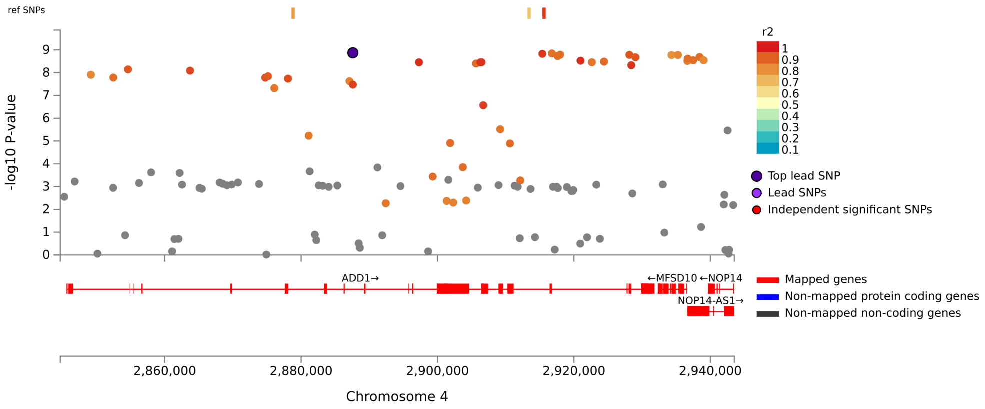

Locus 7, TBC1D14, Isthmus Area, rs529090333

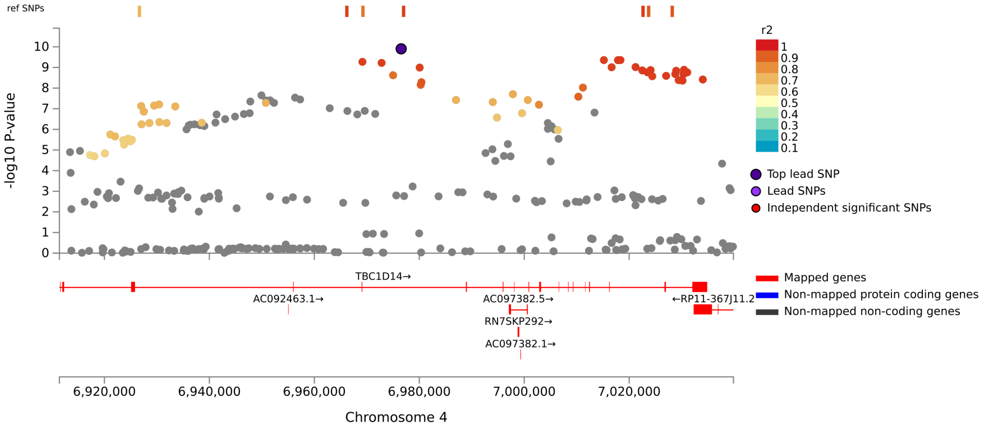


Locus 8, FIP1L1:RP11-89B16.1, Isthmus Area, rs62297576

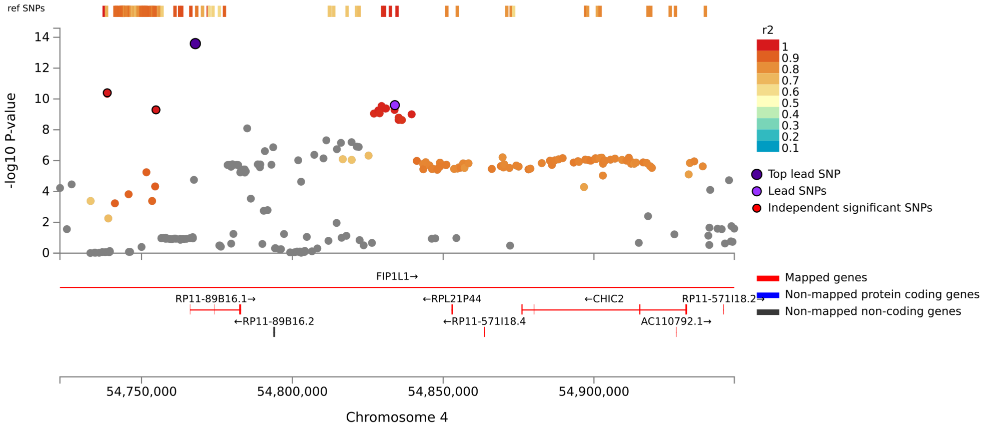

Locus 9, CTB-118N6.2, Isthmus Area, rs3806915

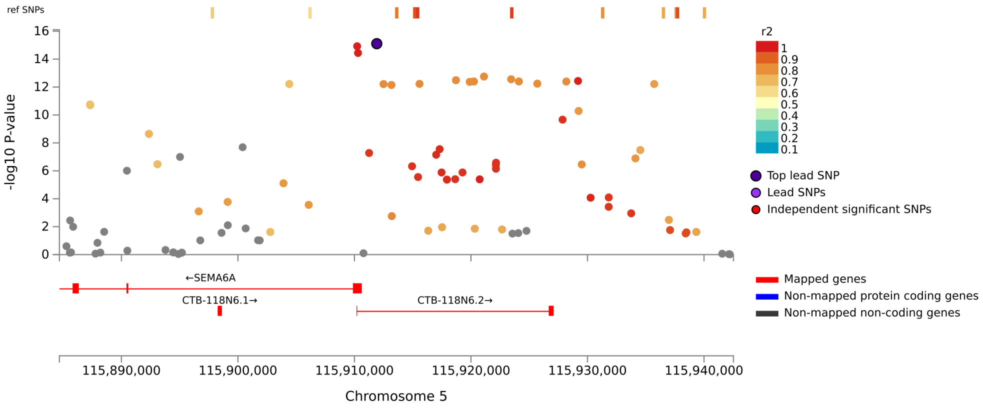


Locus 10, HBEGF, Isthmus Area, rs4150210

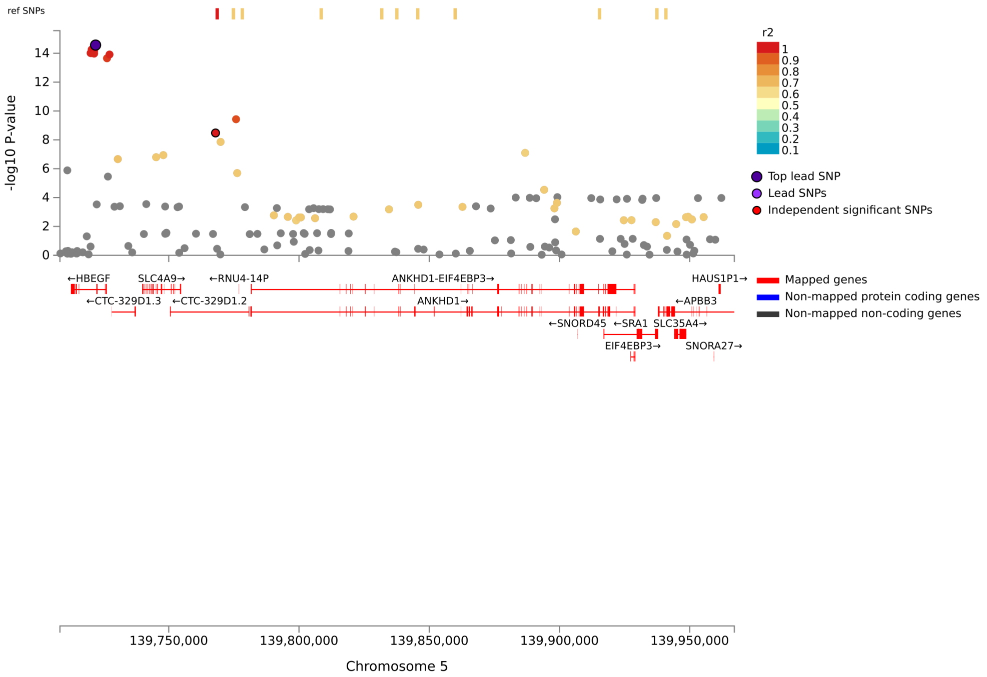

#### Locus 11, FOXO3, Isthmus Area, rs1268163

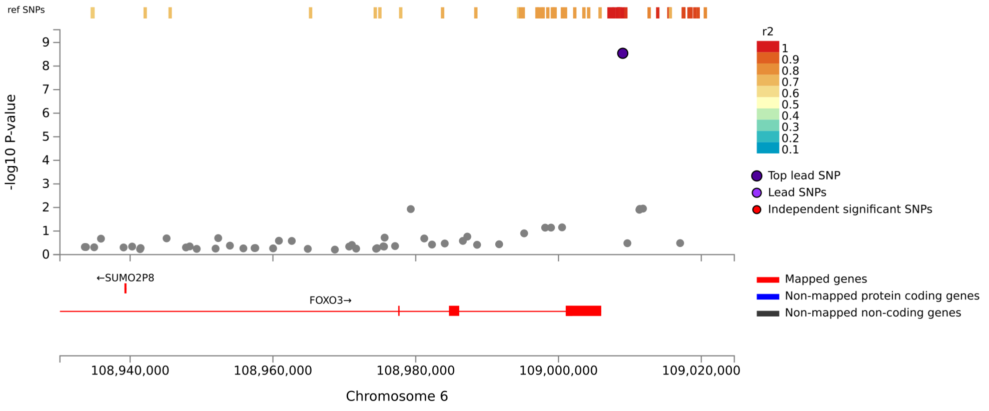


#### Locus 12, PLEC, Isthmus Area, rs6991364

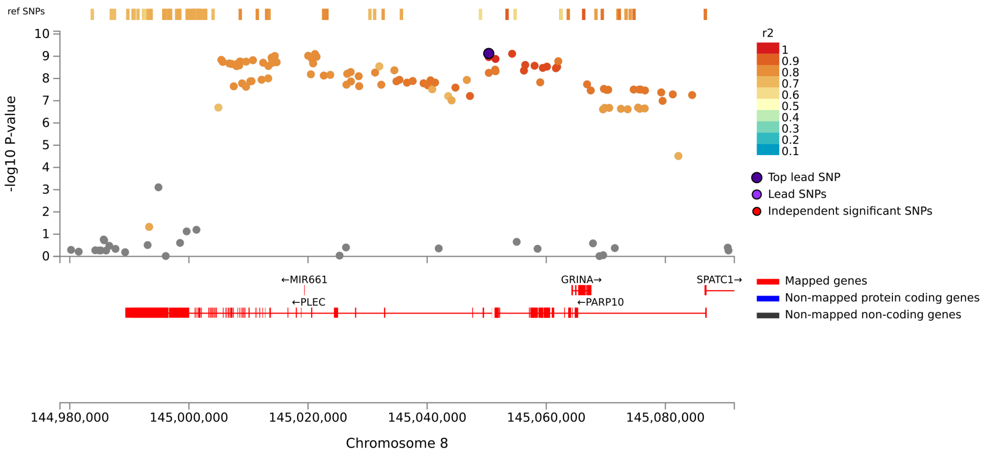

#### Locus 13, FAM107B:RP11-7C6.1, Isthmus Area, rs61845075

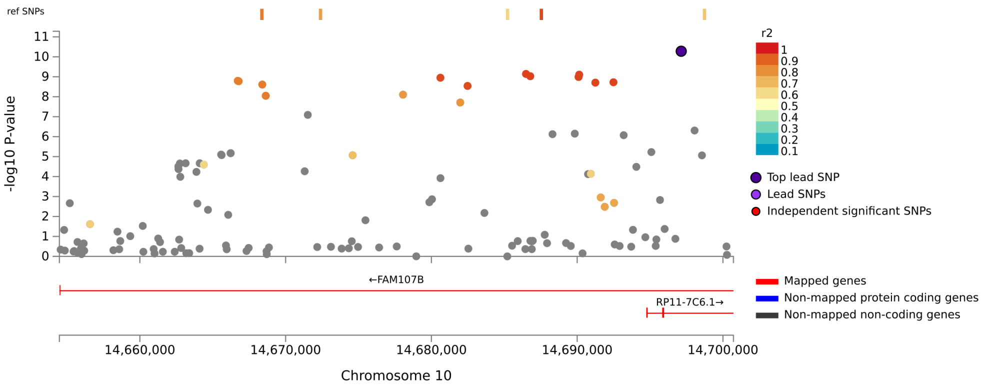


#### Locus 14, KIAA1598, Isthmus Area, rs11197861

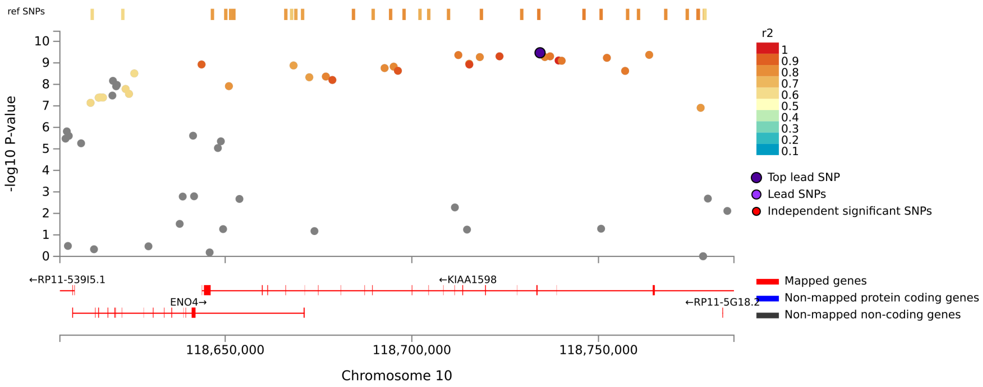

Locus 15, NAV2, Isthmus Area, rs2585757

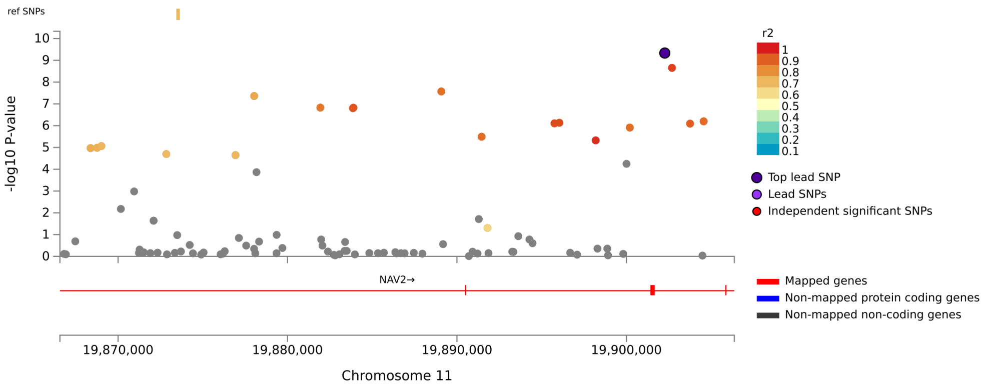


Locus 16, C16orf95, Isthmus Area, rs4843552

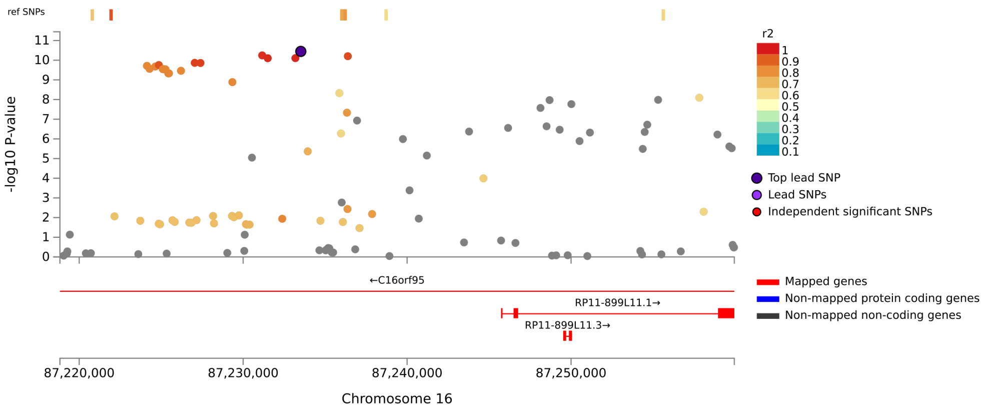

### Locus 17, CABYR, Isthmus Area, rs752797

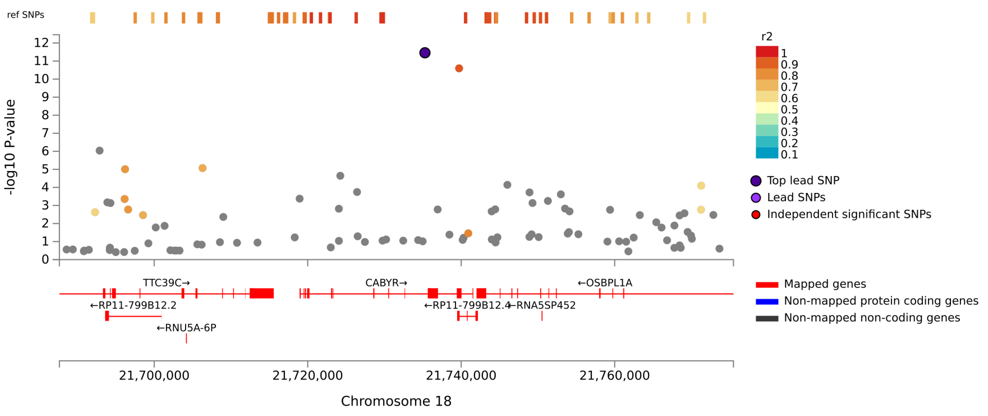

Locus 1, RP1-37C10.3, Isthmus Mean Thickness, rs6682671

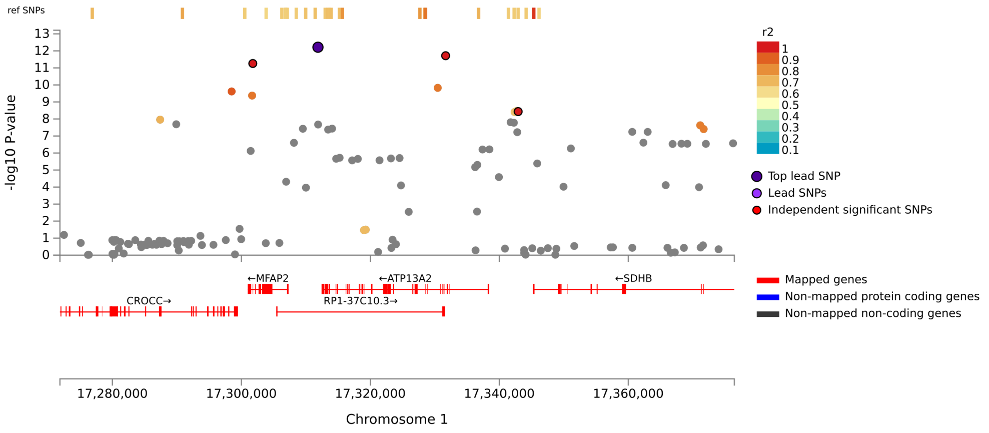


Locus 2, STRN, Isthmus Mean Thickness, rs7561572

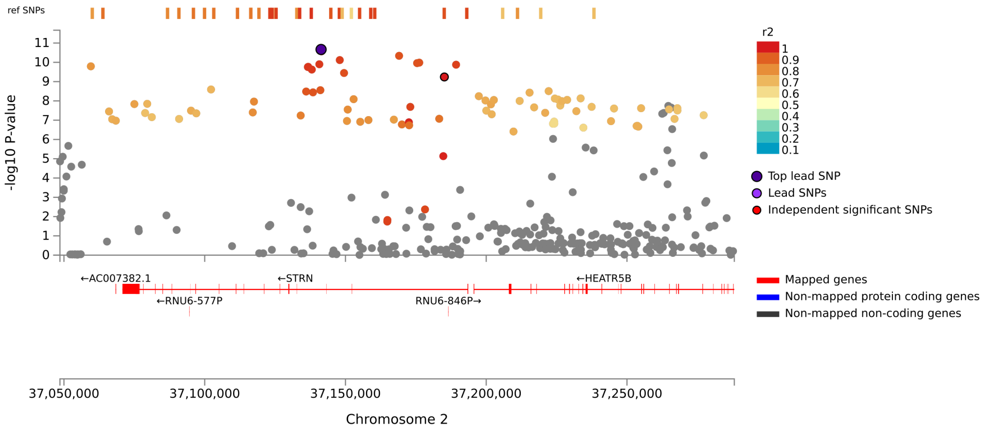

Locus 3, WDPCP, Isthmus Mean Thickness, rs35474183

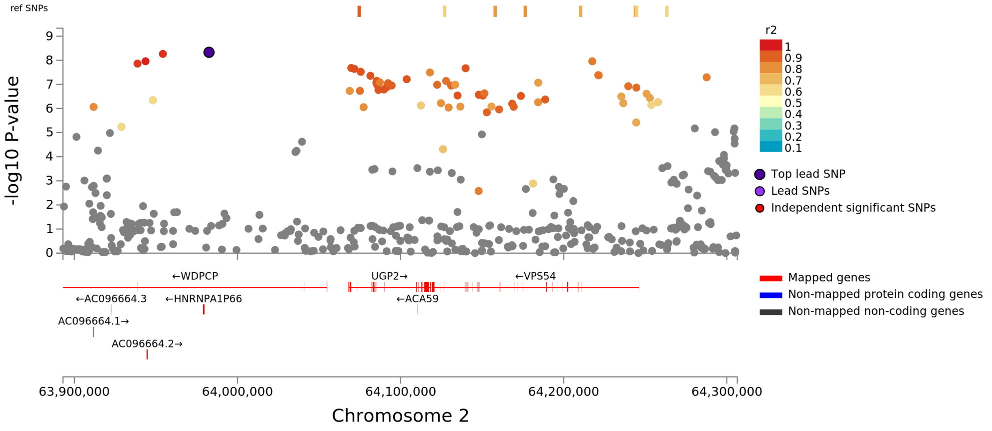


Locus 4, CCDC75P1, Isthmus Mean Thickness, rs9857083

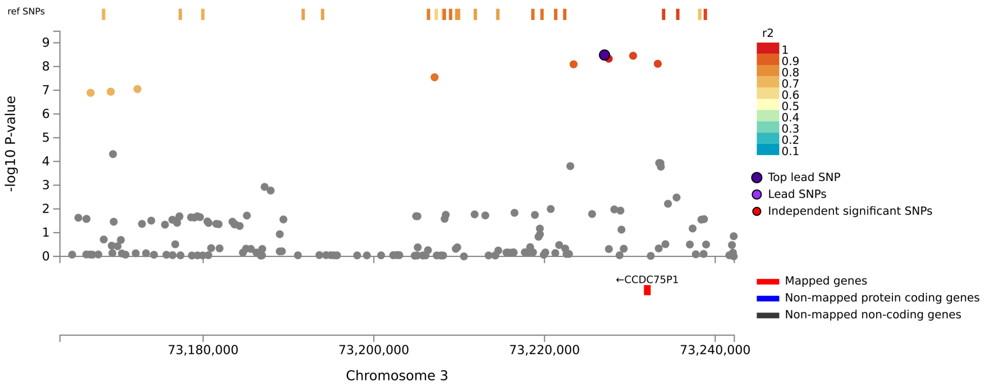

Locus 5, FIP1L1, Isthmus Mean Thickness, rs6554139

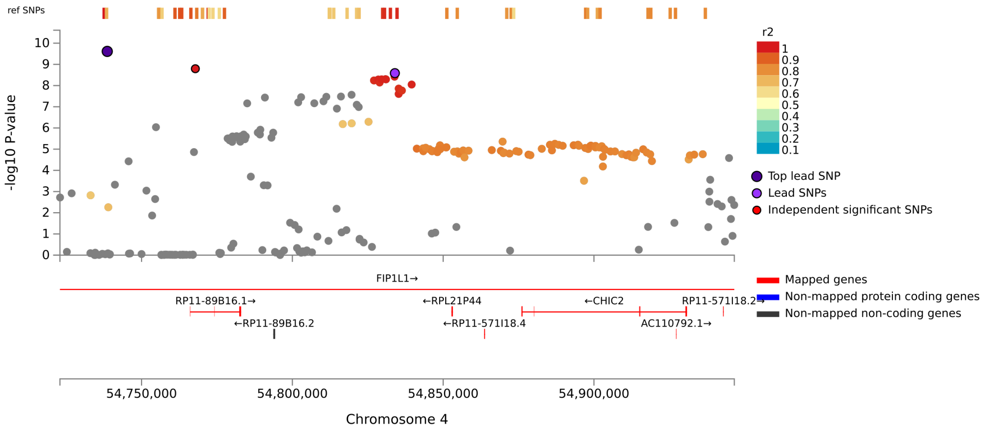


Locus 6, CTB-118N6.2, Isthmus Mean Thickness, rs1345707

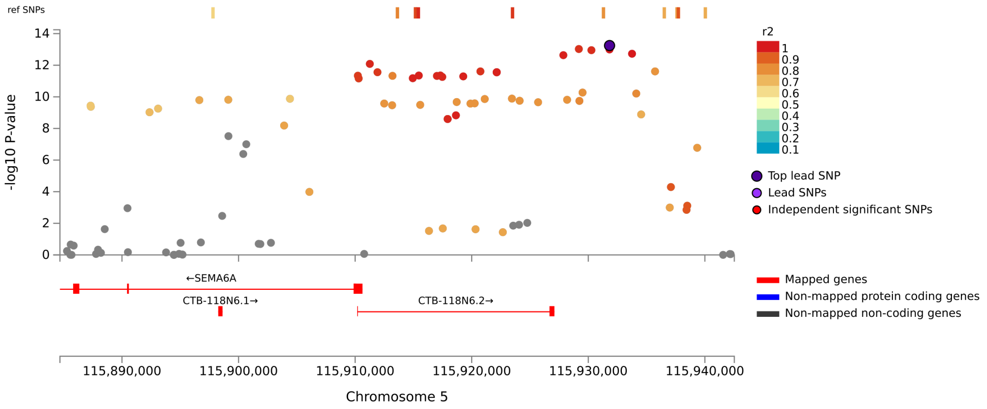

#### Locus 7, HBEGF, Isthmus Mean Thickness, rs58992612

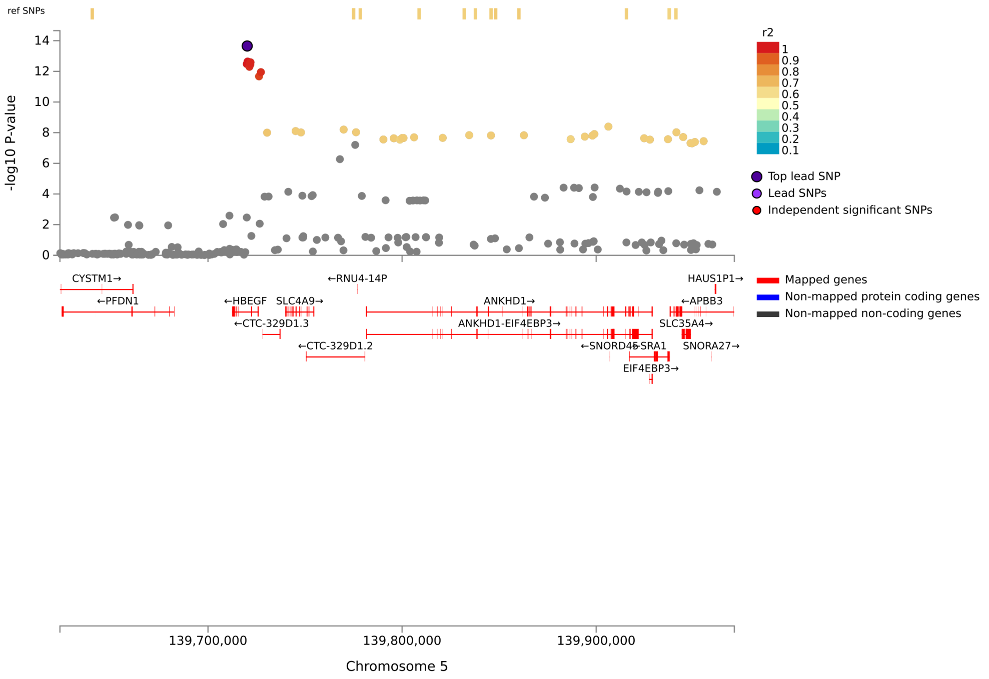


#### Locus 8, SLC45A4, Isthmus Mean Thickness, rs4961258

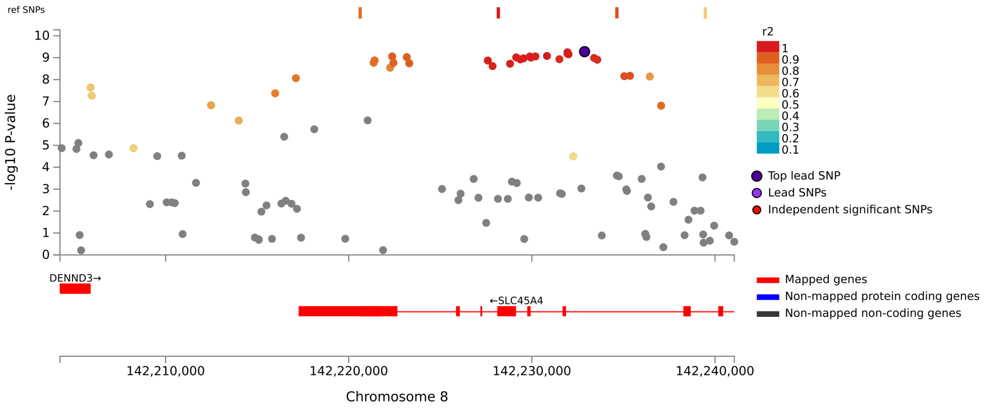

### Locus 9, C16orf95, Isthmus Mean Thickness, rs4843550

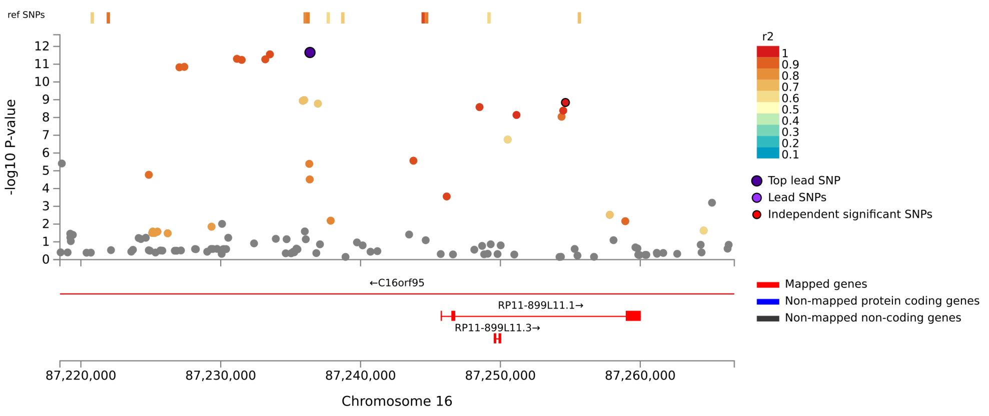

Locus 1, RP1-37C10.3:ATP13A2, Splenium Area, rs2076603

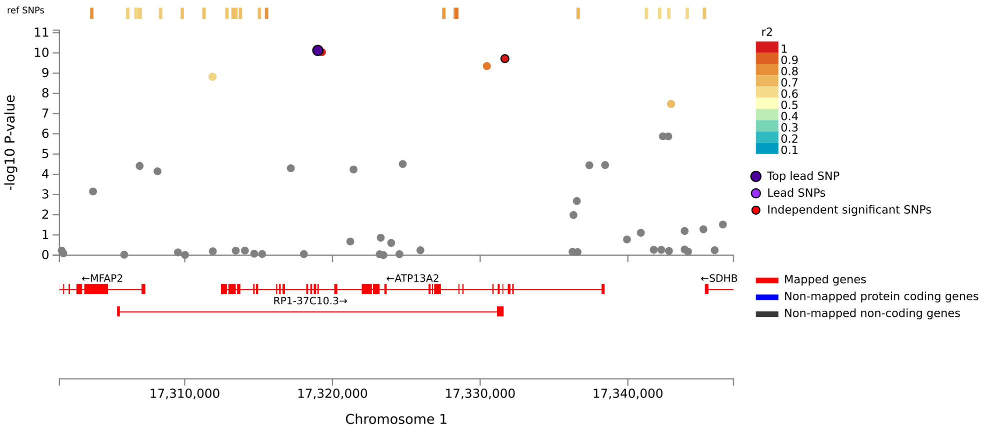


Locus 2, AKT3, Splenium Area, rs67027895

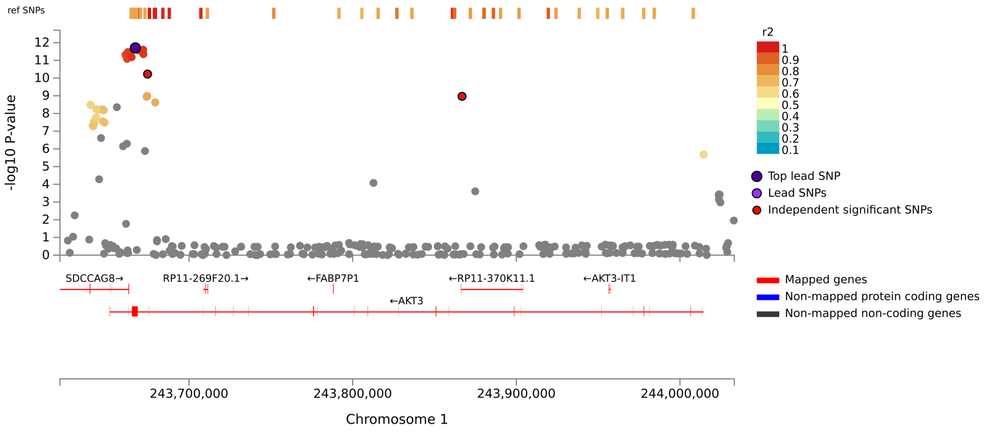

Locus 3, AC007382.1, Splenium Area, rs711244

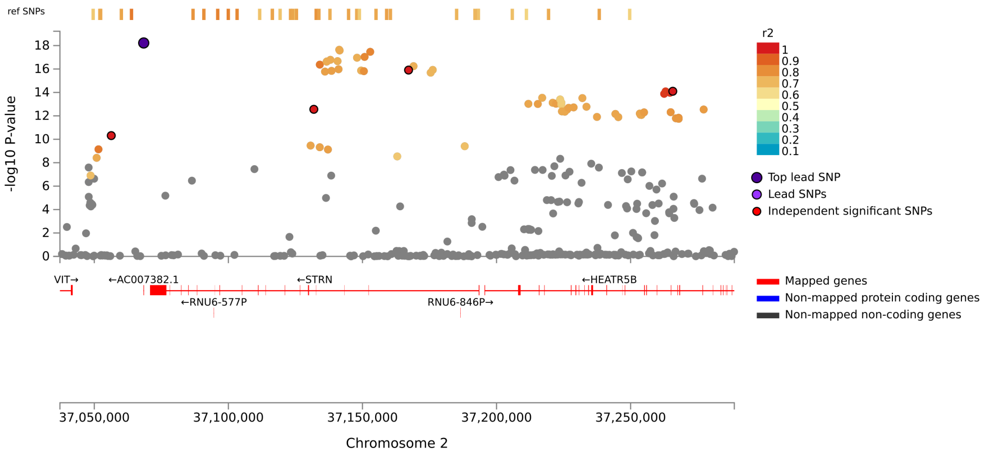


Locus 4, XPO1, Splenium Area, rs7570830

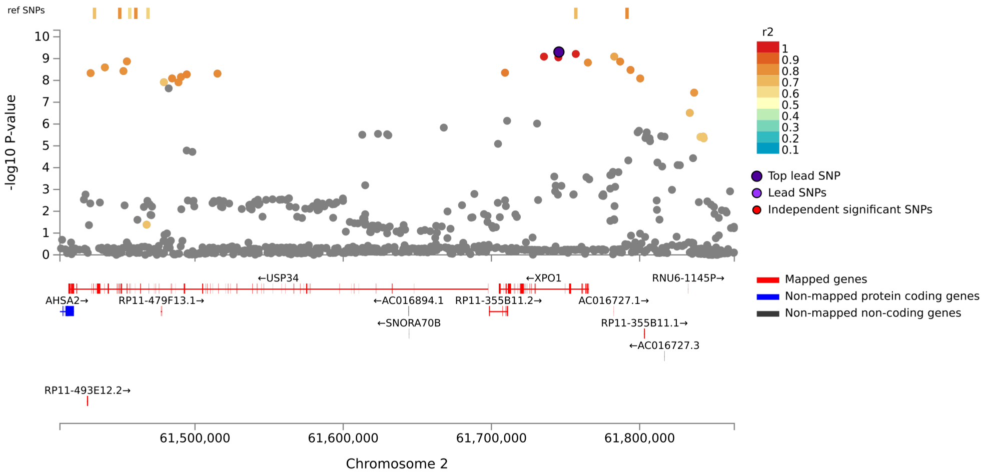

Locus 5, RP11-493K19.3, Splenium Area, rs2071206

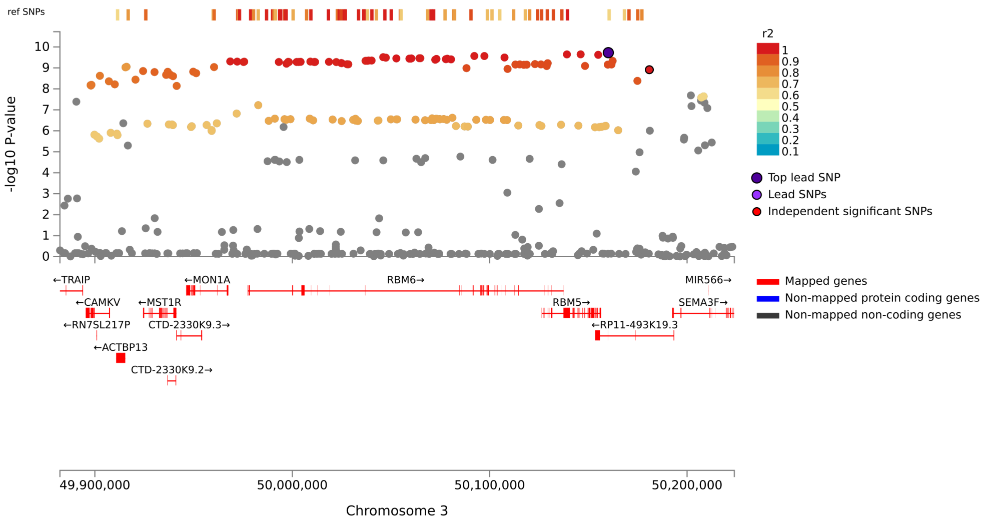


Locus 6, CCDC75P1, Splenium Area, rs56934393

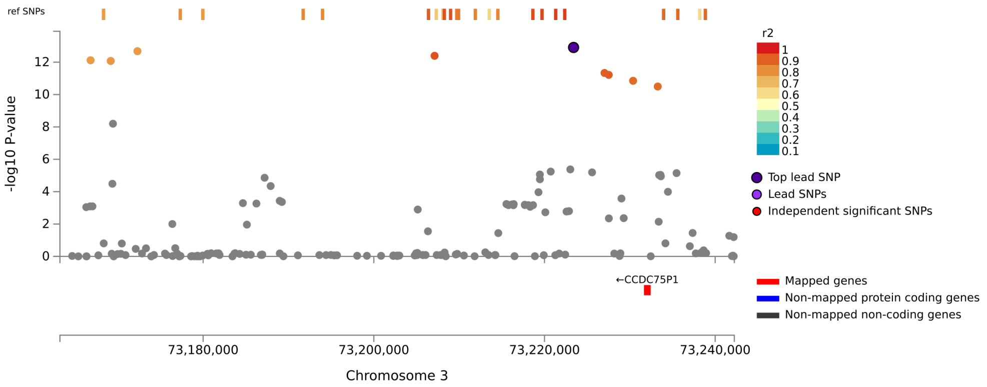

Locus 7, IQCJ-SCHIP1:IQCJ, Splenium Area, rs11717303

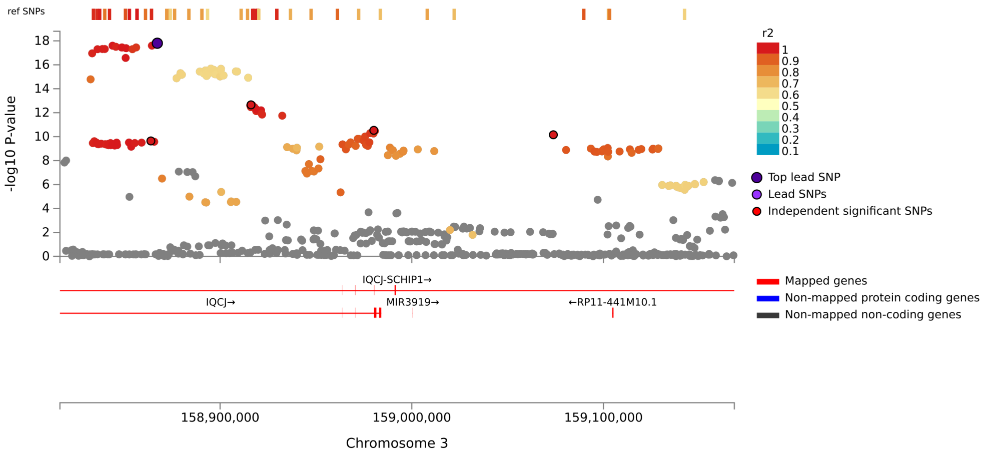


Locus 8, IQCJ-SCHIP1, Splenium Area, rs13064756

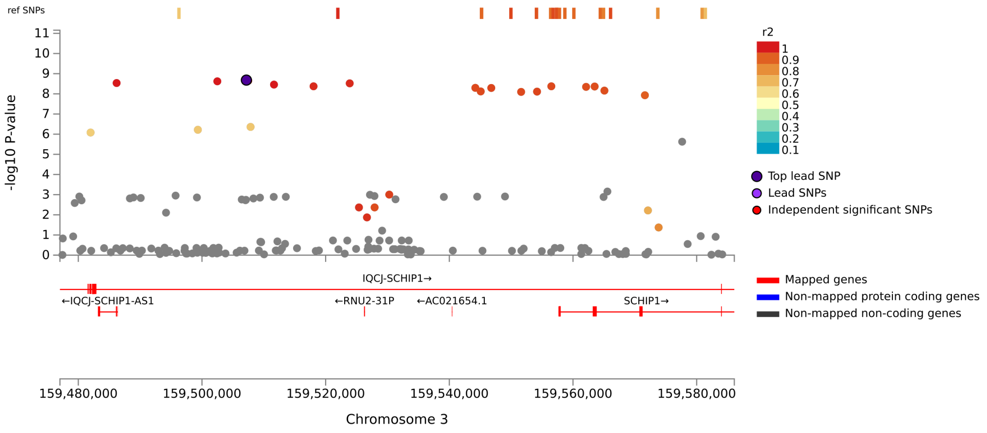

#### Locus 9, TNIK, Splenium Area, rs6444974

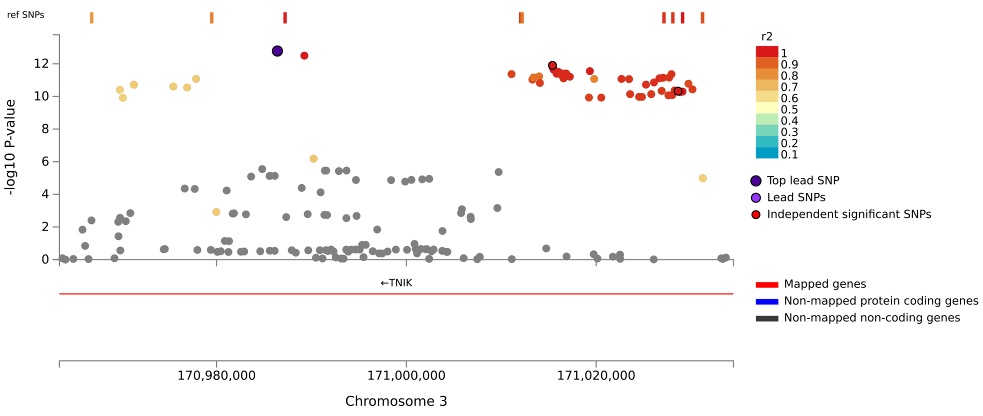


#### Locus 10, TBC1D14, Splenium Area, rs199601576

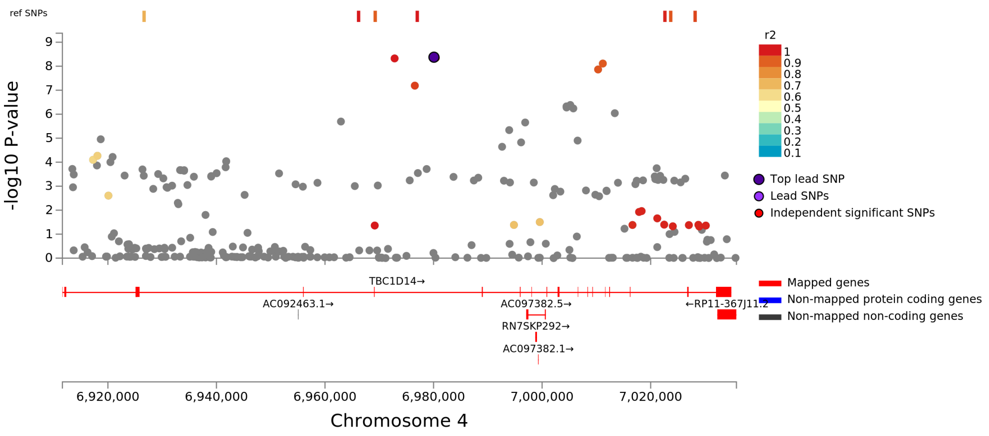

Locus 11, FIP1L1, Splenium Area, rs6849897

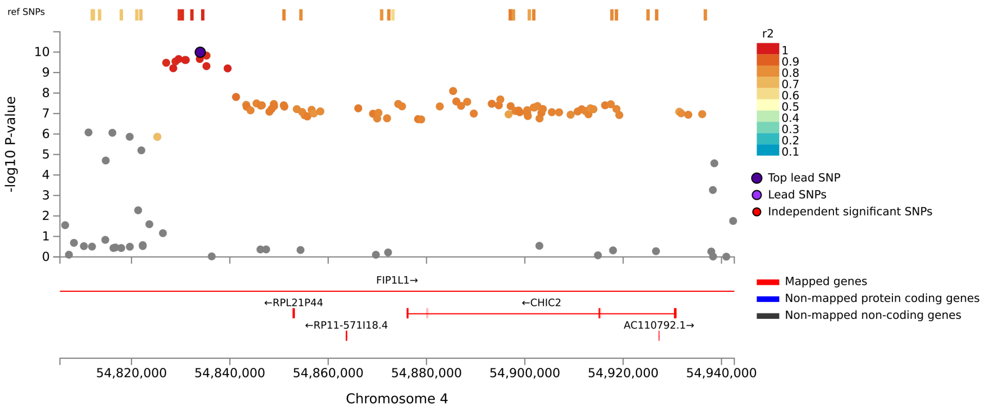


Locus 12, STPG2:RP11-681L8.1:STPG2-AS1, Splenium Area, rs6231

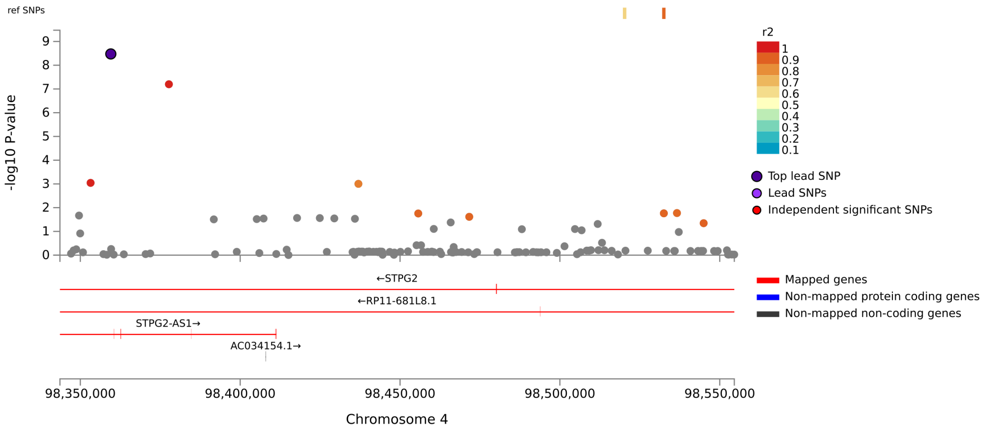

Locus 13, FBXL7, Splenium Area, rs4702099

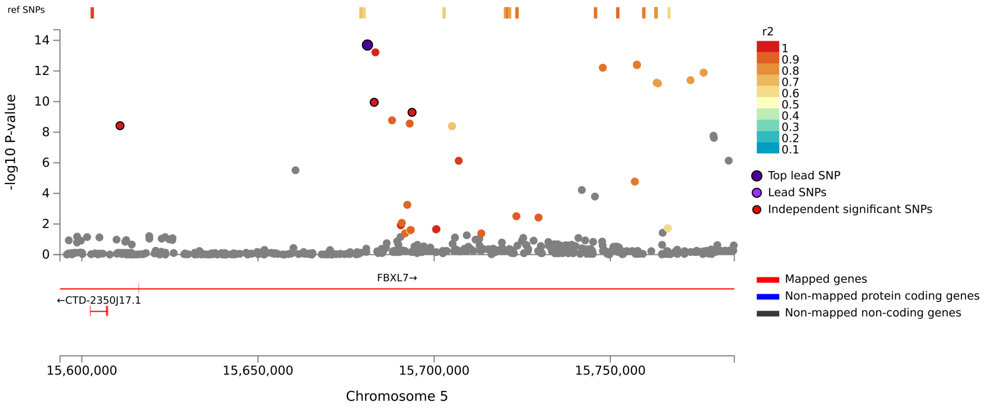


Locus 14, CTB-118N6.2, Splenium Area, rs686664

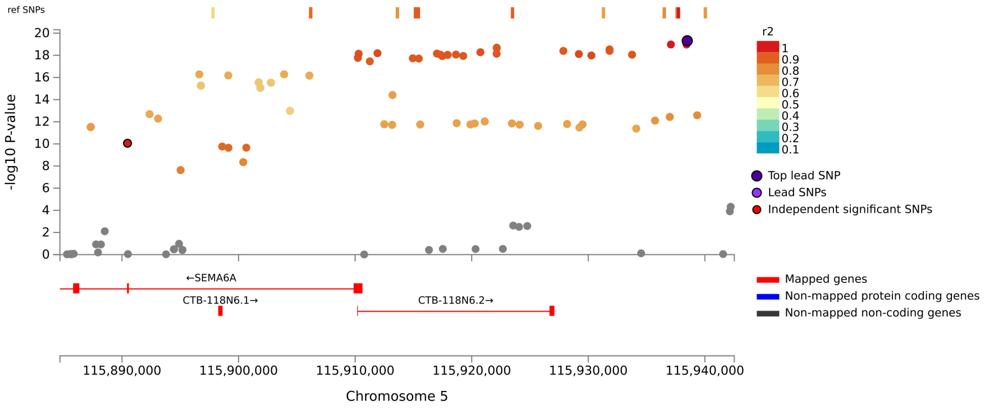

#### Locus 15, HBEGF, Splenium Area, rs58992612

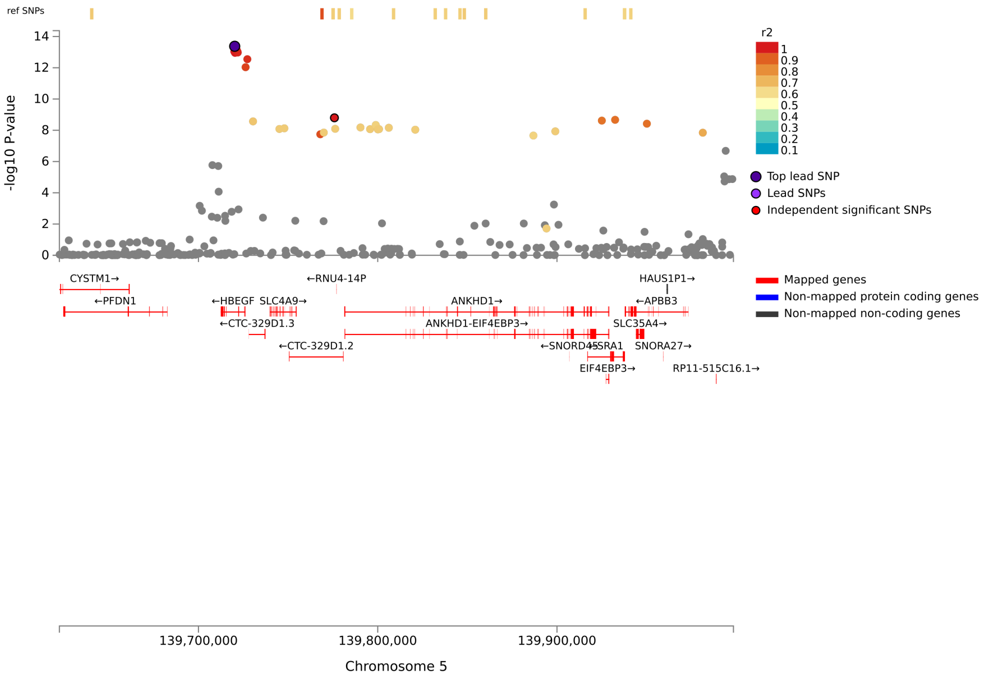


#### Locus 16, HGF, Splenium Area, rs60392694

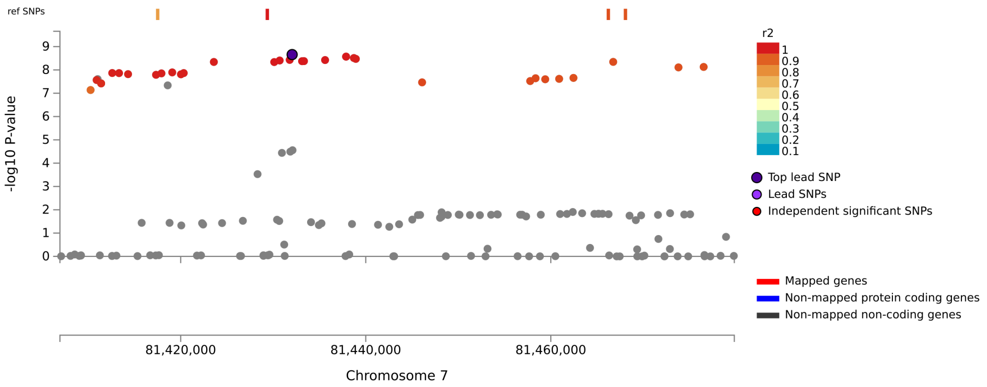

Locus 17, SEMA3A, Splenium Area, rs73712705

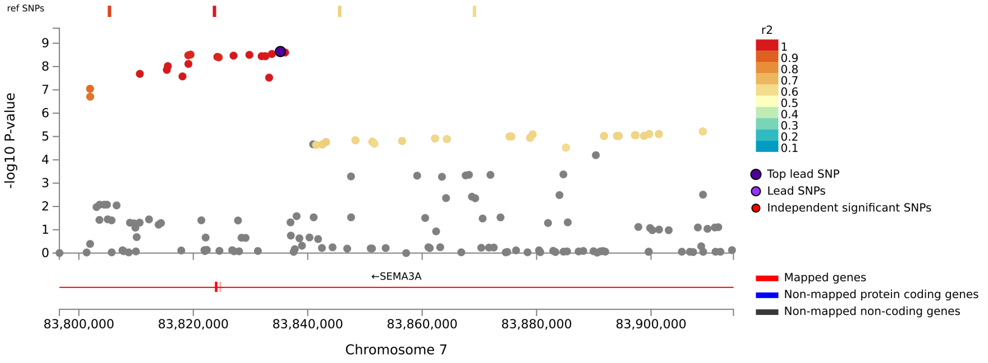


Locus 18, BICD2, Splenium Area, rs10992447

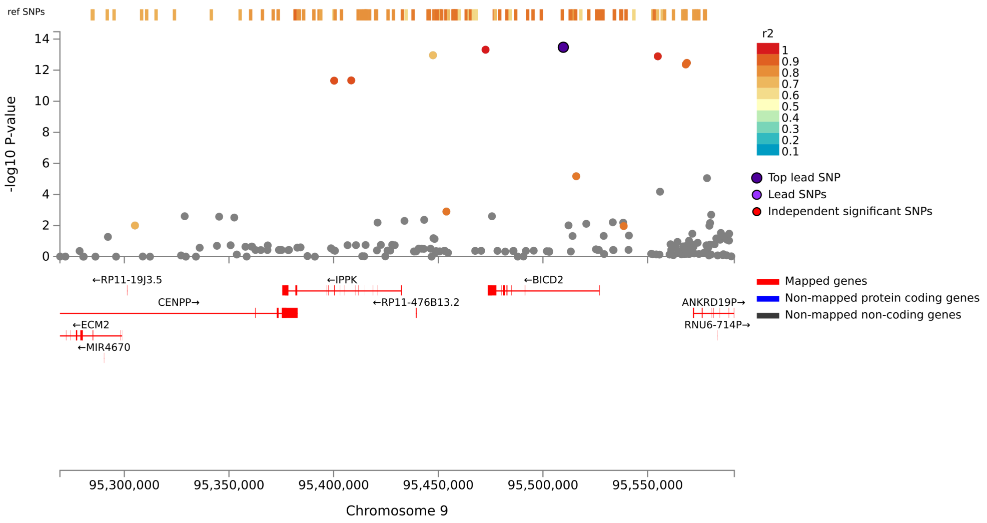

#### Locus 19, FAM107B, Splenium Area, rs10906729

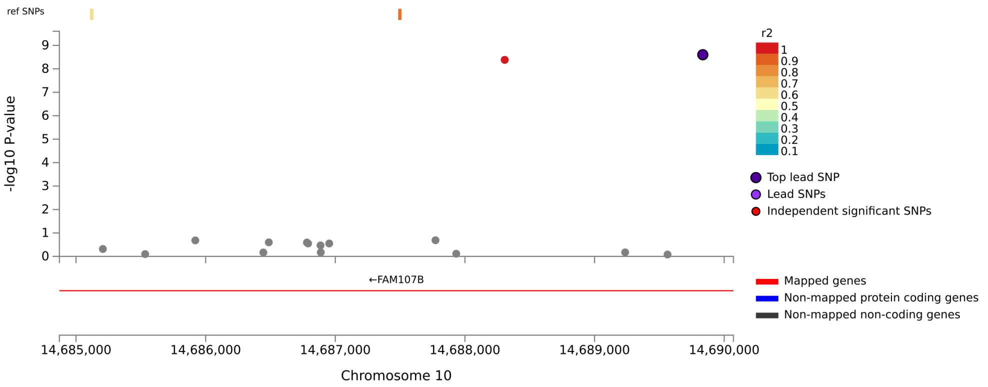


#### Locus 20, ENO4:KIAA1598, Splenium Area, rs10886016

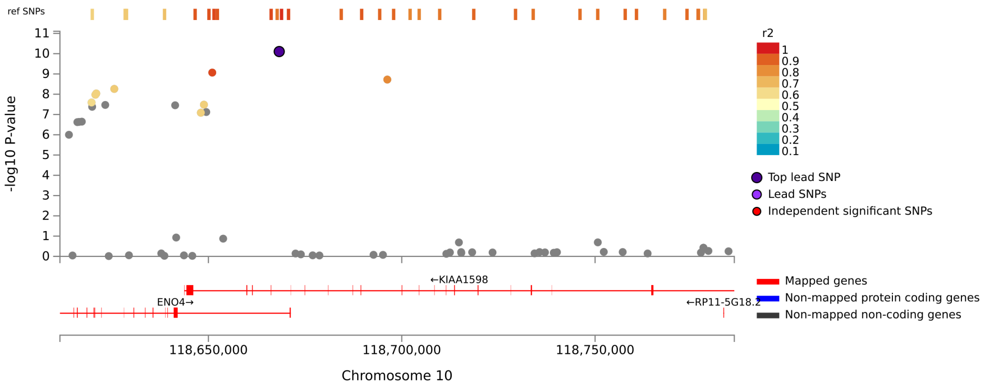

#### Locus 21, RP11-12J10.3:FAM53B, Splenium Area, rs10901814

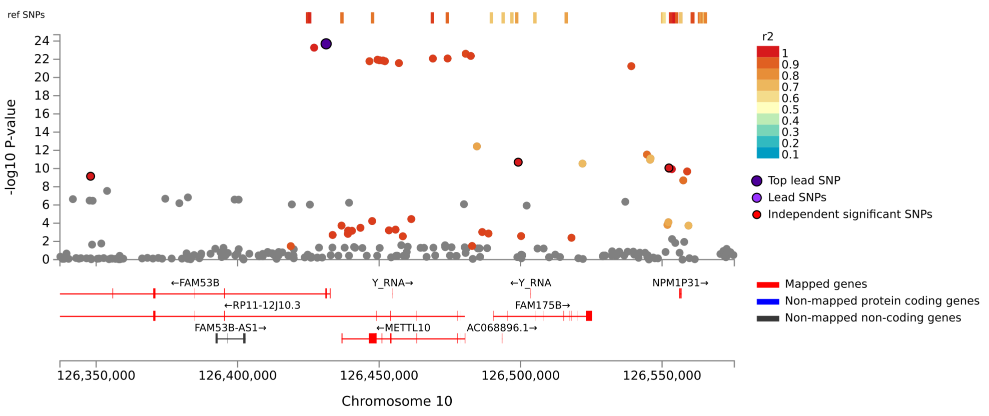


#### Locus 22, BRSK2, Splenium Area, rs7947308

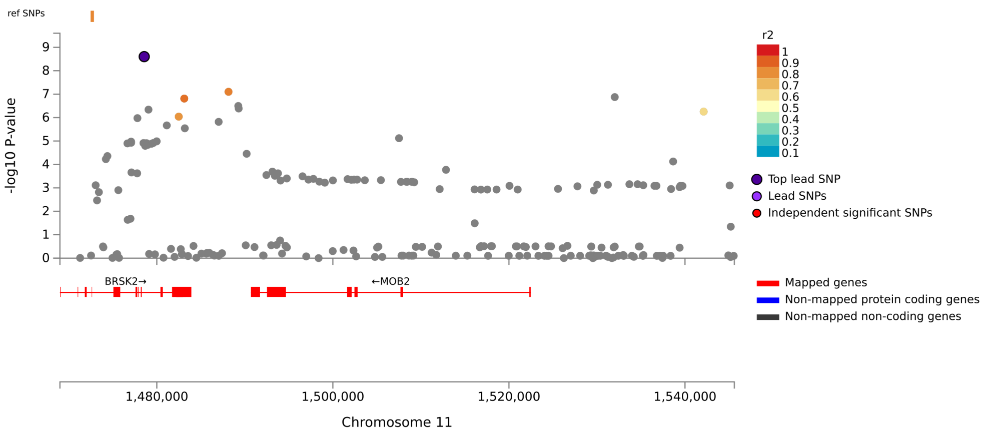

#### Locus 23, RRAS2, Splenium Area, rs35247669

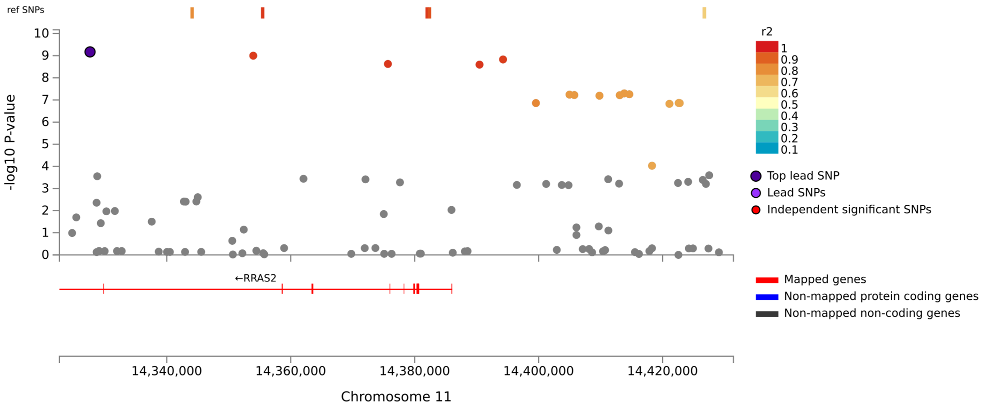


#### Locus 24, TAOK1, Splenium Area, rs8081085

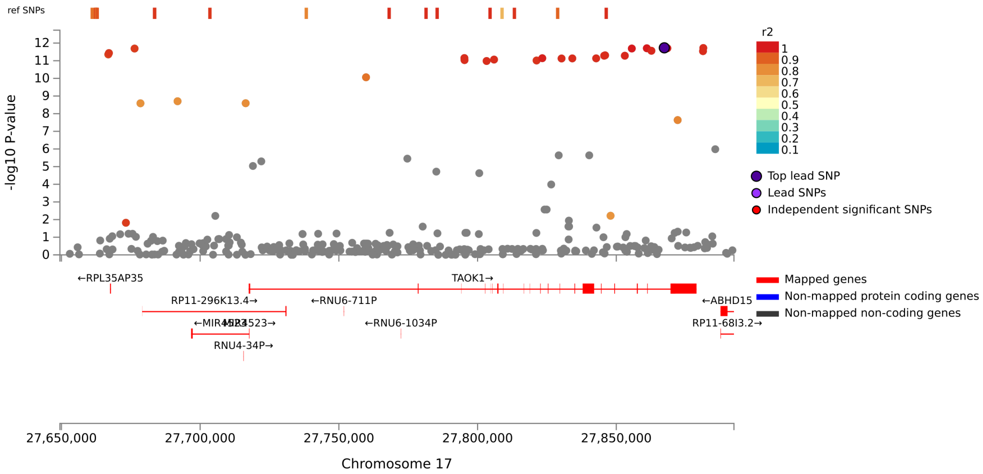

Locus 1, STRN, Splenium Mean Thickness, rs62132522

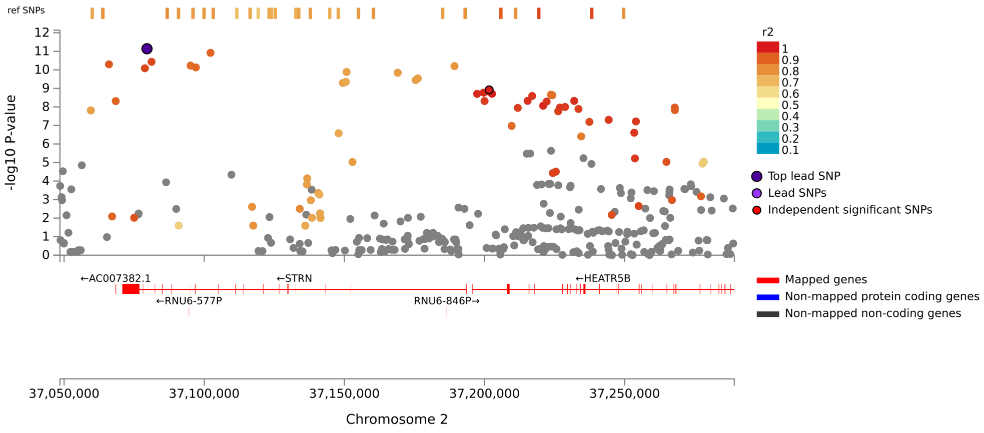


Locus 2, IQCJ-SCHIP1:IQCJ, Splenium Mean Thickness, rs55938743

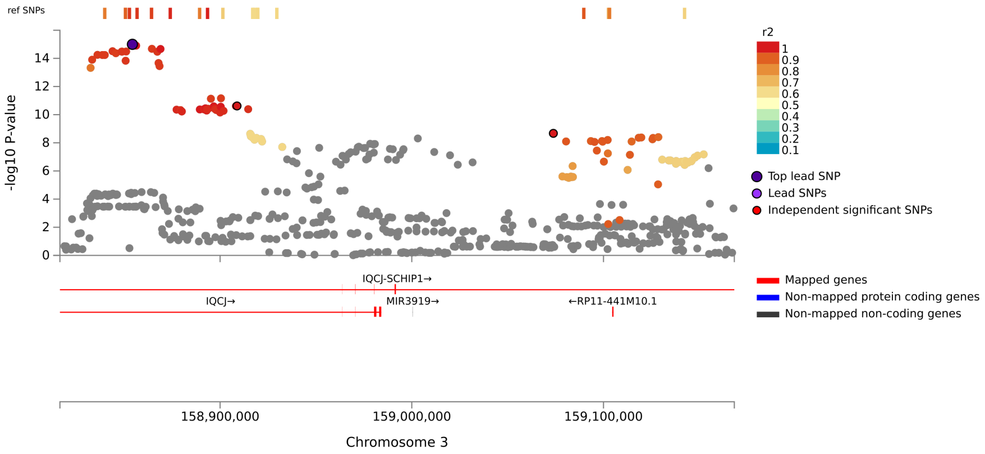

Locus 3, TBC1D14, Splenium Mean Thickness, rs11938701

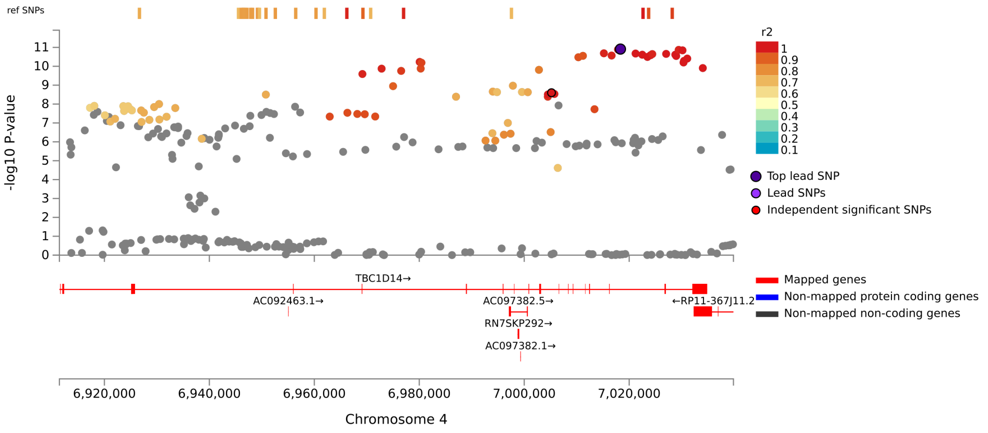


Locus 4, FIP1L1, Splenium Mean Thickness, rs6835429

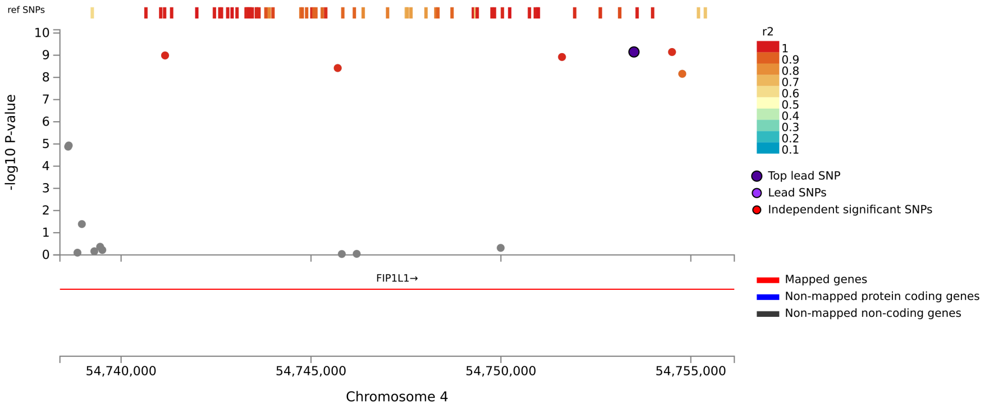

Locus 5, CTB-118N6.2, Splenium Mean Thickness, rs10074788

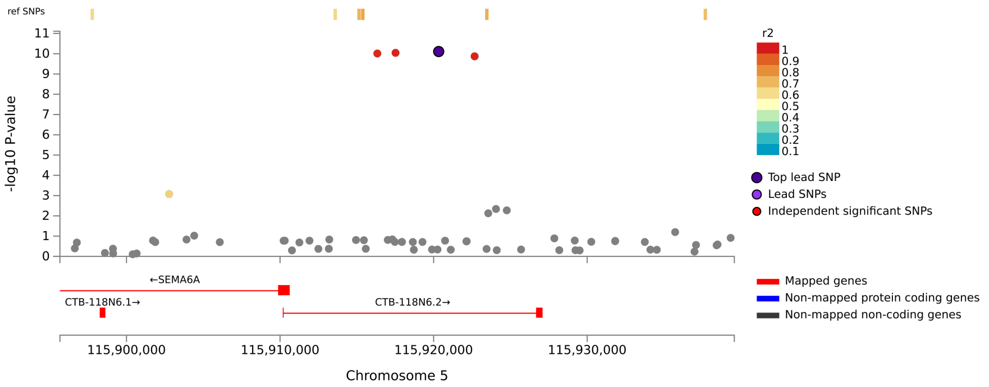


Locus 6, ANKRD19P, Splenium Mean Thickness, rs10992472

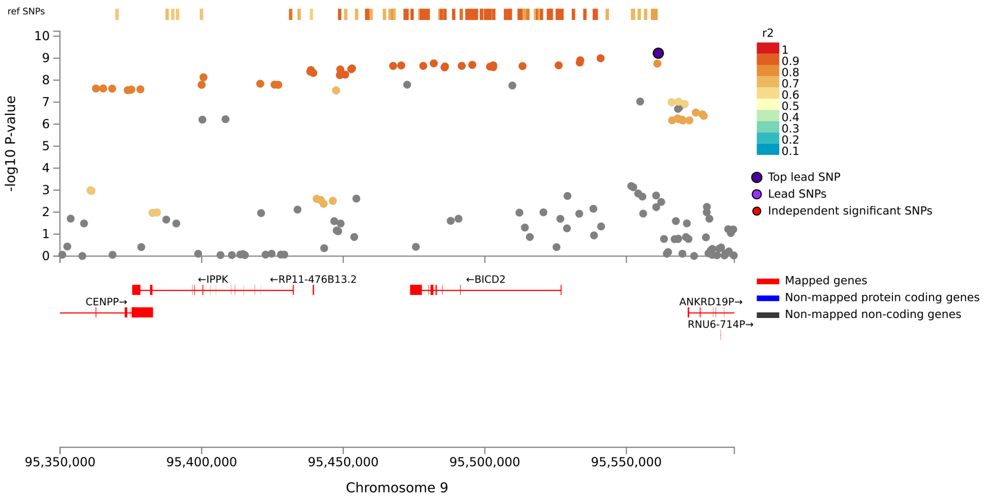

Locus 7, FAM107B, Splenium Mean Thickness, rs10906725

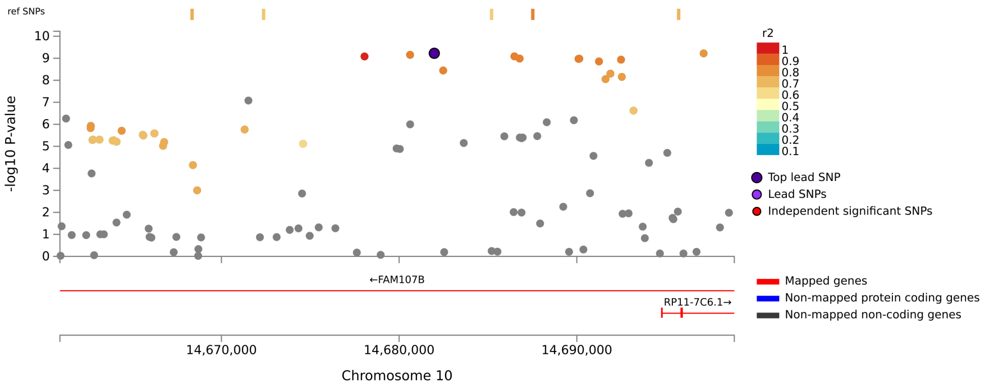


Locus 8, RP11-12J10.3:FAM53B, Splenium Mean Thickness, rs11245

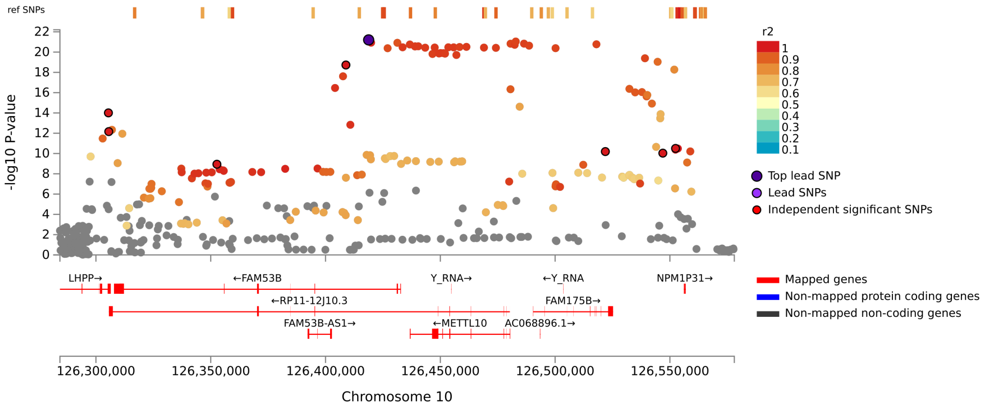

Locus 9, RRAS2, Splenium Mean Thickness, rs35247669

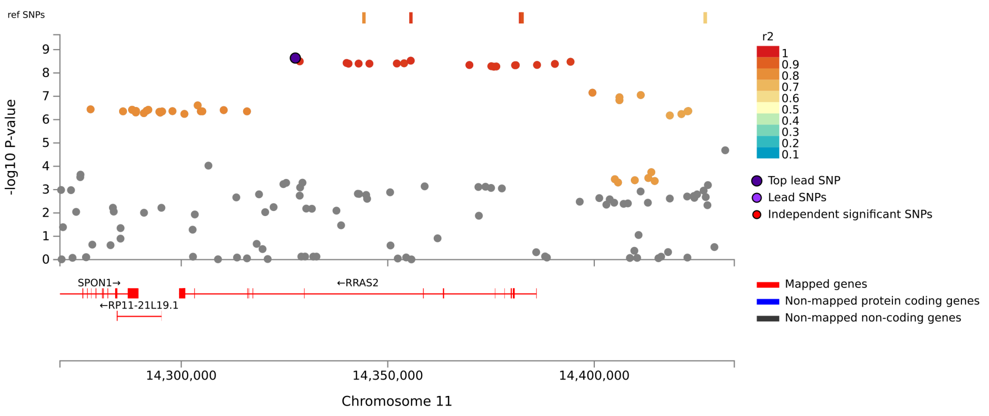
